# Passenger Hotspot Mutations in Cancer

**DOI:** 10.1101/675801

**Authors:** Julian M. Hess, Andre Bernards, Jaegil Kim, Mendy Miller, Amaro Taylor-Weiner, Nicholas J. Haradhvala, Michael S. Lawrence, Gad Getz

## Abstract

Hotspots, or mutations that recur at the same genomic site across multiple tumors, have been conventionally interpreted as strong universal evidence of somatic positive selection, unequivocally pinpointing genes driving tumorigenesis. Here, we demonstrate that this convention is falsely premised on an inaccurate statistical model of background mutagenesis. Many hotspots are in fact passenger events, recurring at sites that are simply inherently more mutable rather than under positive selection, which current background models do not account for. We thus detail a log-normal-Poisson (LNP) background model that accounts for variation in site-specific mutability in a manner consistent with models of mutagenesis, use this model to show that the tendency to generate passenger hotspots pervades all common mutational processes, and apply it to a ~10, 000 patient cohort from The Cancer Genome Atlas to nominate driver hotspots with far fewer false positives compared to conventional methods. As the biomedical community faces critical decisions in prioritizing putative driver mutations for deep experimental characterization to assess therapeutic potential, we offer our findings as a guide to avoid wasting valuable scientific resources on passenger hotspots.

## 1. Introduction

The genome of a cell lineage continually accrues mutations over time. The vast majority of mutations are either selectively neutral “passengers” that leave the lineage phenotypically unaltered, or else selectively negative events that result in slower growth or cell death. However, rare selectively positive mutations (“drivers”) that increase a cell’s proliferative fitness occasionally occur. Such a cell may go on to acquire additional driver events that enable it to outcompete its neighbors, eventually multiplying into a tumor (Cairns, 1975; Stratton et al., 2009).

Selectively positive mutations accumulate in tumor suppressors and oncogenes. These cancer driver genes are recurrently mutated across patients, and can be identified on the basis of having mutational densities significantly above the background passenger density. This requires accurate estimation of the mutational background, a task complicated by its considerable heterogeneity (Hodgkinson and Eyre-Walker, 2011). Some genomic elements are recurrently mutated due not to positive selection but simply to their higher mutability (e.g., late-replicating regions) (Hodgkinson and Eyre-Walker, 2011; Stamatoyannopoulos et al., 2009; Pleasance et al., 2010). Over the past decade, the field has developed increasingly sophisticated statistical models (Lawrence et al., 2013; Weghorn and Sunyaev, 2017; Martincorena et al., 2017; Lawrence et al., 2014; Dees et al., 2012; Getz et al., 2007) to infer and account for the heterogeneous mutational background. This has increased power and specificity to distinguish true drivers (with an excess of positively selected mutations) from false positives (with increased mutation density due to high background mutability alone).

Some of the best-known oncogenes are recurrently mutated at exactly the same amino acid position in many patients: for instance, the V600E mutation in *BRAF*, and mutations at G12, G13, or Q61 in *RAS* genes. These hotspot mutations have been experimentally validated as oncogenic (Davies et al., 2002; Field and Spandidos, 1990). Inspired by these examples, many methods for detecting driver events use exact positional recurrence as a signal of positive selection. This requires a model of background mutability at the site-specific level. The prevailing assumption has been that all equivalent base-pairs within a particular gene (e.g., all sites with the same *k*-mer sequence context) have the same background probability of being mutated. Thus, it is unlikely to observe by chance many patients sharing mutations at a particular base-pair in a gene while the other equivalent base-pairs of the gene remain unmutated. This leads to the conventional assumption that sites with multiple mutations across cancer (i.e., mutational hotspots) *must* reflect true driver events and that there are no “passenger hotspots.”

Here, we present evidence to the contrary. In the same way that certain regions of the genome are more highly mutated than others, and certain fragile sites of the genome are more prone to breakage (e.g., in genes such as *WWOX*, *FHIT*, and others (Schrock and Huebner, 2015; Mitsui and Tsuji, 2012)), it appears that certain individual base-pairs in the genome are more highly mutable, simply because they are more vulnerable to damage and/or more refractory to repair. We first show that the majority of recurrently mutated sites are under no positive selection by examining the distribution of their protein-coding effects (i.e., nonsense, missense, synonymous). We then demonstrate pervasive heterogeneity in the background mutability of individual base-pairs in the exome beyond what can be explained by known genomic features, and introduce a model that accounts for this base-level heterogeneity, allowing true cancer driver hotspots to rise to the top of the list above the numerous passenger hotspots that form at base-pairs with high intrinsic mutability. We use this new model to detect novel driver hotspots and discuss their potential biological functions. Finally, we quantify the extent of base-level heterogeneity of various mutational signatures, such as those associated with APOBEC enzyme hyperactivity, loss of mismatch repair or polymerase proofreading, or mutagens such as ultraviolet light or tobacco smoke. Surprisingly, we find that most mutational processes show a similar amount of heterogeneity in base-pair mutability This may reflect intrinsic properties of the genome, or ubiquitous repair mechanisms, rather than shared properties of mutagens.

## 2. Results

To identify significantly mutated hotspots, we must first be able to statistically estimate position-specific mutation frequencies. This requires a large cohort of very high-quality somatic mutation calls, since small cohorts are underpowered to accurately estimate base-level mutation frequencies, and poor-quality mutation calls typically contain recurrent sequencing artifacts or germline polymorphisms that can severely distort base-level analyses. Here, we examined a cohort of 9,023 quality-controlled whole-exome-sequenced tumor-normal pairs spanning 32 tumor types, generated by the The Cancer Genome Atlas (TCGA) MC3 mutation-calling initiative (Ellrott et al., 2018) (STAR Methods), a dataset containing a total of 2,288,080 somatic single-nucleotide variations (sSNVs).

### 2.1. Discovering significant hotspots based on a statistical model of the site-specific background mutational frequency

Candidate driver hotspots are discovered by identifying sites with a significant excess of mutations beyond the background frequency Because we do not know the true background mutation frequency of each genomic position *a priori*, we must approximate it with a statistical model. Regardless of the model, a common approach to find significantly mutated bases (i.e., “hotspots”) is to compare the observed number of mutations across a cohort at a given site to the distribution of the expected number of mutations predicted by the background model, generating a *p*-value for that site. Sites passing multiple-hypothesis correction (e.g., false discovery rate [FDR] *q*-value ≤ 0.1) are then considered significant hotspots. Obviously, what gets called statistically significant is only biologically meaningful if the underlying statistical approximation of the true background mutation frequency is accurate. Although we cannot directly evaluate the underlying model’s overall accuracy since we lack ground truth background mutation frequencies for every genomic position, we can assess a given model’s specificity via orthogonal criteria for whether mutations deemed significant are indeed under positive selection.

One orthogonal criterion we can use to assess whether a mutation is under positive selection is the distribution of variant protein-coding effects compared to expectation. Variant effects of mutations under no selective pressure will be randomly distributed according to the codon structure and signatures of mutational processes operating throughout the exome. In contrast, positively selected mutations are mostly nonsynonymous or splice altering, since essentially all coding driver mutations alter their corresponding protein. Therefore, an effective proxy for the specificity of a given algorithm is the degree to which the candidate driver hotspots it finds are enriched for nonsynonymous mutations beyond the expected baseline.

We calculate this baseline by using the overall exonic substitution frequencies for each trinucleotide context to generate the expected background frequencies of each coding effect — synonymous, missense, nonsense — in that context (STAR Methods). For example, we expect background C→T mutations within the sequence context T<u>C</u>G to be 27% synonymous, 69% missense, and 4% nonsense (see **Figure S1** for the full distribution). By adding the contributions from all 96 trinucleotide contexts, weighted by their relative frequencies, we find an overall expected baseline distribution of 28% synonymous, 67% missense, and 5% nonsense mutations. Note that throughout our entire analysis, we avoid splice-altering mutations by excluding genomic positions and mutations within 5 basepairs of a splice site, since synonymous events at those positions frequently disrupt splicing (Supek et al., 2014).

### 2.2. Novel hotspots found significant by conventional algorithms are mostly neutral

A simple and, to date, common background model employs the conventional assumption used in many studies up until recently (Van den Eynden et al., 2015; Lohr et al., 2012; Lawrence et al., 2014; Chang et al., 2016; Baeissa et al., 2017; Araya et al., 2016; Miller et al., 2015) that all sites within the same *k*-mer context in a given gene are equally mutable. Our version of this model, which we will refer to as “Uniform-within-gene,” assumes that the background mutation frequency at a given position is proportional to the average exome-wide mutation frequency for the base substitution/trinucleotide combination at that position, weighted by a gene-specific mutability factor that captures the overall number of mutations in the gene (STAR Methods). Expected counts are Poisson-distributed around this background frequency. Applying these assumptions yields a long list of 1,677 significant hotspots within 1,218 genes across our pan-cancer cohort. Although many of these significant hotspots occur in known cancer genes (KCGs) as defined by the Cancer Gene Census (CGC) (v85) (Futreal et al., 2004) (STAR Methods) — 92/218 KCGs contain significant hotspots (**Figure 1a**) and recapitulate known driver events (e.g., *BRAF* V600E) — 1,250 significant hotspots occur in 1,116 genes with uncertain oncogenic roles (non-KCGs) (**Figure 1a**). All of these hotspots have 3 or more patients with the exact same base-pair substitution.

**Figure 1:**
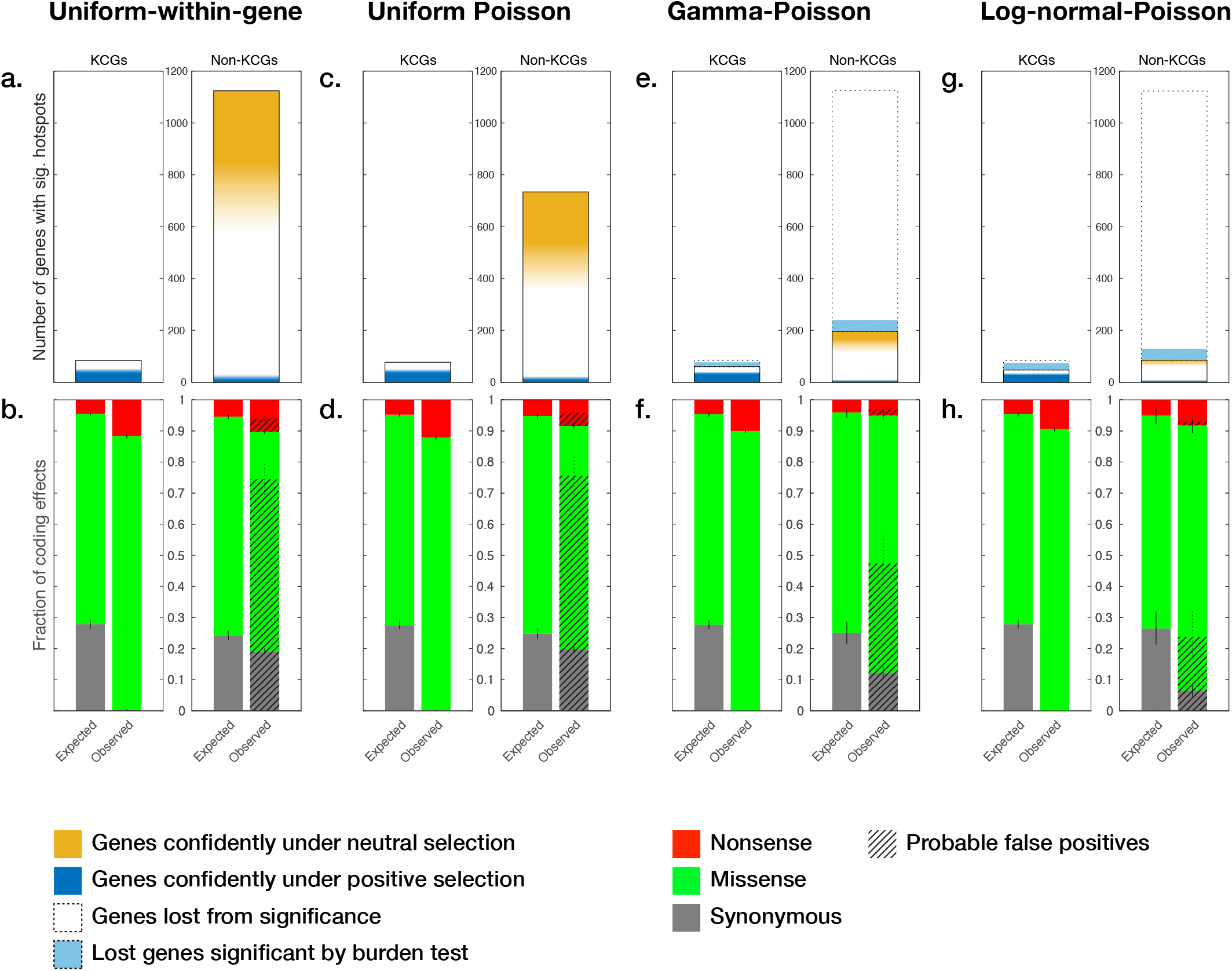
Comparison of number of genes containing hotspots and protein-coding effects of hotspots found significant by the four statistical models. **a., c., e., g**. Number of genes containing significant hotspots (*q* ≤ 0.1) according to each of the four statistical models, segregated by whether the gene is a Known Cancer Gene (KCG) (i.e., in the Cancer Gene Census). Genes under neutral selection are conservatively defined to have ≥80% probability of their *dN/dS* falling between 0.8 and 1.1; orange scale fades to white when this probability falls below 0.5. Genes under positive selection are defined to have ≥97.5% probability *dN/dS* ≥ 1.2; blue scale fades to white when probability falls below 0.7. Genes are denoted as lost from significance relative to the Uniform-within-gene method. **b., d., f., h**. Expected and observed distributions of protein coding effects of hotspot mutations significant by each model (*q* ≤ 0.1), segregated by Known Cancer Gene status. We estimate the overall fraction of false positive passenger mutations (hatched bars) by assuming all significant synonymous hotspots are false positives. Thus, the proportion of observed nonsynonymous mutations concordant with the expected ratio between synonymous and nonsynonymous mutations will also be passengers. Lines between stacked bars denote 95% confidence intervals.

Significant hotspots in KCGs are nearly devoid of synonymous mutations (0.4% of mutations at hotspots vs. the expected 27.9%), providing strong orthogonal evidence that these mutations are indeed under positive selection. If the 1,250 putative driver hotspots in non-KCGs were also drivers, one would expect that they too would be highly depleted of synonymous mutations (STAR Methods). Instead, however, we find that the distribution of their coding effects is very close to what we expect by chance (observed silent: 19.1% vs. expected 24.3%, **Figure 1b**), indicating that the majority of these hotspots are under little-to-no selective pressure and are therefore most likely neutral passenger events, as elaborated below.

If we conservatively assume that mutational patterns in KCGs reflect mutational patterns of driver genes in general, then the overwhelming majority of all silent hotspots are likely to be passenger events, given their extreme paucity in KCGs. We would therefore expect a proportion of observed nonsynonymous mutations concordant with the expected ratio between synonymous and nonsynonymous mutations to also be passenger events. For example, if the expected ratio of synonymous to nonsynonymous mutations were 1:3, and 10% of putative significant hotspots were synonymous, then we would expect an additional 30% of the putative non-synonymous hotspots to also be passengers. By this reasoning, at least 78.8% of putative nonsilent driver hotspots in non-KCGs according to the conventional Uniform-within-gene model are also false positives.

In addition to finding that these hotspot mutations at the base-pair level are neutral, we also noted that a large fraction of the genes containing these hotspots are themselves neutral (i.e., they show the expected ratio of nonsynonymous:synonymous mutations). To determine this, we assessed whether a gene was under neutral selection via the molecular evolution criterion of the ratio of a gene’s nonsynonymous:synonymous somatic mutation densities (*dN/dS*) (Kimura, 1977; Martincorena et al., 2017; Nei and Gojobori, 1986). After normalizing for signature heterogeneity and genes’ codon structures (Greenman et al., 2006) (STAR Methods), we expect parity between these densities (i.e., *dN/dS* ≈ 1) in genes under neutral selection. Computing *dN/dS* for each gene can only yield a confident estimate in genes with sufficient numbers of mutations (i.e., genes with a relatively high background mutation frequency), allowing us to conservatively assess whether they were under neutral selection. We identified 404 (of the ~20,000) genes confidently under neutral selection (≥ 80% probability that the gene’s *dN/dS* is between 0.8 and 1.1). 23.8% of non-KCGs that contained significant hotspots were either within these 404 genes (*n* = 101) or only contained silent hotspots (*n* = 165) (**Figure 1a; Table S1**), further confirming the poor specificity of the conventional Uniform-within-gene model. Overall, we conclude that analysis methods based on the conventional assumption, that equivalent base-pairs (i.e., same *k*-mer context/gene) have the same background mutability, produce lists of candidate hotspots with many false positives.

Thus, the apparent significant recurrence of mutations even at sites/genes under neutral selection is due to an incorrect background model that fails to account for the underlying heterogeneity in the background base-wise mutability. Some sites may simply be more intrinsically mutable than others, rendering them unusually prone to recurrent mutations even in the absence of selective pressure, thus giving rise to passenger hotspots. We therefore need to model mutational recurrences with a site-specific background model.

### 2.3. Currently known covariates cannot account for all base-wise mutational heterogeneity

It is possible that the unaccounted variability might be completely explained by previously reported covariates that affect mutation frequencies on both coarse and fine scales. Covariates such as replication timing (Stamatoyannopoulos et al., 2009), chromatin state (Polak et al., 2015), and gene expression levels (Pleasance et al., 2010) have been reported to influence background mutability on a broad scale (~100kbp–1 Mbp). More recent studies have discovered that other covariates like nucleosome positions or transcription factor binding activity influence mutability on a much smaller scale (10s of basepairs) (Poulos et al., 2016; Sabarinathan et al., 2016; Mao et al., 2018). We tested a fixed regression model in which the base-wise mutation frequency is entirely determined by these covariates (STAR Methods), which we refer to as the “Uniform Poisson” model. However, we find that the covariates alone are incapable of explaining all of the variability of base-wise mutability, as the distribution of protein-coding effects of hotspots in non-KCGs found to be significant by this method is essentially identical to that of the Uniform-within-gene model (observed silent fraction of 19.8% vs. expected 24.8%; **Figure 1d**). Moreover, 26.1% (*n*(*dN/dS* ≈ 1) = 71, *n*(only silent hotspots) = 120) of the 732 non-KCGs containing significant hotspots are confidently under neutral selection (**Figure 1c**). Although additional yet-undiscovered covariates may be able to better explain this variability in the future, we currently are forced to update the model to explicitly allow for variability in base-wise mutation frequencies beyond what can be fully modelled with currently known covariates.

### 2.4. Introducing uncertainty in site-specific mutability improves specificity of finding driver hotspots with minimal loss of sensitivity

In contrast to the Uniform Poisson model, where a site’s mutation frequency is a constant determined unambiguously by its covariates, another way to account for the base-wise variability is by allowing the mutation frequency at each site to be drawn from a probability distribution reflecting the uncertainty in the background mutability of equivalent base-pairs (e.g., same *k*-mer context or mutational process). The choice and parameterization of the underlying distribution can make a large difference in model performance. Recent methods have employed a gamma-Poisson model, both at the gene level (Weghorn and Sunyaev, 2017; Martincorena et al., 2017; Imielinski et al., 2017) and on the base-level (Smith et al., 2016), in which mutation counts are still Poisson-distributed, but the Poisson rates vary according to a fitted gamma distribution, adding an additional parameter to represent the overdispersion. There is no inherent biological rationale for using the gamma distribution to represent the uncertainty of the Poisson rates — it is merely mathematically convenient, as there is an easy closed-form expression for a Poisson distribution whose rate is gamma-distributed: the negative binomial (also known as the gamma-Poisson) distribution. We found that applying the Gamma-Poisson regression model (STAR Methods) indeed explained a large amount of the additional variability, but it still did not capture all of it; while the set of significant hotspots in non-KCGs is depleted in synonymous events (observed 12.3% vs. expected 25.1%, **Figure 1f**), we still see that 19% (36/194) of non-KCGs containing significant hotspots are neutral (**Figure 1g**), suggesting that all the variability is not yet accounted for in this model and there is additional specificity to be gained.

Next, we tried to improve on the Gamma-Poisson model by replacing the gamma distribution with a log-normal distribution (STAR Methods). Unlike the gamma distribution, arbitrarily chosen for its mathematical convenience of its conjugacy to the Poisson distribution, the log-normal distribution is based on the idea that mutant base-pairs do not instantaneously arise but are the net result of many independent consecutive events (e.g., damage and repair processes), each with an independent probability of occurring. For instance, a mutagen has a certain probability of initially damaging a nucleotide, which in turn has a certain probability of being missed by all repair mechanisms before S-phase. Should all of these events occur, the DNA polymerase must also, with a certain probability, fail to recognize the lesion and incorporate the wrong complementary base; if this mutated base survives another cell cycle without being recognized, it becomes a mutated base-pair in the genome. By the geometric central limit theorem, this product of probabilities approaches a log-normal distribution (Sutton, 1997).

Applying this log-normal-Poisson (LNP) model to our cohort identified a list of candidate driver hotspots in non-KCGs that was the most depleted in synonymous events relative to expectation (observed 6.6% vs. expected 26.5%, **Figure 1h**), and the fraction of non-KCGs under neutral selection with significant hotspots was also the lowest compared to the other 3 models (8.4% (7/83) of genes; **Figure 1g**). These results suggest that the LNP model has the highest specificity among the 4 tested models.

Although this new statistical model increases specificity of finding true oncogenic hotspots, this potential advantage may come at the expense of decreased sensitivity. Indeed, 33 KCGs containing significant hotspots according to the Uniform Poisson model are lost from significance by the LNP model; 18 KCGs are lost from significance by the Gamma-Poisson model (**Figure 1e** and **Figure 1g**). Although both the Gamma-Poisson and LNP models find fewer KCGs containing hotspots than both conventional models, not all cancer genes driven by point mutations must have strong mutational hotspots. For example, many tumor suppressors can be inactivated via truncating mutations anywhere in their ORF; thus, while the gene as a whole is recurrently mutated, its mutations do not need to recur at the same specific genomic position to incur the same functional impact. A position with many truncating mutations would not have any additional fitness advantage over any other position with fewer truncating mutations, since any truncating mutations far enough upstream from the C terminus will either induce nonsense-mediated degradation of the mRNA or produce a non-functional partial protein product. Thus, recurrent mutations in cancer genes not driven by hotspots should not be considered false negatives for an algorithm that solely evaluates mutations at the single-site level.

Although such genes may not be driven by hotspots, they will still display an overall excess of nonsilent mutations and should therefore be significant by a gene-level burden test that assesses whether the gene as a whole has an excess of nonsilent mutations. Out of the 33 genes lost from significance under the LNP model, 26 are still significant by a conservative burden test (Prob[*dN/dS* ≥ 1.2] ≥ 0.975). The additional 7 genes have too few mutations to confidently establish an excess of nonsilent events by our conservative criterion but are all deemed significant by more sophisticated methods (e.g., MutSigCV (Lawrence et al., 2013), which incorporates genomic covariates to estimate gene-level background mutation frequencies). Similarly, the 18 genes lost by the Gamma-Poisson model are also detected by gene burden tests. Thus, both the LNP and Gamma-Poisson models retain sensitivity if we specifically differentiate cancer genes driven by hotspots from genes driven by an overall excess of mutations.

### 2.5. Comparative analyses confirm improved performance of the LNP model

To more rigorously assess the sensitivity and specificity tradeoffs of the different methods, we used Receiver Operating Characteristic (ROC) curve analysis employing as ground truth sets: (i) a false-positive set of 404 genes confidently under neutral selection, as defined before (Prob[0.8 ≤ *dN/dS* ≤ 1.1] ≥ 80%); and (ii) a true-positive set of 44 genes, defined as KCGs with a high concentration of nonsilent mutations at recurrently mutated positions (as opposed to an excess of nonsilent mutations distributed across the ORF). We identified these 44 true positive genes by requiring that the *dN/dS* of the genes dropped by more than 5% when we removed all sites near significance (*q* ≤ 0.25) by the least conservative Uniform-within-gene model (**Table S1**). We then plotted ROC curves based on these conservative ground truth sets for the four methods, using each method’s minimum q-value across all sites in the gene as the discrimination threshold (**Figure 2a, Figure S2a**). We observed that the LNP model had the highest area under the curve (AUC), followed by the Gamma-Poisson, Uniform Poisson, and Uniform-within-gene methods. On each curve, we denote the positions on the ROC curves corresponding to the standard significance threshold of *q* ≤ 0.1. At *q* ≤ 0.1, the LNP model identified 494 significant hotspots in 134 genes. Out of these 134 genes, 40 belong to either the positive (39 genes containing 169 hotspots) or negative (1 gene/1 hotspot) truth sets. The 1 false positive gene out of the 40 yields a 3% FDR, corresponding to a sensitivity of 89% (CI_95%_[75%, 95%]) with a false positive rate (FPR) of 0.25% (CI_95%_[0.006%, 0.91%]). At the same threshold, the Gamma-Poisson model achieved an insignificantly higher sensitivity of 95% (CI_95%_[85%, 99%]) but a 12-fold higher FPR of 3.5% (CI_95%_[1.91%, 5.44%]), corresponding to an FDR of ~25% (14 false positive genes out of 56). The specificity losses for the conventional models are even higher, with FPRs exceeding 15% and FDRs exceeding 60%. This ROC analysis suggests that the LNP model performs the best among the 4 models, having the highest specificity without a significant loss in sensitivity.

**Figure 2:**
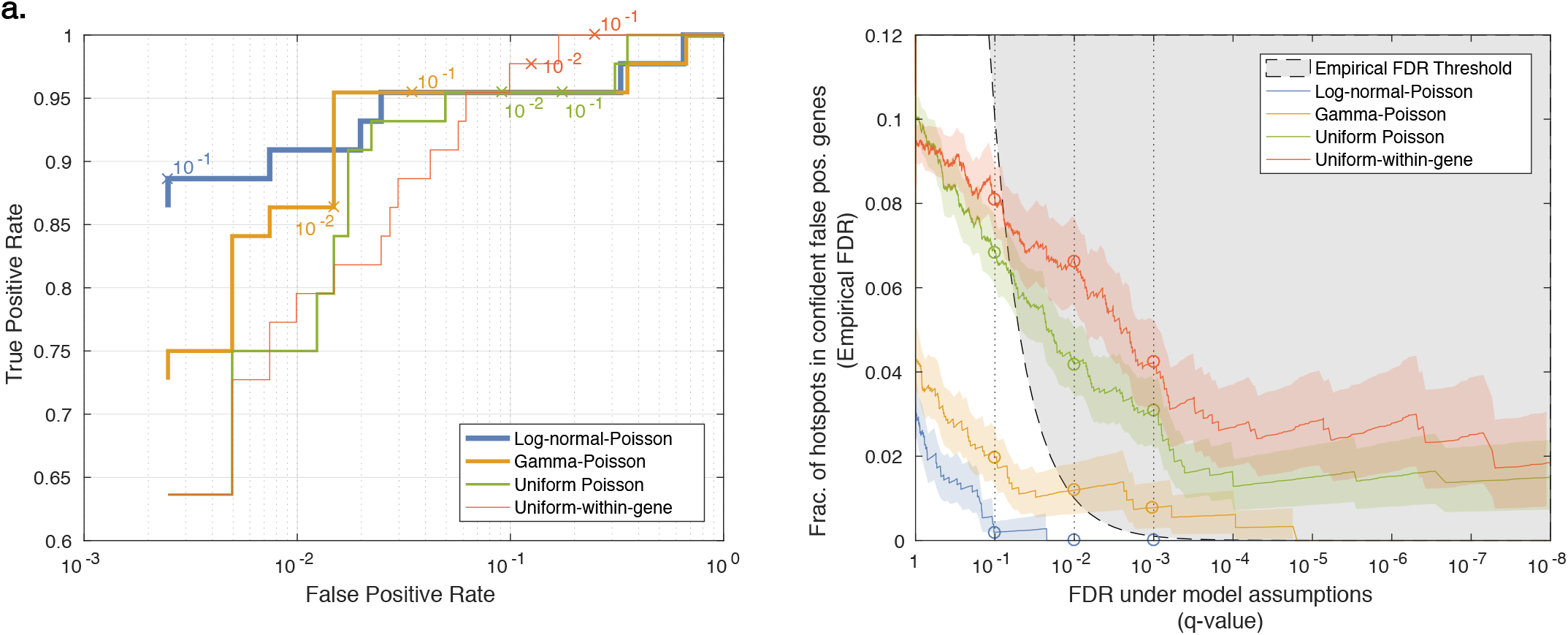
Performance of the four methods quantified by ROC and FDR analysis. **a.** Receiver Operating Characteristic (ROC) curves for each method evaluated at the gene level. Truth set used for estimating false positive rate comprises genes confidently under neutral selection; truth set used for estimating true positive rate comprises known cancer genes with a high proportion of mutations at recurrently mutated positions. A gene is considered a hit for the ROC analysis if it has at least one hotspot more significant than a specific q-value cutoff. Q-value cutoffs of 0.1 and 0.01 are marked on each curve. **b.** Fraction of loci falling in false positive truth set genes (empirical FDR) as a function of q-value. Gray area indicates region for which the empirical FDR exceeds the *q*-value; methods whose curves lie in this region yield more false positives than expected by the q-value cutoff. Colored regions indicate 95% beta distribution confidence intervals on fractions. *q*-value thresholds of 0.1, 0.01, and 0.001 are shown as vertical dotted lines, with circles denoting where they intersect the curves.

Another way to quantify the inflation of significant results of different methods is to examine their quantile-quantile (QQ) plots. Since we expect that most sites in the genome do not harbor driver events, we expect their *p*-values to be uniformly distributed. Indeed, when comparing the QQ plots of the four different methods, we observe that the QQ plots of the conventional methods are inflated, demonstrating deviation from the uniform distribution towards more significant *p*-values in a large fraction of genomic loci (**Figure S2b**). The inflation of the models also affects the resulting *q*-values and produces lists of significant hotspots (and genes) that may contain more false positives than expected by the q-value cutoff. In the case of a well-calibrated model (and hence a well-behaved QQ plot), setting a specific *q*-value threshold (e.g., *q* ≤ 0.1) would result in a list of significant hits that contain (on average) at most the desired fraction (10%) of false positives. It is therefore important to test whether this is indeed the case.

We used our set of false positives to measure the empirical false discovery rate. Since this is a conservative list, we expect to have an even lower FDR than the chosen *q*-value cutoff. We compare the empirical FDR as a function of *q*-value among the 4 models (**Figure 2b**). The LNP model is the only model for which the empirical FDR did not exceed the desired FDR (**Table 1**). By contrast, even at the extreme q-value threshold of 10^-8^, approximately 5% of hotspots significant by the Uniform-within-gene or Uniform Poisson models are in false-positive genes. Thus, these data confirm that the LNP model has the highest specificity of the four tested models and is the only model in which the *q*-value cutoff properly bounds the false discovery rate.

**Table 1:** Fraction of hotspots in confident false positive genes (empirical FDR) at the indicated q-value cutoff. Number of hotspots in false positive genes and total number of significant hotspots at each q-value cutoff are indicated in parentheses.

| Method | $q \leq 0.1$ | $q \leq 0.01$ | $q \leq 0.001$ |
| --- | --- | --- | --- |
| Uniform-within-gene | 0.08 (136/1678) | 0.07 (61/920) | 0.04 (25/590) |
| Uniform Poisson | 0.07 (106/1547) | 0.04 (42/1006) | 0.03 (22/713) |
| Gamma-Poisson | 0.02 (14/710) | 0.01 (6/504) | 0.01 (3/387) |
| Log-normal-Poisson | 0.002 (1/511) | 0.0 (0/290) | 0.0 (0/191) |

### 2.6. The LNP model produced the most accurate estimates of neutral mutation frequency

In addition to evaluating model performance by examining the protein-coding effects of hotspots found significant by each model, we can also assess how well each model predicts the expected number of mutated patients at each genomic position. A well-calibrated model accurately infers the background mutation frequency at each position, so the expected mutation frequencies at positions under neutral selection (and thus mutated solely due to background processes) will be concordant with the observed frequencies. On the other hand, a poorly-calibrated model that inaccurately models the background frequency will predict mutational frequencies that significantly deviate from the observed frequencies.

To compare the accuracy of the different background models, we calculated the observed distribution of recurrent synonymously mutated sites (i.e., the fraction of sites with a specific number of mutations out of all sites that can harbor synonymous mutations) (**Figure 3a**). We then compared to the expected distributions predicted by each of the four models. While the conventional models underestimate the fraction of sites mutated in 3 or more patients — synonymous sites mutated in exactly 3 patients are 3.5 x more likely than expected by the Uniform-within-gene model; overall, synonymous mutations recurring in ≥3 patients are 12x more likely than expected by the model — both overdispersed models (Gamma-Poisson and LNP) are more accurate, with the LNP model most correctly recapitulating the observed distribution even for highly recurrent events, with synonymous mutations recurring in ≥3 patients 1.84x more likely than expected by the Gamma-Poisson model, but only 1.05x more likely by the LNP model.

**Figure 3:**
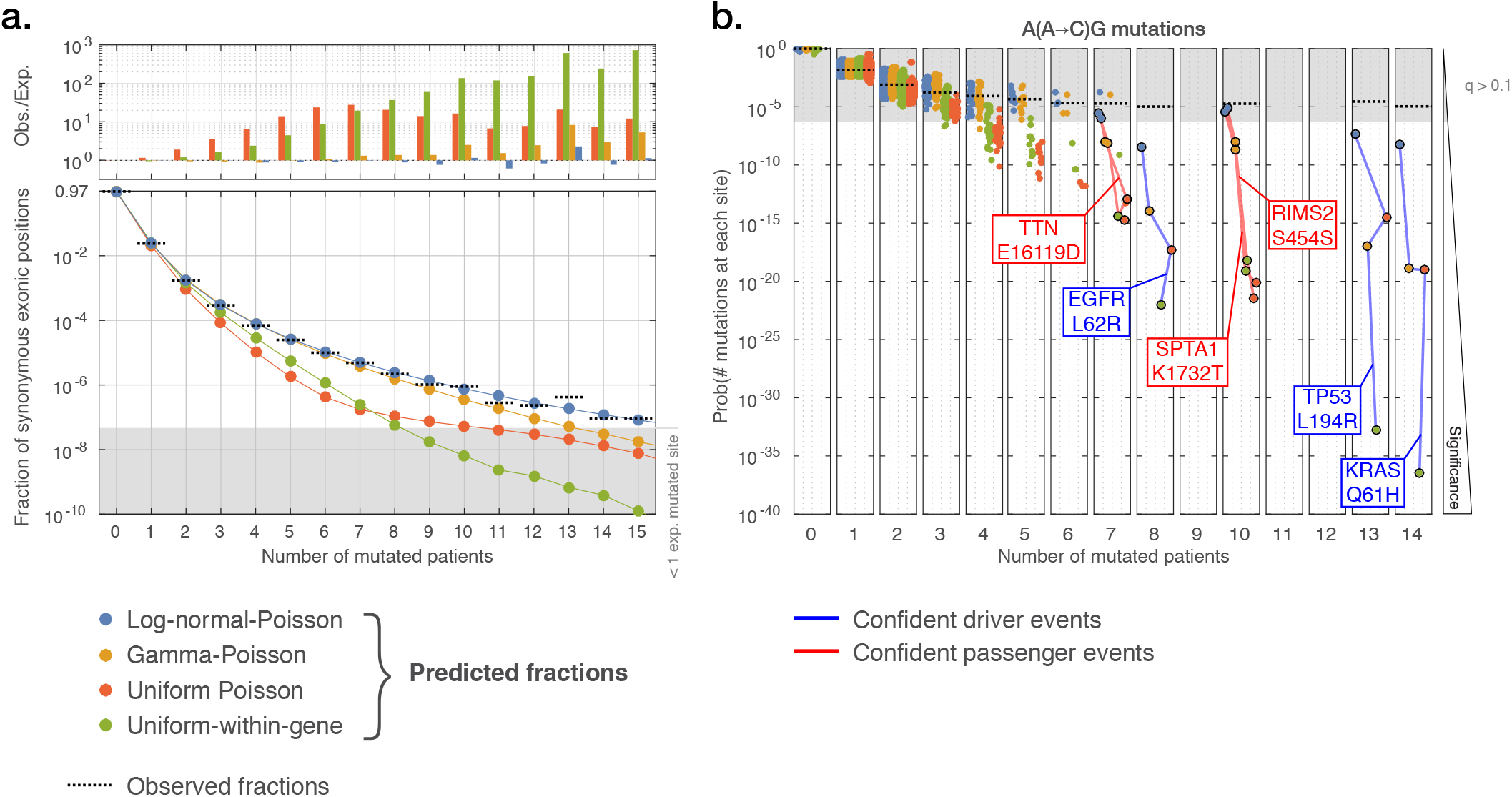
Predicted frequencies of recurrent mutations according to the four methods. **a.** Fraction of synonymous hotspot mutations observed in multiple patients (0 to 15; total cohort size 9,023). Colored lines represent the expected fraction of sites as predicted by each of the four models; dashed black lines represent the observed fractions. Ratio of observed:expected fractions for each model are plotted above each recurrence level. Log-normal-Poisson model best matches the observed fractions. Grey region indicates fraction corresponding to <1 base-pair of synonymous exonic territory (21.4 million possible base-pair substitutions that could yield a synonymous codon change). **b.** Probabilities of observing every mutation in sequence context A(A→C)G as predicted by the four models; each dot corresponds to an observed mutation. Lines connect the different models’ predictions for the same mutation; blue lines highlight confident driver mutations, while red lines highlight confident passenger mutations. Grey region indicates model predictions with q-value >0.1 (i.e., non-significant recurrence), indicating that while driver hotspot mutations are significant by all models, only the log-normal-Poisson model correctly infers passenger hotspot mutations as non-significant.

Although the two overdispersed models performed much better in predicting mutation frequencies than the non-overdispersed models, they are not identical. We illustrate a specific instance of this by looking at a sequence context containing recurrent mutations in likely passenger genes that only the LNP model avoids calling significant (Figure 3b). Three of the most recurrently mutated positions in the sequence context A(A→C)G occur in Spectrin alpha (*SPTA1;* 10 missense mutations: 6 in stomach adenocarcinoma and one each in bladder, cervical, colon, and lung squamous), Titin (*TTN*; 7 missense mutations: 3 in stomach; 2 in colorectal; 1 each in lung squamous and esophageal), and regulating synaptic exocytosis protein 2 (*RIMS2;* 10 synonymous mutations; 5 in stomach, 3 in colon, 1 each in esophageal and liver). All three of these mutations are found to be significant by the Gamma-Poisson model (*q* = 5 x 10^-4^, *q* = 0.002, *q* = 0.002, respectively) and by both conventional models (*q*-values < 10^-13^ for all genes and models) but not by the LNP model (*q* = 0.97, *q* = 0.72, and *q* = 1.0). As all of these genes are only expressed in specific tissue types (*SPTA1* in red blood cell progenitors, *TTN* in muscle cells, and *RIMS2* in neurons), it is highly likely that these represent passenger hotspots. Additional evidence that these genes are passengers is provided by the *dN/dS* analysis, which assigns the genes tight confidence intervals around 1 (*SPTA1:* mean *dN/dS* 1.05, CI_95%_[0.92,1.19]; *TTN*: mean 0.98, CI_95%_[0.94,1.02]; *RIMS2* mean 1.02, CI_95%_[0.86,1.22]), and by the fact that the hotspot in *RIMS2* is synonymous. By contrast, three of the other most recurrently mutated positions in the context fall in known drivers (*EGFR*, *TP53*, and *KRAS*, mutated in 8, 13, and 14 patients, respectively), and are significant by all four methods. Thus, taken together, these data suggest that the overdispersed LNP model most accurately predicts actual mutation frequencies and makes the fewest false-positive calls.

### 2.7. LNP model analysis reveals significant hotspots in potential novel oncogenes

The main scientific interest of any significance analysis method is to discover promising novel driver candidates in the resulting significant gene list. Since the LNP model performed well in excluding many more false-positive passenger mutations than the other 3 tested models, we can have more confidence that the genes in the resulting list of still-significant novel hotspots are true drivers. The LNP model yielded 494 significant hotspots (*q* ≤ 0.1) in 134 genes (49 KCGs, containing 405 hotspots, and 85 non-KCGs, containing 89 hotspots) (**Table S2**). The KCGs contain 29 of the conservative true positive genes (including *KRAS*, *BRAF*, and *PIK3CA*) and also 20 other genes not in the truth set but still with significant hotspots, including *PTEN, SMAD4*, and *CDKN2A*.

Since we use an FDR threshold of *q* ≤ 0.1, approximately 49 out of the 494 significant hotspots should be false positives. If we assume that none of the 405 hotspots in KCGs are false positives, then we expect that 40(= 89 – 49) out of the 89 hotspots in 85 non-KCGs are true positives.

Within the 85 non-KCGs, 26 have been previously experimentally implicated in cancer but are not yet well-known enough to be included in the CGC, or are in the CGC but not due to somatic point mutations (e.g., implicated by germline risk alleles or copy-number/structural alterations). Below, we discuss four of these genes, with evidence on other genes noted in **Table S3**.

One such gene is *MYC*, which is well-known as one of the most recurrently amplified oncogenes in many cancer types (Beroukhim et al., 2010). We find an S146L mutation in 8 patients (3 head-and-neck, 2 colon, and 1 each for cervical, lung adeno, and melanoma). Nearby amino acid residue K143 is an acetylation site (Zhang et al., 2005), and c-Myc acetylation suppresses its ubiquitination, preventing degradation (Patel et al., 2004). The S146L substitution may affect ubiquitination at K143, preventing c-Myc degradation, as suggested by phosphorylation at a different site in lymphoma (Bahram et al., 2000). Indeed, *MYC* S146L is mostly copy neutral relative to amplified wildtype *MYC* in tumors from the same tissues (absolute copy number ranksum *p*-value = 0.02, **Figure S3d**).

Another such gene is *ERCC2*, which encodes a DNA helicase essential for nucleotide excision repair (NER) (Fuss and Tainer, 2011). We identify an N238S mutation recurrent in 9 bladder patients. Germline SNPs in *ERCC2* are associated with xeroderma pigmentosum, which greatly elevates cancer risk (Lehmann et al., 2014). However, *ERCC2’s* status as a somatic driver had not been characterized until recently (Kim et al., 2016). Mutations across the helicase binding domain (including N238S) were reported to strongly correlate with activity of a distinct mutational process (COSMIC Signature 5), providing strong evidence that Signature 5 mutations are the result of unrepaired DNA damage due to *ERCC2* loss-of-function (Kim et al., 2016). While the selective advantage of *ERCC2* loss in particular is unknown, loss of certain repair mechanisms has been shown to allow cells to evade DNA damage-induced apoptotic responses or checkpoints that would slow cell proliferation (Calvo et al., 2013).

Two non-KCGs belong to members of the Ras superfamily, *RRAS2* and *RHOB*. RRAS2 shares the majority of its amino acid sequence with canonical Ras subfamily oncogenes *HRAS, KRAS*, and *NRAS* (**Figure S3a**). The Q72L mutation observed in 11 patients spans a wide variety of tumor types (head-and-neck, lung adeno/squamous, prostate, testicular, and endometrial) and is paralogous to the well-known Q61 hotspot in *H/K/NRAS*, which abrogates GTP hydrolysis, leaving the protein constitutively active (Marcus and Mattos, 2015). The same constitutive activation in *RRAS2* has been experimentally confirmed both in mechanism and oncogenic potency (Graham et al., 1994), suggesting that *RRAS2* oncogenic activity is analogous to that of the canonical Ras subfamily members.

*RHOB*, a member of the Rho GTPase subfamily, has two hotspots near significance: E172K (*q* = 0.12, mutated in 7 bladder cancers), and P75[ST] (mutated in 3 bladder cancers). Mutations of E172 have been shown to be destabilizing, significantly reducing protein half-life (Hurst et al., 2017). E172 lies in the PKN-binding domain (Maesaki et al., 1999), which is conserved across Rho GTPases *RHOA/B/C* (**Figure S3b**). RHOB’s binding and activation of PKN specifically induces degradation of growth factor receptor EGFR (Gampel et al., 1999). Together, these lines of evidence suggest that mutant RHOB may contribute to oncogenesis by allowing excess EGFR to accumulate, thereby promoting growth. *EGFR* is known to be focally amplified in bladder cancer. The effects of P75 mutation is less clear — it occurs towards the end of the switch II domain, which binds to GEFs to facilitate guanine nucleotide exchange (Reijnders et al., 2017). This domain is both highly conserved and has a high burden of mutations across the 4 Rho GTPase members (**Figure S3b**). The same residue is mutated in *RAC1* and *RHOA*.

Next, we turn our attention towards the 59 remaining genes that have not yet been experimentally implicated in cancer. Following the same logic as above, if we conservatively assume that none of the 26 genes that were implicated in cancer due to other alterations are false positives, then we would only expect 10(= 59 — 49) out of these 59 to be true cancer genes. Although many of these genes are poorly studied, we briefly discuss four genes of interest below.

SLC27A5 ligates acyl-CoA to very long chain fatty acids, which is required for their subsequent metabolism. Mutation T554I recurs in 13 melanoma patients, arising via two distinct point mutations: a single C→T nucleotide substitution inducing codon change ACC→ATC (3 patients), and a dinucleotide CC→TT substitution inducing codon change ACC→ATT (10 patients). The presence of two distinct genomic events causing the same amino acid change is strong evidence that the amino acid substitution is under positive selection. T554 occurs within a highly conserved motif (GDTFR-WKGENV) across the solute carrier 27 family (**Figure S3c**) and in many orthologs. This motif is suspected to comprise an acyl-CoA synthetase domain (Pei et al., 2004), important to the function of the protein; the exact function of these mutations, however, is unknown.

Another gene involved in lipid metabolism is *SPTLC3*, which is mutated in 12 melanomas and one endometrial cancer. *SPTLC3* is highly expressed in skin tissue at an order of magnitude higher than most other tissue types (GTEx Consortium et al., 2017), suggesting a functional role.

EEF1A1 is a subunit of the elongation factor-1 complex that affects cell proliferation by various mechanisms, including mRNA translation downstream of TGFβ-receptors (Lin et al., 2010), affecting epithelial-mesenchymal transition as part of the BAT complex (Hussey et al., 2011). The hotspot in T432I is located in a phosphorylation site (Eckhardt et al., 2007) and appears in 4 liver tumors, 1 head-and-neck, and 1 GBM tumor.

UPF2 targets transcripts with premature stop codons for nonsense-mediated decay (Lykke-Andersen et al., 2000), and its E1033D mutation occurs 8 times (3 endometrial, 3 stomach, and 2 colon cancers). Nonsense-mediated decay has a major effect on tumors with microsatellite instability (MSI) since many transcripts can have frameshift insertions or deletions (indels). Indeed, the 8 mutated tumors are enriched with indels (ranksum *p*-value = 2.3 × 10^-7^; **Figure S3e**) relative to other patients in the same tumor types as the mutants. Therefore, altering nonsense-mediated decay in these tumors suggests a potential functional role for these mutations.

Overall, we demonstrate that the LNP model produces a highly accurate list of driver hotspots that provide clear biological hypotheses, some of which have already been supported by experimental data. Future experiments will be needed to validate the functional role of the other hotspots.

### 2.8. Hotspot-generating mutational processes have similar base-wise heterogeneity despite having vastly different mutation frequencies

Our results provide evidence of pervasive variability in base-wise mutation frequency across cancer, irrespective of mutagen or underlying mutational process. Next, we tested whether different mutational processes have different levels of variability. Each mutational process has specificity for particular genomic contexts and features, defining a set of “bases-at-risk” for that process. We might expect differences in the degree to which different mutational processes diverge from uniformly targeting their bases-at-risk and thus different levels of variability. Because there is considerable heterogeneity in the overall mutation frequencies of different processes (ranging from ~10 muts/million bases-at-risk for CpG deamination to >5, 000 muts/million bases-at-risk for POLE hypermutation), we might predict a correspondingly wide range in the variability across bases-at-risk of different processes. Tellingly, many published analyses have been forced to exclude hypermutated tumors from significance analyses because they yield too many significantly mutated genes/hotspots, largely due to the inaccurate background models in wide use (Bailey et al., 2018; Martincorena et al., 2017). One might surmise from this that high-mutation-frequency processes are more prone to generating hotspots, reflecting greater variability in their base-wise mutation frequencies.

The LNP framework provides a natural way of quantifying both the mutation frequency and variability of different mutational processes. We fit the model to all bases-at-risk for each process; the log-normal parameters *e^μ^* and *e^σ^* are equal to the geometric mean and geometric standard deviation, respectively, of the base-wise mutation frequency. The geometric mean is simply equivalent to the median mutation frequency across all bases-at-risk. The geometric standard deviation *e^σ^* is a dimensionless scale factor indicating the average multiplicative distance from the mean; for example, *e^σ^* = 2 signifies that at 1 standard deviation above the mean, bases will be twice as mutable as the average, while at 1 standard deviation below the mean, bases will be half as mutable as the average. At its minimum value *e^σ^* = 1, there is no variability in the base-wise mutation frequency. Hence, the same value of *e^σ^* indicates an equivalent amount of base-wise variability, irrespective of mutation frequency. Our fully Bayesian model finds not only the optimal values of *e^μ^* and *e^σ^* but also the posterior distribution on the parameters, allowing us to quantify the uncertainty on their values. In general, mutational processes that generate few mutations will have high model uncertainty, while processes that generate many mutations will have lower uncertainty. The values and uncertainties of *e^σ^* of two processes can be used to test whether they indeed have different levels of variability.

We selected the following 8 mutational processes to examine by the LNP model because they are mostly of known etiologies and very specific to certain patients and sequence contexts, making assignments of patients to these signatures unambiguous: APOBEC (3A+3B, COSMIC Signatures 2 and 13), aging (spontaneous CpG deamination, COSMIC Sig. 1), esophageal (Dulak et al., 2013) (COSMIC Sig. 17), MSI (COSMIC Sig. 6), POLE (COSMIC Sig. 10), POLE+MSI (Haradhvala et al.,2018) (COSMIC Sig. 14), smoking (COSMIC Sig. 4), and UV (UV-A only, COSMIC Sig. 7). We used SignatureAnalyzer (Kim et al., 2016), which is based on a Bayesian implementation of non-negative matrix factorization (NMF), to infer the probabilities of each mutation being assigned to each process. We then identified 8 subcohorts of patients (each comprising between 44 and 4,739 patients) in which each of these processes dominated (≥ 75% assignment probability) at their relevant sequence contexts (e.g., C→T mutations at CpG sites for aging; T(C→A)T, T(C→T)G, and A(A→C)A contexts for POLE cohort; and C→T mutations at pyrimidine dimers for UV). The mutational spectra of these subcohorts is shown in **Figure S4**. For each of these “process-centric” subcohorts, we fit the LNP model to the relevant contexts.

We plotted the posterior distributions of *e^μ^* and *e^σ^* for each pentamer context belonging to the 8 mutational processes to test whether processes with higher mutation frequencies also have higher base-wise variability. To our surprise, our results show that despite extreme heterogeneity in overall mutation frequency (spanning 5 orders of magnitude, from ~0.1 mutation/million positions for esophageal to >5, 000 mutations/million positions for POLE), most mutational processes show a similar amount of nonzero base-wise variability (**Figure 4a**), with *e^σ^* roughly between 2 and 3.

**Figure 4:**
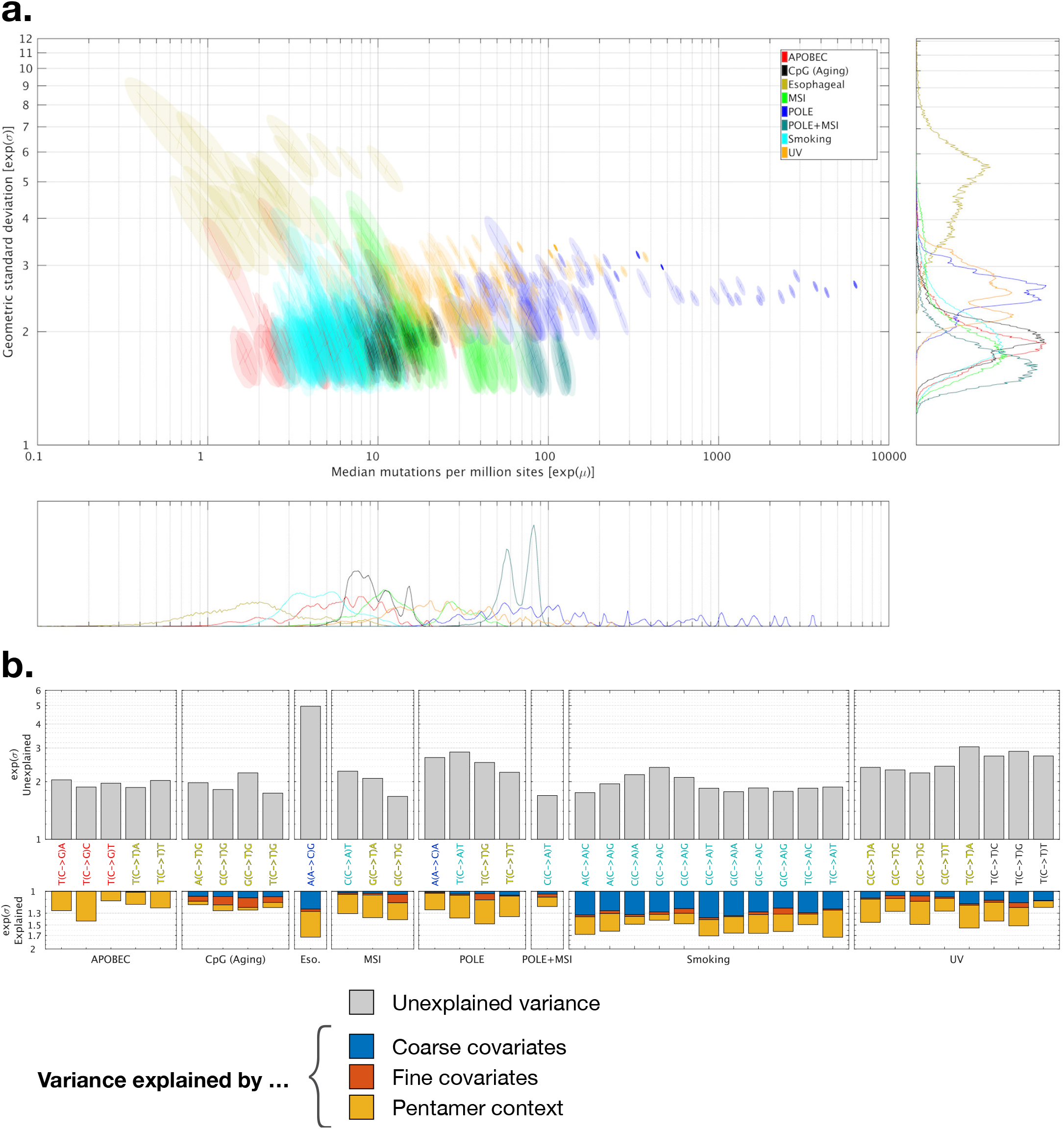
Comparison of the heterogeneity of base-wise mutability for different mutational processes as inferred by the log-normal-Poisson model. **a.** Log-normal-Poisson posterior distributions of base-wise mutation frequency (median mutations per million sites *e^μ^*) and mutations’ deviation from being uniformly Poisson distributed (geometric standard deviation *e^σ^*) for different mutational processes. Each colored area represents the posterior 95% confidence region for a pentamer context associated with a given mutational process (**Figure S4**). *e^σ^* = 1 corresponds to uniformly Poisson distributed mutations. Since *e^σ^* > 1 for all processes, we see that base-wise variability is universal and pervasive. Marginal distributions of *e^μ^* and *e^σ^* are shown below and to the right of the plot, respectively. Lines within each region are principal axes of the posterior density (i.e., eigenvectors of the covariance matrix of the estimated posterior density). **b.** Amount of base-wise mutation rate variability *e^σ^* explained by model covariates (colored), and remaining unexplained variance after all covariates have been incorporated (gray), for each relevant trinucleotide context in each mutational process.

Notably, the only exception was the esophageal mutational process, which showed the highest variability despite having one of the lowest mutation frequencies. This may indicate that additional yet-undiscovered factors may correspond to elevated mutability at specific bases-at-risk for this process. It was recently reported that the esophageal signature disproportionately mutates positions within CTCF binding sites (Katainen et al., 2015), possibly due to bound CTCF transcription factors occluding damaged bases from repair processes.

To test whether variability was independent of mutation rate even within individual processes, we partitioned the high mutation frequency processes between hypermutated samples (top decile of mutation frequency) and non-hypermutants, and compared their *e^σ^* values. We observed similar levels of base-wise variability between the two partitions (**Figure S5**), indicating that there is nothing unusual about the distribution of mutations in hypermutants, and nothing that should warrant their exclusion in significance analyses.

### 2.9. Explanatory power of genomic covariates differs among mutational processes

Although we previously showed in the Uniform Poisson regression model analysis that genomic covariates cannot fully explain all base-wise mutational variability, their explanatory power is nonzero. The LNP model provides a natural way to quantify the contribution of each covariate towards explaining this variability: the amount that *e^σ^* decreases as we incorporate additional covariates corresponds directly to the variance explained by the added covariates (STAR Methods). The value of *e^σ^* after all covariates have been incorporated is the unexplained variance, which would approach *e^σ^* = 1 if the covariates completely explained the base-wise variability, as there would be no extra variance beyond what the covariates predict.

We grouped the covariates by genomic scale: replication timing and expression (Lawrence et al., 2013) influence mutation frequencies on a coarse scale (~100 kbp–1 Mbp), while nucleosome positions and DNase hypersensitivity influence mutation frequencies on a fine scale (~10 bp). We also quantified the effect of accounting for pentamer context specificity on mutation frequency (i.e., considering the flanking ±2 upstream/downstream positions in addition to the immediate 5’/3’ positions).

In **Figure 4b**, we plot the amount of total variance explained by the aforementioned covariate sets (coarse, fine, and context) and the amount of remaining unexplained variance for each of the trinucleotide contexts associated with each mutational signature. From this analysis, the following universal patterns stand out: (i) unexplained variance is higher than explained variance; (ii) pentamer contexts are almost always important (with the notable exception of the aging signature); and (iii) there is no correlation between the amount of unexplained variance and the amount of explained variance, and processes with more total variance do not necessarily have more variance explained by covariates.

For certain processes, genomic covariates are completely non-explanatory. For example, although certain APOBEC trinucleotides display considerable variability amongst pentamer contexts (Chan et al., 2015), neither coarse nor fine covariates explain any additional variability. Conversely, coarse covariates — namely gene expression — explain a substantial amount of smoking mutational variability. This is expected since the C→A mutations comprising the smoking signature are caused by benzo[a]pyrene guanine adducts (Denissenko et al., 1996), which are often corrected by transcription-coupled repair (Fousteri and Mullenders, 2008) that occurs more frequently in highly expressed genes (Pleasance et al., 2010).

## 3. Discussion

Discovering cancer drivers from sequencing data requires overcoming the problem of detecting signal (mutational recurrence) above a background of non-random noise (variable intrinsic mutability). We detect drivers by formulating a statistical model of the background mutability and then looking for recurrence that significantly exceeds the expected background. This approach is only fruitful and reliable if the underlying background model is accurate. As the corpus of cancer genomes has exponentially grown over the last decade, we have become statistically powered to observe background mutational variability at increasingly fine genomic scales, which we must accordingly account for in our background models. Initially, around 2007, cohorts were so small (~10 patients) that we lacked the power to observe any variability at all. Thus, models at the time assumed that every genomic region was equally mutable, which led to the belief that all recurrent mutation beyond a uniform background level stemmed from positive selection (Getz et al., 2007; Sjöblom et al., 2006). As cohorts grew to ~1,000 patients, around 2013, we became powered to observe heterogeneity on the scale of a gene, and infer that, contrary to the convention that every recurrently mutated gene had to be a driver, the majority of recurrently mutated genes were actually passengers with high intrinsic mutability. We therefore had to update our background models accordingly to avoid driver lists swamped by false positives. However, cohorts were still too small to estimate mutability on levels smaller than a gene, so models did not account for it. This led to the assumption that any base-pair recurrently mutated beyond the overall background level of its gene had to be a driver (Van den Eynden et al., 2015; Lohr et al., 2012; Lawrence et al., 2014; Chang et al., 2016; Araya et al., 2016; Baeissa et al., 2017).

We are able to challenge this assumption in the current era of cohorts comprising ~10,000 patients, which powers us to estimate background mutability on the smallest possible genomic scale: that of the individual base-pair. In this paper, we have described a log-normal-Poisson (LNP) regression model that accurately models base-wise mutability, and in applying it to a ~10, 000 patient cohort, we demonstrate that base-wise mutational variability is so extreme that a large proportion of recurrently mutated base-pairs are in fact passengers. We show this by contrasting the LNP method to two toy models (Uniform-within-gene and Uniform Poisson) that purposefully do not account for base-wise variability. Unlike the majority of hotspots found significant by the LNP model, the majority of significant hotspots under the toy models are under no positive selection by orthogonal criteria.

Although our toy models are merely illustrative, many recently published algorithms intended for actual driver discovery do not account for base-wise variability in their background models. Using these inaccurate models, various group reported long lists of significant hotspots with many likely false positives; in one example, 1,202 hotspots were found, even when using a highly stringent FDR cutoff of *q* ≤ 0.01 (Chang et al., 2016). Since FDRs are only meaningful if their underlying statistics are well calibrated, rigorous validation of the statistical models should be performed whenever the significance levels are overly confident. Failure to do so has potentially severe consequences in situations like identifying genes for deep experimental follow-up or as novel drug targets. Since such gene identification often begins by selecting recurrent alleles observed in sequencing data, followed by expensive follow-up experiments, properly accounting for base-wise variability is essential when selecting these candidates to avoid wasting valuable scientific resources on passenger hotspots.

In addition to providing an improved model for assessing mutational significance, our new LNP model also sheds light on the fundamental nature of mutagenesis. It has long been known that background mutability correlates with coarse scale genomic features, and more recent studies have shown that specific fine scale genomic features undergo localized hypermutation. We show here that neither of these factors can explain the amount of observed base-wise variability, suggesting that there are yet-undiscovered properties of the genome that affect mutability. To demonstrate this, we examined the effect of adding XR-seq coverage — an extrinsic measurement of nucleotide excision repair (NER) activity, used to repair pyrimidine dimers resulting from UV damage — as a covariate in our model. We found that this covariate explained an additional 10% of variance in the UV process-centric subcohort, indicating that many passenger UV hotspots occur at loci predictably refractory to NER. Future studies may find more basic genomic features that could predict the XR-seq coverage or similar empirical measurements of mutability. These may include factors such as: (i) yet-unknown proteins bound to the DNA; (ii) tertiary structure of the DNA helix (Harteis and Schneider, 2014) (which can influence binding of such proteins); (iii) a combination of the two (a recent study (Mao et al., 2018) suggests that the binding of ETS transcription factors rotates adjacent pyrimidines into a more favorable conformation to form a cyclobutane dimer between adjacent C5-C6 bonds); (iv) local chromatin structure (which could affect gene expression or accessibility to mutagens or repair enzymes); or (v) sequence-specific polymerase error modes that our datasets are not yet powered to detect.

Accounting for all potential covariates, however, may never be able to fully explain the observed variability since the mutations we observe in a tumor genome are merely a static snapshot representing the product of many dynamically fluctuating mutational processes that have been active over the course of the tumor’s life history. These processes’ activity levels and bases-at-risk vary as a function of continuously changing factors like the tumor’s microenvironment, mutagen exposure, or epigenetic state, which are complicated to model with static covariates. We probabilistically represent the unexplained variability with a log-normal distribution because it represents the net product of consecutive molecular events, each with some unknown probability of occurring.

Regardless of the underlying causes of the dramatic heterogeneity we observe in base-wise mutability, its existence has implications in the fields of molecular evolution and population genetics. We find that base-wise heterogeneity is pervasive across all mutational processes, including methylated-CpG deamination, which is overwhelmingly responsible for *de novo* germline mutations (Ehrlich and Wang, 1981; Hodgkinson and Eyre-Walker, 2011). This suggests that the infinite sites model underpinning many population genetics assumptions may be incorrect — for instance, the probability of identical alleles in multiple unrelated individuals originating from different common ancestors may be much higher than a naïve coalescent theory would predict. In addition, large-scale genomic organization has been thought to reflect coarse variability in background mutability (Chuang and Li, 2004), wherein genes more tolerant of mutation are thought to reside in more highly mutable regions of the genome. Variability at the base-pair level may equivalently mold genomic architecture on fine scales. For example, it has long been speculated that the sequence composition of immunoglobulin variable chains is specifically biased to induce AID hypermutation hotspots within domains encoding antigen-binding sites (Jolly et al., 1996) in order to accelerate the process of antigen selection. Genome-wide selective pressure may be analogously guided or constrained due to variable site-specific mutability.

In conclusion, further growth of cancer sequencing datasets will allow us to survey the landscape of drivers with even greater precision, reveal the intricacies of mutational processes, and even elucidate how these mutational processes shape evolutionary selection. But as datasets continually grow, we must continually challenge the statistical assumptions we make in analyzing them. As datasets evolve, so must our statistical methods and the conclusions we draw from them.

## 4. Author Contributions

Conceptualization, J.M.H., M.S.L., G.G.; Methodology, J.M.H., G.G.; Software, J.M.H.; Formal Analysis and Investigation: J.M.H.; SignatureAnalyzer results, J.K.; Data Curation, J.M.H., M.S.L., A.B.; Visualization, J.M.H.; Writing: Original Draft, J.M.H., M.S.L., G.G.; Writing: Review and Editing, G.G., M.S.L., M.M., A.T.-W., N.J.H.; Supervision, G.G., M.S.L.; Funding Acquisition, G.G.

## 5. Acknowledgements

We thank Daniel Rosebrock and Dimitri Livitz for providing *MYC* absolute copy number calls, Yosef Maruvka for providing microsatellite indel calls with respect to *UPF1* mutant status, and Chip Stewart for helpful comments on the manuscript. J.M.H. was funded by … M.S.L. was partially funded by G.G. funds at Broad Institute and M.S.L. startup funds at Massachusetts General Hospital. G.G. was partially funded by Paul C. Zamecnik Chair in Oncology, MGH Cancer Center.

