## Supplementary Methods, Figures, and Tables for "Passenger Hotspot Mutations in Cancer"

#### STAR★Methods A: Mutation calling

We obtained the MC3 TCGA mutation calls from Synapse, a cohort comprising 10,510 unique patients (3,850,525 sSNVs called by two or more mutation callers) before filtering. MC3 is a curated set of high-quality mutation calls run through numerous QC filters with failing samples/mutations annotated in the mutation annotation file (MAF), allowing the end user to exclude them.

We excluded all patients failing MC3 sample-level filters (i) excessive cross-sample contamination (MAF filter `contest`); (ii) whole genome amplified libraries (MAF filters `wga_only`, `native_wga_mix`); and (iii) bad sequencing runs due to mid-run sequencer failure (MAF filter `badseq`). This left us with a cohort of 9,023 patients, from which we further excluded mutations failing the following MC3 site-level filters:

(i) Panel-of-normals (PoN): for each genomic position, the PoN encodes the distribution of alternate read fractions (AFs) across $\approx$ 8,000 TCGA normals. We removed candidate variant calls that occurred at sites recurrently harboring alternate reads across the PoN whose AFs were consistent with the candidate variant. For a full description of the panel-of-normals filter, see the supplementary material of (Ellrott et al., 2018). (MAF filter `broad_PoN_v2`)

(ii) ExAC: site appeared in ExAC (Lek et al., 2016) ($\approx$ 60,000 normal samples) with allele count $\geq 50$. This gave us greater power than the PoN to filter germline polymorphisms somehow missing from the matched normal. (MAF filter `common_in_exac`)

(iii) Low normal coverage: sequencing coverage of the matched normal at the position of the somatic variant was fewer than eight reads, making it difficult to distinguish the somatic call from a rare germline polymorphism. (MAF filter `ndp`)

(iv) OxoG: variant was likely a sequencing artifact caused by oxidative damage during shearing in library preparation. For a full description of this artifact mode, see (Costello et al., 2013). (MAF filter `oxog`)

In total, 861,664 failing mutations were excluded, leaving us with 2,988,861 passing sSNVs.

In addition to removing MC3 flagged calls, we applied two additional filters not employed in MC3:

(i) We performed another round of PoN filtering using a panel comprising exomes captured using Illumina's ICE protocol, which was used only for TCGA samples sequenced late in the project. The ICE protocol introduces recurrent artifacts not observed in exomes captured using Agilent's capture kit, which was used for the majority of TCGA and comprises the entirety of the PoN used by MC3.

(ii) We found that many recurrent mutations were in fact false positives caused by read misalignments, verifying this by BLATing reads supporting the variant calls and seeing that they mapped with fewer mismatches elsewhere in the reference genome. To mitigate this problem, we excluded calls in 75mer windows (chosen to match the typical read length of TCGA exomes) that were not completely unique (Derrien et al., 2012) in the reference.

This left us with 2,288,080 analysis-ready sSNVs.

Finally, we annotated all mutations' gene names and protein-coding effects according to our own transcript definitions, namely all GENCODE v19 protein coding transcripts with unambiguous translation start/stop sites and splice sites.

We re-annotated because accurate calculation of dN/dS (STAR★Methods C) requires knowing the precise number of sequence contexts in each gene that can give rise to each coding effect (e.g., number of T(C→T)G sites that yield synonymous mutations), weighted by sequencing coverage across the cohort. Such exact transcript definitions were unavailable in MC3, whose calls were annotated using a pipeline that does not expose them.

Furthermore, these exact transcript definitions are required for most conservatively estimating overall protein-coding effect distributions (STAR★Methods B), which requires accounting for loci in overlapping transcripts that yield different protein-coding effects. We annotated mutations with the most deleterious effect across all transcripts at each position (synonymous $<$ missense $<$ nonsense) to avoid erroneously misclassifying mutations that are synonymous on one transcript but protein-altering on a different overlapping transcript.

### STAR★Methods B: Exome-wide Coding Effects

Given a set of genes and their observed coding mutations, we wish to calculate the distribution of the expected fractions of protein-coding effects (i.e., synonymous, missense, nonsense), assuming the mutations were distributed uniformly throughout the coding sequences of these genes, conserving sequence context (in order to normalize for mutational processes).

For example, in the cartoon coding sequence (Figure 5), the three effects of all possible strand-collapsed T(C→T)G mutations (highlighted in blue) are: (i) a synonymous substitution at codon 4 (Ile.4→Ile); (ii) a missense substitution at codon 7 (Arg.7→Gln, note mutation is on the reverse complement strand, shown as hatching); and (iii) a nonsense substitution also at codon 7 (Arg.7→Stop).

Assuming every TCG in the cartoon sequence is equally mutable, we would therefore expect random T(C→T)G mutations to result in an even distribution of coding effects — 1/3 synonymous, 1/3 missense, and 1/3 nonsense substitutions.

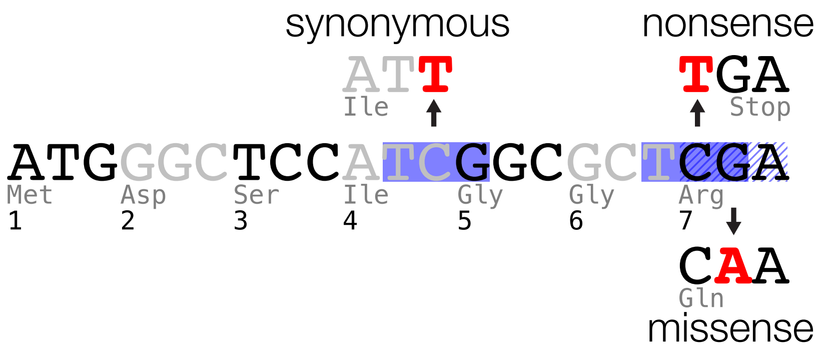


**Figure 5:** Cartoon illustrating the coding effect distribution of random C→T substitutions at strand-collapsed TCG contexts. Codons are alternately colored black/grey; TCG sites are highlighted in blue, with hatching to indicate a TCG on the nonreference strand.

We can calculate the expected distribution of coding effects across a set of genes, if we assume that the variability of mutation rates across genes is independent from the gene-specific tendency to generate synonymous, missense, or nonsense mutations. Given a large enough set of genes, this assumption is reasonable.

#### B.1 Calculating expected effect fractions

For each trinucleotide context $t$, we calculate $F_t(e_j|c_i)$, the fraction of genomic positions that match $t$ across the whole exome (or a subset of genes, e.g., known cancer genes only) that would yield coding effect $e_j$, given base change $c_i$, where $j \in \{1, 2, 3\}$ corresponds to the three coding effects (synonymous/missense/nonsense), and $i \in \{1,2,3\}$ corresponds to the three possible base substitutions. We compute these fractions by enumerating the codon changes for every possible substitution at every coding position in our gene set.

For a given set of observed coding mutations at context $t$ within this gene set, we denote the fraction that are of the $i$th base change as $f_{t,c_i}$. For example, if 95% of observed mutations at $t = \mathrm{TCG}$ are $c_i = \mathrm{C} \to \mathrm{T}$, then $f_{\mathrm{TCG,C\to T}} = 0.95$.

For each of the three coding effects $e_j$, the joint probability of seeing $\mathbf{n}_{t,e} \equiv \{n_{t,e_1}, n_{t,e_2}, n_{t,e_3}\}$ mutations given $N_t$ total mutations within context $t$ is multinomial distributed

$$\mathbf{n}_{t,e}|\mathbf{F}_t, \mathbf{f}_t, N_t \sim \mathrm{mult}(\underbrace{\textstyle\sum_i F_t(e_1|c_i) f_{t,c_i}}_{p(e_1|\mathbf{f}_t)}, \textstyle\sum_i F_t(e_2|c_i) f_{t,c_i}, \sum_i F_t(e_3|c_i) f_{t,c_i}, N_t).$$

However, because some trinucleotide contexts have low mutation counts for a given base change, there can be considerable uncertainty on $f_{t,c_i}$. We quantify this uncertainty by modeling the $f$'s not as single values, but drawn from a distribution. The ideal distribution for the $f$'s is a Dirichlet distribution, which quantifies the uncertainty on multinomial probabilities derived from counts, defining the hierarchical model

$$\begin{aligned}\mathbf{n}_{t,e}|\mathbf{F}_t, \mathbf{f}_t, N_t &\sim \mathrm{mult}(p(e_1|\mathbf{f}_t), p(e_2|\mathbf{f}_t), p(e_3|\mathbf{f}_t), N_t)\\ \mathbf{f}_t|\mathbf{n}_{t,c} &\sim \mathrm{dir}(n_{t,c_1}+1, n_{t,c_2}+1, n_{t,c_3}+1).\end{aligned}$$

We obtain the probability distribution on $\mathbf{n}_{t,e}$ by marginalizing over $\mathbf{f}_t$, i.e.,

$$\begin{aligned}p(\mathbf{n}_{t,e}|\mathbf{n}_{t,c}, N_t) &= \int d\mathbf{f}_t\ \mathrm{mult}\left(n_{t,e_1}, n_{t,e_2}, n_{t,e_3}|p(e_1|\mathbf{f}_t), p(e_2|\mathbf{f}_t), p(e_3|\mathbf{f}_t), N_t\right)\\ &\qquad \times \mathrm{dir}(f_{t,c_1}, f_{t,c_2}, f_{t,c_3}|n_{t,c_1}+1, n_{t,c_2}+1, n_{t,c_3}+1)\\ &= \int d\mathbf{f}_t \prod_{j=1}^{3} \frac{p(e_j|\mathbf{f}_t)^{n_{t,e_j}}}{n_{t,e_j}!} \prod_{j=1}^{3} f_{t,c_j}^{n_{t,c_j}}.\end{aligned} \tag{B.1.1}$$

Since there is no closed form for this integral over the 2D Dirichlet simplex, we compute it by sampling the full joint distribution $p(\mathbf{n}_{t,e}, \mathbf{f}_t|\mathbf{n}_{t,c}, N_t)$ via Markov Chain Monte Carlo (MCMC), and only considering the values drawn for $\mathbf{n}_{t,e}$. Our full conditionals are

$$\begin{aligned}p(\mathbf{n}_{t,e}|-) &\propto \prod_{j=1}^{3} \frac{p(e_j|\mathbf{f}_t)^{n_{e_j}}}{n_{e_j}!}\\ &\propto \mathrm{mult}(n_{t,e_1}, n_{t,e_2}, n_{t,e_3}|p(e_1|\mathbf{f}_t), p(e_2|\mathbf{f}_t), p(e_3|\mathbf{f}_t), N_t)\\ p(\mathbf{f}_t|-) &\propto \prod_{j=1}^{3} p(e_j|\mathbf{f}_t)^{n_{t,e_j}} \prod_{j=1}^{3} f_{t,c_j}^{n_{t,c_j}},\end{aligned}$$

the first of which we can sample from directly, the second of which cannot be analytically sampled from and requires Metropolis-Hastings sampling.

To compute the overall expected fraction of coding effects for the set of observed mutations, we convolve over the count distributions for the 32 strand-collapsed trinucleotides and 3 substitutions from (B.1.1),

$$n_{\mathrm{tot},e_1}, n_{\mathrm{tot},e_2}, n_{\mathrm{tot},e_3} \sim \mathop{\Huge *}_{t=1}^{32} \mathop{\Huge *}_{c_i=1}^{3} p(\mathbf{n}_{t,e}|\mathbf{n}_{t,c}, N_t) \tag{B.1.2}$$

and then consider the fractions of each effect type (i.e., after dividing the counts by $N = \sum_t N_t$) as the Dirichlet distribution

$$f_{\mathrm{tot},e_1}, f_{\mathrm{tot},e_2}, f_{\mathrm{tot},e_3} \sim \mathrm{dir}(n_{\mathrm{tot},e_1}, n_{\mathrm{tot},e_2}, n_{\mathrm{tot},e_3}).$$

#### STAR★Methods C: Gene dn/ds Calculation

In the previous section, we computed the expected *fractions* of protein coding effects, given a set of mutations and their contexts/substitutions. A related but distinct problem is computing whether the nonsynonymous mutation *density* significantly exceeds the synonymous mutation density for a given gene. This ratio-of-densities, referred to as $dN/dS$ in molecular evolution, is a good proxy for inferring whether a gene as a whole is under any kind of selective pressure:

- A gene under **neutral selection** will be mutated randomly and will have no bias towards generating nonsynonymous over synonymous events, after normalizing for the gene sequence and codon structure. Therefore, the $dN/dS$ of genes under neutral selection will be very close to 1.
- A gene under **positive selection** will have an excess of nonsynonymous events, relative to synonymous events, since phenotypes that change fitness (positive or negative) arise overwhelmingly more often from protein altering mutations than from synonymous mutations. Therefore, the $dN/dS$ of genes under positive selection will exceed 1.
- A gene under **negative selection** will have a dearth of nonsynonymous events, since protein altering substitutions would be selected out, keeping relatively more synonymous substitutions. Therefore, $dN/dS$ of genes under negative selection will be less than 1.

We are interested in identifying genes that are confidently drivers (to define a true positive set) or genes that are confidently passengers (for a true negative set). As in the previous section, we are not simply interested in a point estimate of $dN/dS$ for a given gene, but rather its distribution. It is crucial to estimate the *uncertainty* on $dN/dS$ to confidently conclude whether a gene is indeed under positive or neutral selection. Uncertainty in $dN/dS$ decreases as the number of mutations in the gene increases.

##### C.1 Computing somatic dN/dS

As mentioned previously, we must normalize for heterogeneous mutation frequencies at different genomic contexts and the codon structure of the gene. For each of the 96 trinucleotide channels (16 contexts × 6 base changes), the expected number of mutations with coding effect $e$ in gene $g$ is simply

$$\lambda_{g,e,c} \equiv \langle n_{g,e,c} \rangle = r_c N_{g,e,c},$$

where $r_c$ is the exome-wide mutation frequency for channel $c$ and $N_{g,e,c}$ is the number of positions (corrected for sequencing coverage) in the ORF of $g$ within channel $c$ that would yield a codon substitution of a particular effect $e$ (i.e., synonymous, missense, nonsense).

If mutation frequencies were constant across all genes, then the observed number of mutations for channel $c$/effect $e$ in any gene would be Poisson distributed around this average, i.e.,

$$n_{g,e,c} \sim \text{pois}(\lambda_{g,e,c}). \tag{C.1.1}$$

We know this is not the case, as different genes have different background mutabilities, so the overall number of mutations could be higher or lower than this expectation. Furthermore, driver genes are under positive selection, so their number of nonsynonymous mutations could be higher than this expectation. We correct for this heterogeneity with a gene-specific scale factor we infer from the data.

We assume background mutability/selective pressure is channel-independent — for example, we do not expect a more highly mutable gene to be more mutable only with respect to T(C→T)G mutations, or a driver gene to be enriched for nonsynonymous events preferentially for A(A→T)G mutations. Because of this channel-independence assumption, we scale $\lambda_{g,e,c}$ by a channel-agnostic correction factor $\beta_{g,e}$, so the Poisson distribution in (C.1.1) becomes

$$n_{g,e,c} \sim \text{pois}(\beta_{g,e}\lambda_{g,e,c}). \tag{C.1.2}$$

$\beta_{g,e}$ reflects the overall excess/dearth of mutations with coding effect $e$ in gene $g$. For example, $\beta_{g,\text{nonsyn}}$ close to 1 means that the number of nonsynonymous mutations is close to what we expect given average mutation frequencies; values significantly higher than 1 indicate an excess of nonsynonymous events; values close to 0 indicate a paucity. Of course, $\beta_{g,\text{nonsyn}} \gg 1$ is not necessarily indicative of positive selection — a highly mutable gene under neutral evolution would have a comparable excess of both nonsynonymous *and* synonymous events. To infer selective pressure on gene $g$, we are interested in the *ratio* of these scale factors,

$$dN/dS \equiv \frac{\beta_{g,\text{nonsyn}}}{\beta_{g,\text{syn}}}.$$

We calculate $\beta_{g,e}$ from its likelihood, which is the product of (C.1.2) over the 96 channels,

$$\begin{aligned}
\mathcal{L}(\beta_{g,e}|\{n_{g,e,c_i}, \lambda_{g,e,c_i}\}_{i\in\{1\ldots 96\}}) &= \prod_{i=1}^{96} \text{pois}(n_{g,e,c_i}; \beta_{g,e}\lambda_{g,e,c_i}) \\
&\propto \prod_{i=1}^{96} (\beta_{g,e}\lambda_{g,e,c_i})^{n_{g,e,c_i}} \exp(-\beta_{g,e}\lambda_{g,e,c_i}) \\
&\propto \beta_{g,e}^{\sum_i n_{g,e,c_i}} \exp(-\beta_{g,e}\sum_i \lambda_{g,e,c_i}).
\end{aligned}$$

For genes with many mutations, we could simply find the maximum likelihood estimate of $\beta_{g,e}$, as the likelihood would be sharply peaked. However, since many genes have only a few mutations and have considerable uncertainty around the maximum likelihood estimate (MLE), we use the full posterior on $\beta_{g,e}$. Luckily, the Poisson likelihood is proportional to the gamma distribution, allowing us to explicitly define this posterior:

$$\begin{aligned}
p(\beta_{g,e}; \mathbf{n}_{g,e}, \boldsymbol{\lambda}_{g,e}) &\propto \mathcal{L}(\beta_{g,e}|\mathbf{n}_{g,e}, \boldsymbol{\lambda}_{g,e}) \\
&\propto \beta_{g,e}^{\sum_i n_{g,e,c_i}} \exp(-\beta_{g,e}\sum_i \lambda_{g,e,c_i}) \\
&\propto \text{gamma}(\beta_{g,e}|1 + \sum_i n_{g,e,c_i}, \sum_i \lambda_{g,e,c_i}).
\end{aligned}$$

We can then treat both $\beta_{g,\text{nonsyn}}$ and $\beta_{g,\text{syn}}$ as gamma random variables, with the ratio $dN/dS$ their quotient,

$$\frac{dN}{dS} \sim \frac{\beta_{g,\text{nonsyn}}}{\beta_{g,\text{syn}}} \qquad \begin{array}{l} \beta_{g,\text{nonsyn}} \sim \text{gamma}(1 + \sum_i n_{g,\text{nonsyn},c_i}, \sum_i \lambda_{g,\text{nonsyn},c_i}) \\ \beta_{g,\text{syn}} \sim \text{gamma}(1 + \sum_i n_{g,\text{syn},c_i}, \sum_i \lambda_{g,\text{syn},c_i}) \end{array} .$$

Finally, we obtain the distribution of $dN/dS$ via a Monte Carlo simulation.

We can generalize this approach to infer the distribution of arbitrary protein-coding effect ratios — for example, whether genes are enriched for *truncating* mutations relative to synonymous mutations, thereby increasing sensitivity to detect tumor suppressors, which often harbor truncating events.

#### STAR★Methods D: Description of Significance Methods

In this section, we detail the four models presented in the main text: the two conventional methods, **uniform-within-gene** and **uniform Poisson regression**, which both assume that base-wise mutation frequencies have no latent variability beyond what can be predicted by sequence context and genomic covariates, and the two overdispersed methods **gamma-Poisson regression** and **log-normal-Poisson regression**, which allow mutation frequencies to have additional underlying variability beyond what the covariates predict (modeled as gamma or log-normal distributions, respectively).

##### D.1 Uniform-within-gene model

The uniform-within-gene model assumes that all sites of the same $k$-mer context in the same gene are equally mutable. There are many methods that employ this conventional assumption, with varying implementations (Miller et al., 2015; Van den Eynden et al., 2015; Lohr et al., 2012; Lawrence et al., 2014; Chang et al., 2016; Baeissa et al., 2017; Araya et al., 2016).

One particular method described by (Chang et al., 2016) assumes that the background mutation rate at a given position with trinucleotide context $t$ in gene $g$ is proportional to the average exome-wide mutation rate across all contexts $t$, weighted by a gene- and codon-specific mutability factor. In other words, it assumes that every context/codon combination $t, c$ within gene $g$ is equally mutable. It also does not account for differing substitution rates, e.g., it considers all mutations at TCG contexts equally likely, irrespective of base change.

Because we consider mutations solely on the positional level, not on the codon level, we re-implemented a method similar to that of Chang et al., which we call **uniform-within-gene**, that does not include codon-specific factors, but does account for different substitution rates.

Let $n_{s,c}$ equal the total number of mutations with base substitution $s$ at $k$-mer context $c$, and $N_c$ equal the total number of contexts $c$ in the exome. The overall mutation rate for this context/substitution is

$$r_{s,c} = n_{s,c}/N_c. \tag{D.1.1}$$

If all genes were equally mutable, the expected number of mutations $x_{s,c}$ at any genomic position of context $c$ with base substitution $s$ would simply be Poisson distributed (assuming binomial convergence) with rate parameter $r_{s,c}$,

$$x_{s,c} \sim \text{pois}(r_{s,c}). \tag{D.1.2}$$

The uniform-within-gene model accounts for the fact that different genes have different intrinsic mutabilities by scaling $r_{s,c}$ by a gene-specific mutability factor $F_g$, which explicitly assumes that all contexts/substitutions within the same gene are equally scaled. Suppose the coding sequence of gene $g$ contains $N_{g,c}$ positions of context $c$. Assuming no gene-specific scaling, the expected total number of mutations in $g$ would be $\lambda_{g,s,c} = r_{s,c}N_{g,c}$, with $r_{s,c}$ from (D.1.1). The observed numbers of mutations $x_{g,s,c}$ for this gene/context/substitution is again Poisson distributed around this average value, but this time scaled by an unknown parameter $F_g$:

$$x_{g,s,c} \sim \text{pois}(\lambda_{g,s,c} F_g).$$

To calculate $F_g$, we first compute its likelihood over all $4^k/2$ strand-collapsed contexts $\times 3$ substitutions (equal to 96 for $k = 3$),

$$\mathcal{L}(F_g | \{x_{i,j}\}) = \prod_{i=1}^{4^k/2} \prod_{j=1}^{3} \text{pois}(x_{i,j} | \lambda_{g,i,j} F_g).$$

Next, we find the maximum likelihood estimate for $F_g$,

$$\hat{F}_g = \frac{\sum_{i,j} x_{g,i,j}}{\sum_{i,j} \lambda_{g,i,j}}. \tag{D.1.3}$$

Scaling the rate parameter of (D.1.2) by (D.1.3), we can compute a $p$-value (i.e., the probability of seeing at least the observed number of mutations by chance) for a particular gene, site, and substitution type by plugging in the observed number of mutations, $\tilde{x}_{g,c,s}$, into the upper cumulative distribution function (CDF) of the Poisson distribution

$$P(\tilde{x}_{g,c,s}) \equiv \Pr(x \geq x_{g,c,s}) = \sum_{x=\tilde{x}_{g,c,s}}^{\infty} \text{pois}(x | r_{s,c} \hat{F}_g).$$

. . . . . . . . . . . . . . . . . .

#### D.2 Uniform Poisson regression model

Rather than estimate the background mutability of a gene from its own mutation burden as in the uniform-within-gene model, we can estimate it by regressing against covariates known to influence background mutability. Because covariates can be associated with genomic regions of different scales (e.g., from the single base-pair to entire chromatin regions), this approach is not limited to learning mutability solely on the gene scale, as is the case with the uniform-within-gene model, but rather allows pooling information from genomic regions of arbitrary scales.

We implement this model via standard Poisson regression, i.e., a fixed effect general linear model (GLM) with a log link function. For a given genomic position $i$ mutated $x_{i,k}$ times across the cohort with context and substitution $k$ and covariate vector $\vec{c}_i$, we assume the observed mutation count is Poisson distributed, with the log of its rate parameter linearly defined by the covariates, i.e.,

$$x_{i,k} | \vec{c}_i \sim \text{pois}(\lambda_{i,k}) \qquad \log \lambda_{i,k} = \beta_{k0} + \vec{\beta}_k \cdot \vec{c}_i. \tag{D.2.1}$$

To find the intercept $\beta_{k0}$ and slope $\vec{\beta}_k$ parameters, we maximize their likelihood function with respect to all exonic positions of context and substitution $k$, whose set we denote $\mathcal{E}_k$:

$$\mathcal{L}(\beta_{k0}, \vec{\beta}_k | \{x_{i,k}\}, \vec{\mathbf{c}}) = \prod_{i \in \mathcal{E}_k} \text{pois}(x_{i,k} | \exp(\beta_{k0} + \vec{\beta}_k \cdot \vec{c}_i)).$$

This likelihood function is log-convex so its maximum $\hat{\beta}_{k0}, \hat{\vec{\beta}}_k$ can be found via simple optimization methods, e.g., gradient descent. In practice, we use Newton-Raphson because the number of parameters is low enough that computing their full Hessian ($O(n^2)$ in number of parameters) is feasible.

Once we have found our model parameters for each $k$, we can assign a $p$-value for any position with context and substitution $k$ with counts $\tilde{x}_{i,k}$ and specific covariates $\tilde{\vec{c}}_i$ from the upper CDF of the Poisson distribution, as before:

$$P(\tilde{x}_{i,k}|\tilde{\vec{c}}_i) \equiv \Pr(x \geq \tilde{x}_{i,k}|\tilde{\vec{c}}_i) = \sum_{x=\tilde{x}_{i,k}}^{\infty} \text{pois}(x| \exp(\hat{\beta}_{k0} + \hat{\vec{\beta}}_k \cdot \tilde{\vec{c}}_i)).$$

. . . . . . . . . . . . . . . . . .

#### D.3 Gamma-Poisson regression model

The previous two models both assume that the base-wise mutability (i.e., the Poisson distribution's rate parameter) is fixed once the model parameters are found. However, this assumption does not allow for additional uncertainty in the base-wise mutability. To allow for additional uncertainty, we can probabilistically model the base-wise mutability using an arbitrary distribution $p(\boldsymbol{\theta})$. This forms a hierarchical model for observed the mutation counts $x$,

$$x \sim \text{pois}(\lambda) \qquad \lambda \sim p(\boldsymbol{\theta})$$

whose probability density function (PDF) is the compound distribution

$$p(x|\boldsymbol{\theta}) = \int_0^\infty d\lambda\ \text{pois}(x|\lambda)p(\lambda|\boldsymbol{\theta}), \tag{D.3.1}$$

with the parameters $\boldsymbol{\theta}$ of the latent distribution of the base-wise mutability learned from the data, e.g., using maximum likelihood.

Unfortunately, the integral in (D.3.1) is only analytically tractable for a few choices of $p(\lambda|\boldsymbol{\theta})$. One common choice is the gamma distribution; as mentioned in the main text, there is no intrinsic biological motivation for using this distribution — it is merely mathematically convenient. If $p(\lambda|\boldsymbol{\theta}) \equiv \text{gamma}(\lambda|a, b)$, (D.3.1) becomes

$$\begin{aligned} p(x|a,b) &= \int_0^\infty d\lambda\ \text{pois}(x|\lambda)\,\text{gamma}(\lambda|a,b) \\ &= \frac{1}{x!}\frac{b^{-a}}{\Gamma(a)}\int_0^\infty d\lambda\ \exp(-\lambda)\lambda^x \times \exp(-\lambda/b)\lambda^{a-1}. \end{aligned} \tag{D.3.2}$$

This integral is of the form $\int_0^\infty dx\ \exp(-ax)x^{b-1} = a^{-b}\Gamma(b)$, so (D.3.2) becomes

$$\begin{aligned} p(x|a,b) &= \frac{1}{x!}\frac{b^{-a}}{\Gamma(a)}\left(\frac{b}{1+b}\right)^{x+a}\left(\frac{b}{1+b}\right)^{a}\Gamma(a+x) \\ &\Downarrow \\ p(x|a,p) &= \frac{\Gamma(a+x)}{\Gamma(x+1)\Gamma(a)}p^x(1-p)^a, \end{aligned}$$

the PDF of the negative binomial distribution for $p = b/(1+b)$. Unlike the Poisson distribution, which is immediately amenable to parameterization as a GLM whose log mean can be linearly parameterized by covariates (D.2.1), the negative binomial must be re-written in terms of its mean, $\mu = pa/(1-p)$, yielding the GLM

$$p(x_{i,k}|a, \beta_0, \vec{\beta}, \vec{c}_i) \propto \left(\frac{\mu_i}{a+\mu_i}\right)^{x_{i,k}} \left(\frac{a}{a+\mu_i}\right)^{a} \qquad \log \mu_{i,k} = \beta_{k0} + \vec{\beta}_k \cdot \vec{c}_i.$$

As before, optimal values for model parameters (intercept $\beta_{k0}$, slope vector $\vec{\beta}_k$, and dispersion parameter $a$) can be found via maximum likelihood estimation. Finally, we assign $p$-values as with the other methods by computing the upper CDF of the negative binomial distribution parameterized by its MLE values.

. . . . . . . . . . . . . . . . . .

#### D.4 Log-normal-Poisson regression model

A more principled choice for the latent distribution for the base-wise mutability is the log-normal distribution. Mutations are the product of independent consecutive events, each with an independent probability of occurring; by the geometric central limit theorem, this product of probabilities approaches a log-normal distribution (Sutton, 1997).

As in the uniform Poisson regression model (D.2.1), the covariates linearly determine the log of the Poisson rate, but this time we include an additional linear factor $\epsilon_{i,k}$, which is a random variable that represents additional latent mutation rate variability at position $i$ (with context and substitution $k$) not explained by the covariates. For genomic position $i$ mutated $x_i$ times with covariate vector $\vec{c}_i$ in channel $k$, our model is

$$x_{i,k}|\vec{c}_i \sim \text{pois}(\lambda_{i,k}) \qquad \log \lambda_{i,k} = \vec{\beta}_k \cdot \vec{c}_i + \mu_k + \sigma_k \epsilon_{i,k} \qquad \epsilon_{i,k} \sim \mathcal{N}(0,1).$$

The PDF of $x_{i,k}$ is then

$$p(x_{i,k}|\mu_k, \sigma_k, \vec{\beta}_k, \vec{c}_i) = \int_0^\infty d\epsilon_{i,k}\ \text{pois}(x_{i,k}|\exp(\vec{\beta}_k \cdot \vec{c}_i + \mu_k + \sigma_k \epsilon_{i,k}))\mathcal{N}(\epsilon_{i,k}|0,1). \quad \text{(D.4.1)}$$

Since $\epsilon_{i,k}$ is a standard normal random variable, $\exp(\vec{\beta}_k \cdot \vec{c}_i + \mu_k + \sigma_k \epsilon_{i,k})$ is log-normally distributed with geometric mean $\vec{\beta}_k \cdot \vec{c}_i + \mu_k$ and geometric standard deviation $\sigma_k$. These parameters are interpretable: $\mu_k$ represents the overall geometric mean mutation frequency for context/substitution $k$, $\sigma_k$ the overall geometric standard deviation around this mean (i.e., the average multiplicative distance from the mean), and $\vec{\beta}_k$ the geometric covariate slope vector associated with $k$ (i.e., the multiplicative effect each covariate has on the mutation rate).

In addition to being better statistically calibrated than the other methods (see **Results**), the log-normal-Poisson model carries other advantages:

1. Because $\sigma_k$ is multiplicative, it quantifies overdispersion irrespective of mutation frequency, allowing for comparisons between different mutational processes. For example, $\exp(\sigma_k) = 2$ indicates that bases in context-substitution pair $k$ that are one geometric standard deviation above the mean will be approximately *twice* as mutable as sites at the mean, while bases falling one g. standard deviation below the mean will be approximately *half* as mutable, regardless of the mean mutation frequency.

2. The log-normal-Poisson model accommodates nested variance (variance is additive). Suppose that we are analyzing mutations within trinucleotide context $c$. We would fit the model to all exonic sites of that sequence context. Now suppose that we would like to account for additional variance explained by the 16 flanking pentanucleotide contexts. The set of base-pairs belonging to the trinucleotide context $c$ is a superset of the base-pairs belonging to a pentamer context flanking that trinucleotide. Thus, we can express the total variance for base-pair $i$ at context/substitution $k$ as the sum of the overall trinucleotide variance and the additional variance explained by the pentamer context of $i$:

$$\log \lambda_{i,k} = \vec{\beta}_k \cdot \vec{c}_i + \mu_{k_{\text{pent}}} + \sigma_{k_{\text{tri}}} \epsilon_{i_{\text{tri}}} + \sigma_{k_{\text{pent}}} \epsilon_{i_{\text{pent}}}, \tag{D.4.2}$$

with $\epsilon_{i_{\text{tri}}}$ and $\epsilon_{i_{\text{pent}}}$ representing independent standard normal random variables. The total variance for the pentamer context is $\sigma^2_{k_{\text{tri}}} + \sigma^2_{k_{\text{pent}}}$, since the variance of the sum of independent normal random variables is the sum of their variances. In general linear mixed model parlance, the pentamers are *categorical random effects.*

3. The natural parameterization in terms of mean and variance allows us to naturally quantify the amount of variance explained by model factors (e.g., covariates). The amount that $\sigma^2$ drops after adding a covariate is equivalent to the variance explained by that covariate, given the other covariates already present in the model. To obtain the average linear contribution of each covariate, we fit the model for all possible sets of covariates, and then perform an ordinary least squares fit to find each marginal linear contribution. For example, if there are three covariates, we fit a model for each of the $2^3 = 8$ possible combinations, obtaining $\sigma_0^2 \ldots \sigma_7^2$, the observed values of model parameter $\sigma^2$ for each covariate set (0: no covariates, 7: all covariates). We then fit the following linear model,

$$\begin{bmatrix} 1 & 0 & 0 & 0 \\ 1 & 0 & 0 & 1 \\ 1 & 0 & 1 & 0 \\ 1 & 0 & 1 & 1 \\ \vdots & \vdots & \vdots & \vdots \\ 1 & 1 & 1 & 1 \end{bmatrix} \vec{\sigma}^2_{\text{lin}} = \begin{bmatrix} \sigma_0^2 \\ \sigma_1^2 \\ \sigma_2^2 \\ \sigma_3^2 \\ \vdots \\ \sigma_7^2 \end{bmatrix},$$

with a 1 in the data matrix representing the presence of each covariate. The first column is the intercept and thus entirely comprises ones. $\vec{\sigma}^2_{\text{lin}}$ is then the marginal linear contribution of each covariate; its first element (i.e., the intercept) is the remaining amount of unexplained variance after all covariates have been incorporated; the other three elements are the variance explained by their respective covariate.

To fit the log-normal-Poisson model's parameters, we employ a fully Bayesian approach, sampling from the parameters' posterior distribution $p(\mu_k, \sigma_k, \vec{\beta}_k | \{x_{i,k}\}, \{\vec{c}\})$ via MCMC. We use MCMC instead of a standard likelihood (or quasi-likelihood) approach because:

1. The model parameters provide meaningful summary statistics of the base-wise mutation rate; as such, we are not merely interested in the model's predictions (i.e., significance values), but also its parameters. It is therefore useful to not only obtain their optimal best-fit values, but their overall posterior distribution, to quantify our confidence in the model.

2. The integral in (D.4.1) is analytically intractable, but it is amenable to approximation via Gibbs sampling, given the hierarchical nature of the model.

We describe the MCMC in detail in STAR★Methods E.

To compute $p$-values for the previous three models, we first find a point estimate $\hat{\boldsymbol{\theta}}$ of optimal model parameters (e.g., via maximum likelihood estimation), and then use the upper CDF parameterized by $\hat{\boldsymbol{\theta}}$, for an observed number of mutations $\tilde{x}_i$ with covariates $\tilde{\vec{c}}_i$ at base-pair $i$,

$$P(\tilde{x}_i|\tilde{\vec{c}}_i) = \sum_{x=\tilde{x}_i}^{\infty} p(x|\hat{\boldsymbol{\theta}}, \tilde{\vec{c}}_i).$$

Because the log-normal-Poisson model is fully Bayesian, we no longer use a point estimate for $\hat{\boldsymbol{\theta}}$, but rather integrate the CDF over the full domain $\boldsymbol{\Theta}$ of the posterior on $\boldsymbol{\theta}$, yielding posterior predictive $p$-values

$$P(\tilde{x}_i|\tilde{\vec{c}}_i) = \sum_{x=\tilde{x}_i}^{\infty} \int_{\boldsymbol{\Theta}} d\boldsymbol{\theta}\, p(x|\boldsymbol{\theta}, \tilde{\vec{c}}_i) p(\boldsymbol{\theta}|\{x\}, \{\vec{c}\}). \tag{D.4.3}$$

#### STAR★Methods E: MCMC Implementation

In this section, we detail the implementation of the Markov Chain Monte Carlo (MCMC) sampler used to fit the log-normal-Poisson regression model. As with most non-conjugate hierarchical models, the log-normal-Poisson probability density function (D.4.1) is analytically intractable — its most general form is the compound distribution

$$p(x|\mu,\sigma) = \int_0^\infty d\epsilon \; \mathrm{pois}(x|\epsilon) \log\mathcal{N}(\epsilon|\mu,\sigma), \tag{E.0.1}$$

which has no closed form, so analytically sampling any distribution derived from the log-normal-Poisson PDF (e.g., the posterior on model parameters) will also lack closed form.

#### E.1 Posterior Calculation

**E.1.a** *Model statement.*
Recall the model hierarchy of the general linear mixed model defined in STAR★Methods **D.4**. For observed number of mutated patients $x_i$ at the $i$th base-pair whose covariate vector is $\vec{c}_i$, we have

$$x_i|\vec{c}_i \sim \mathrm{pois}(\lambda_i) \qquad \log\lambda_i = \vec{\beta}\cdot\vec{c}_i + \mu + \sigma\epsilon_i \qquad \epsilon_i \sim \mathcal{N}(0,1).$$

For a set $\{x\}$ of $n$ base-pairs with covariates $\{\vec{c}\}$, the full posterior distribution across all parameters — i.e., including the latent $\epsilon$ in (E.0.1) or (D.4.1) — is

$$p(\mu,\sigma,\vec{\beta},\epsilon_1,\ldots,\epsilon_n|\{x\},\{\vec{c}\},\mathcal{H}) \propto \prod_{i=1}^{n}\Big[p(x_i|\exp(\vec{\beta}\cdot\vec{c}_i+\mu+\sigma\epsilon_i))p(\epsilon_i)\Big]\,p(\mu,\sigma,\vec{\beta}|\mathcal{H}) \tag{E.1.1}$$

for set of hyperparameters $\mathcal{H}$. In the case of nested variability (i.e., when adding $m$ total categorical random effects), the model hierarchy becomes

$$x_i|\vec{c}_i,\mathbf{1}_{i\cdot} \sim \mathrm{pois}(\lambda_i) \qquad \log\lambda_i = \vec{\beta}\cdot\vec{c}_i + \sum_{j=1}^{m}(\mu_j+\sigma_j\epsilon_{ij})\mathbf{1}_{ij} + \sigma_0\epsilon_{i0} \qquad \{\epsilon_{ij}\} \sim \mathcal{N}(0,1), \tag{E.1.2}$$

where $\mathbf{1}_{ij}$ is an indicator function for whether the $i$th base is of category $j$, and the set $\{\epsilon_{ij}\}$ comprises *independent* standard normal random variables. The full posterior (E.1.1) becomes

$$\begin{aligned} &p(\mu_1,\ldots,\mu_m,\sigma_0,\sigma_1,\ldots,\sigma_m,\vec{\beta},\{\epsilon\}|\{x\},\{\vec{c}\},\mathcal{H}) \\ \propto &\prod_{i=1}^{n}\left[p(x_i|\exp(\vec{\beta}\cdot\vec{c}_i+\textstyle\sum_{j=1}^{m}(\mu_j+\sigma_j\epsilon_{ij})\mathbf{1}_{ij}+\sigma_0\epsilon_{i0}))p(\epsilon_{i0})\displaystyle\prod_{j=1}^{m}p(\epsilon_{ij})^{\mathbf{1}_{ij}}\right]p(\{\mu\},\{\sigma\},\vec{\beta}|\mathcal{H}). \end{aligned} \tag{E.1.3}$$

In our analyses, we only use the categorical random effects to represent flanking pentamer contexts within each trimer. Suppose that we fit the model to the set of all base-pair substitutions $\{x_k\}$ at trinucleotide substitution $k$, e.g. TCT→TAT. Our categories within $\{x_k\}$ are the $j = 1 \dots 16$ pentamer contexts flanking $k$ (e.g., **A**TCT**G**), which are unique for each base-pair. Then (E.1.2) becomes (D.4.2); the sum in (E.1.2) disappears, since $\mathbf{1}_{ij} = 0$ for all but a single value of $j$ — each base-pair can only have a single pentamer context. We denote the pentamer context associated with the $i$th base-pair as $j(i)$.

Explicitly writing out the full posterior distribution for the single random effect model for $n$ base-pairs in trinucleotide context $k$, we have

$$p(\mu_1, \dots, \mu_{16}, \sigma_0, \sigma_1, \dots, \sigma_{16}, \vec{\beta}, \{\epsilon\} | \{x_k\}, \{\vec{c}\}, \mathcal{H})$$
$$\propto \prod_{i=1}^{n} \Big[ p(x_i | \exp(\underbrace{\vec{\beta} \cdot \vec{c}_i + \mu_{j(i)} + \sigma_0 \epsilon_{i0} + \sigma_{j(i)} \epsilon_{ij(i)}}_{\lambda_i})) p(\epsilon_{i0}) p(\epsilon_{ij(i)}) \Big] p(\{\mu\}, \{\sigma\}, \vec{\beta} | \mathcal{H}) \quad \text{(E.1.4)}$$

By sampling from the full posterior in (E.1.4) and ignoring the samples for latent parameters $\boldsymbol{\epsilon}$, we simultaneously sample the distribution on the parameters of interest ($\{\mu\}$, $\{\sigma\}$, $\vec{\beta}$) while marginalizing out $\{\epsilon\}$, i.e., performing the integral in (E.0.1). Because these full posteriors can have millions of parameters (each $\epsilon_{ij} \in \{\epsilon\}$ corresponds to a single base-pair, and we fit the model to all base-pairs within a trinucleotide context), they are impractical to sample in a single draw. We therefore use a Metropolis-within-Gibbs scheme: we partition the joint posterior's parameters into uncorrelated blocks, and independently sample from the full conditional of each block. In the next sections, we detail these parameter blocks and their full conditionals. We will focus exclusively on the nested model in (E.1.4), since the non-nested model is a simplified subset of it.

• • • • • •

**E.1.b** *Metropolis-within-Gibbs algorithm description.*
The Gibbs sampler allows us to draw from a high dimensional probability distribution by iteratively sampling lower dimensional subsets of its parameters conditioned on all the other parameters. For example, if sampling in three dimensions from the distribution $p(x, y, z)$ is difficult, iterative univariate samples from full conditional distributions $p(x|y, z)$, $p(y|x, z)$, and $p(z|x, y)$ will converge to multivariate samples $(x, y, z)$ from the full joint. We will denote full conditional distributions as $p(x|-)$: $x$ conditioned on all other parameters in the joint, i.e., $(y, z)$.

For notational convenience, let us first expand (E.1.4) in terms of the PDFs of the Poisson and standard normal distributions corresponding to $p(x_i|\lambda_i)$ and $p(\epsilon_{ij})$, respectively. Recall that $\lambda_i$ is a function of $\mu_{j(i)}$, $\sigma_0$, $\sigma_{j(i)}$, $\vec{\beta}$, $\{\epsilon\}$, and $\vec{c}_i$; we omit the arguments for brevity.

$$p(\{\mu\}, \{\sigma\}, \vec{\beta}, \{\epsilon\} | \{x_k\}, \{\vec{c}\}, \mathcal{H}) \propto \prod_{i=1}^{n} \left[ \exp(\lambda_i x_i - \exp(\lambda_i)) \exp\left(-\frac{\epsilon_{i0}^2}{2}\right) \exp\left(-\frac{\epsilon_{ij(i)}^2}{2}\right) \right]$$
$$\times\, p(\{\mu\}, \{\sigma\}, \vec{\beta} | \mathcal{H}) \quad \text{(E.1.5)}$$

We see that the $\epsilon$'s are independent, so each $\epsilon_{ij}$ can be sampled independently as its own 1-dimensional block. To allow for correlation between each $\mu_j$ and each $\sigma_j$, we sample $(\mu_j, \sigma_j)$ as a 2-D block. Likewise, if there is correlation between covariates, we will expect correlation between elements of $\vec{\beta} \in \mathbb{R}^b$, and will therefore sample $\vec{\beta}$ as a $b$-dimensional block.

Because none of the full conditionals can be sampled from analytically, we will sample from them using the Metropolis-Hastings algorithm: given some target distribution $p(x)$ that is impossible to analytically sample from, we randomly sample $x^*$ from some easy-to-sample proposal distribution $q(x^*|x)$ that approximates $p(x)$, and accept the new value $x^*$ with probability

$$\Pr(x \to x^*) = \min\left\{1, \frac{p(x^*)q(x|x^*)}{p(x)q(x^*|x)}\right\}.$$

Iterating this procedure will yield random samples from target $p(x)$.

Of course, performance of this method critically depends on choosing a proposal distribution that well-approximates the distribution being sampled. All of the full conditionals are log-concave; furthermore, terms above second-order in their series expansions around their maxima are negligible, so we will use proposal distributions that quadratically approximate the full conditionals centered at their maxima. We find each full conditional's global maximum, and propose using a (multivariate) $t$ distribution whose mean is the global maximum and whose (co)variance is the curvature at the maximum,

$$\hat{\vec{\mu}} = \langle \hat{x}, \hat{y} \rangle = \underset{x,y}{\operatorname{argmax}}(\log p(x, y|-)) \qquad \hat{\mathbf{\Sigma}} = -\mathbf{H}^{-1}[\log p(\hat{x}, \hat{y}|-)]$$

$$q(x^*, y^*|x, y) \sim t_\nu(\hat{\vec{\mu}}, \hat{\mathbf{\Sigma}}),$$

where $\mathbf{H}^{-1}$ is the inverse Hessian. $\nu$ is the degrees-of-freedom of the $t$ distribution, which can be tuned to achieve better approximation of the target distribution.

To find the maxima, we use Newton-Raphson iterations (since we are already calculating the Hessian), augmented with backtracking linesearch to avoid potentially overshooting.

· · · · · ·

**E.1.c** *Metropolis-within-Gibbs sampling blocks.*
Here, we detail the sampling procedure used for each full conditional block.

**E.1.c.i** *$\mu_j, \sigma_j$ block.* The full conditional of each $(\mu_j, \sigma_j)$ block is

$$\log p(\mu_j, \sigma_j|-) \propto \sum_{i \in \mathbf{J}} \Big[(\mu_j + \sigma_j \epsilon_{ij})x_i - \exp(\vec{\beta} \cdot \vec{c}_i + \mu_j + \sigma_0 \epsilon_{i0} + \sigma_j \epsilon_{ij})\Big] + \log p(\mu_j, \sigma_j|\mathcal{H}).$$

Until this point, we have not specified the form of our priors. Because

$$X \sim \mu_j + \sigma_j \mathcal{N}(0, 1) \equiv X \sim \mathcal{N}(\mu_j, \sigma_j^2),$$

it makes sense to specify a joint prior $p(\mu_j, \sigma_j|\mathcal{H})$ on $(\mu_j, \sigma_j)$ conjugate to the normal distribution, to make interpretation of hyperparameters $\mathcal{H}$ easy. In this case, this is the normal-inverse-gamma distribution:

$$\mu_j, \sigma_j^2 \sim \mathcal{NG}^{-1}(A, B, M, S). \tag{E.1.6}$$

where $A$ and $B$ are the shape/scale parameters of the gamma distribution, and $M$ and $S$ are the mean/variance parameters of the normal distribution, respectively. These parameters signify that $\sigma_j^2$ was estimated from $2A$ observations with sample mean $M$ and sum of sample squared deviations $2B$; $\mu_j$ was estimated from $S$ observations with sample mean $M$. To actually implement this prior, we make change-of-variable $\tau_j = 1/\sigma_j^2$, turning the normal-inverse-gamma prior into a normal-gamma prior

$$\mu_j, \tau_j \sim \mathcal{NG}(A, B, M, S).$$

With our prior in hand, we need to compute the gradient and Hessian of the full conditional in order to perform our Newton-Raphson iterations. Our gradient is

$$\begin{aligned}
\vec{\nabla} \log p(\mu_j, \tau_j|-) &= \left\langle \frac{\partial}{\partial \mu_j} \log p(\mu_j, \tau_j|-), \frac{\partial}{\partial \tau_j} \log p(\mu_j, \tau_j|-) \right\rangle \\
\frac{\partial}{\partial \mu_j} \log p(\mu_j, \tau_j|-) &= \sum_{i \in \mathbf{J}} x_i - \exp(\mu_j) \sum_{i \in \mathbf{J}} \exp\left(\vec{\beta} \cdot \vec{c}_i + \frac{\epsilon_{i0}}{\sqrt{\tau_0}} + \frac{\epsilon_{ij}}{\sqrt{\tau_j}}\right) \\
&\quad \underbrace{- S\tau_j(\mu_j - M)}_{\frac{\partial}{\partial \mu_j} \log \mathcal{NG}(\mu_j, \tau_j | A, B, M, S)} \\
\frac{\partial}{\partial \tau_j} \log p(\mu_j, \tau_j|-) &= -\frac{1}{2\tau_j^{3/2}} \sum_{i \in \mathbf{J}} \epsilon_{ij} x_i \\
&\quad + \exp(\mu_j) \sum_{i \in \mathbf{J}} \exp\left(\vec{\beta} \cdot \vec{c}_i + \frac{\epsilon_{i0}}{\sqrt{\tau_0}} + \frac{\epsilon_{ij}}{\sqrt{\tau_j}}\right) \frac{\epsilon_{ij}}{2\tau_j^{3/2}} \\
&\quad \underbrace{+ \frac{A-1}{\tau_j} - \frac{1}{B} - \frac{S}{2}(\mu_j - M)^2 + \frac{1}{2\tau_j}}_{\frac{\partial}{\partial \tau_j} \log \mathcal{NG}},
\end{aligned}$$

and our Hessian components are

$$H_{\mu,\mu} = -\exp(\mu_j) \sum_{i \in \mathbf{J}} \exp\left(\vec{\beta} \cdot \vec{c}_i + \frac{\epsilon_{i0}}{\sqrt{\tau_0}} + \frac{\epsilon_{ij}}{\sqrt{\tau_j}}\right) \underbrace{- S\tau}_{\frac{\partial^2}{\partial \mu_j^2} \log \mathcal{NG}}$$

$$\begin{aligned}
H_{\tau,\tau} = \frac{3}{4\tau_j^{5/2}} \sum_{i \in \mathbf{J}} \epsilon_{ij} x_i + \exp(\mu_j) \sum_{i \in \mathbf{J}} \exp\left(\vec{\beta} \cdot \vec{c}_i + \frac{\epsilon_{i0}}{\sqrt{\tau_0}} + \frac{\epsilon_{ij}}{\sqrt{\tau_j}}\right) \epsilon_{ij} \left[-\frac{3}{4\tau_j^{5/2}} - \frac{\epsilon_{ij}}{4\tau_j^3}\right] \\
\underbrace{- \frac{2A-1}{2\tau_j^2}}_{\frac{\partial^2}{\partial \tau_j^2} \log \mathcal{NG}}
\end{aligned}$$

$$H_{\mu,\tau} = \exp(\mu_j) \sum_{i \in \mathbf{J}} \exp\left(\vec{\beta} \cdot \vec{c}_i + \frac{\epsilon_{i0}}{\sqrt{\tau_0}} + \frac{\epsilon_{ij}}{\sqrt{\tau_j}}\right) \frac{\epsilon_{ij}}{2\tau_j^{3/2}} \underbrace{- T(\mu_j - M)}_{\frac{\partial^2}{\partial \mu_j \partial \tau_j} \log \mathcal{NG}}$$

To actually use this gradient/Hessian to perform Newton-Raphson iterations, and since Newton-Raphson updates are only stable on functions whose domain is $\mathbb{R}$, we need to make change-of-variable $g^{-1} : \mathbb{R}^+ \to \mathbb{R},\ \tau_j \mapsto \log(\tau_j)$, because $\tau_j \in (0, \infty)$. Recall that when transforming parameters of a probability distribution with some function $g^{-1}$, we need to multiply by the Jacobian of the inverse transform $g$ to preserve the volume element. Thus, for $g^{-1}(\tau_j) = \log(\tau_j)$,

$$p(\mu_j, g(\tau_j)|-)\frac{dg}{d\tau_j} = p(\mu_j, \exp(\tau_j)|-)\exp(\tau_j).$$

As a result, our log-space gradient/Hessian are actually

$$\vec{\nabla}[\log p(\mu_j, \exp(\tau_j)|-) + \tau_j] \qquad \mathbf{H}[\log p(\mu_j, \exp(\tau_j)|-) + \tau_j]$$

Performing this change-of-variables requires invoking the chain rule. For the gradient, this is straightforward:

$$\vec{\nabla} f(x, g(y)) = \left\langle \frac{\partial f}{\partial x}, \frac{\partial f}{\partial g}\frac{\partial g}{\partial y} \right\rangle,$$

so for $g(y) = \exp(y)$, we simply multiply the untransformed $\tau$ component of the gradient ($\partial_g f$) by $\exp(\tau_j)$, and then add 1 to account for the Jacobian.

For the Hessian, we need to invoke the chain rule twice, to account for the second derivative. $H_{xx}$ is unchanged. The other components transform as follows:

$$\begin{aligned} H_{yy} &= \frac{\partial}{\partial y}\left[\frac{\partial f}{\partial g}\frac{\partial g}{\partial y}\right] & \qquad H_{xy} &= \frac{\partial}{\partial y}\left[\frac{\partial f}{\partial x}\right] \\ &= \frac{\partial f}{\partial g}\frac{\partial^2 g}{\partial y^2} + \left(\frac{\partial g}{\partial y}\right)^2 \frac{\partial^2 f}{\partial g^2} & &= \frac{\partial g}{\partial y}\frac{\partial^2 f}{\partial g \partial x} \end{aligned}$$

$\partial_g f$ is the untransformed $\tau$ component of the gradient; $\partial_y^2 g = \exp(\tau_j)$; $(\partial_y g)^2 = \exp(2\tau_j)$; $\partial_g^2 f$ is the untransformed $(\tau, \tau)$ component of the Hessian; and $\partial_{gx}^2 f$ is the untransformed $\mu, \tau$ component of the Hessian. Because the Jacobian term in the gradient is a constant 1, it disappears entirely in the Hessian.

**E.1.c.ii** $\sigma_0$ *block.* In addition to each $(\mu_j, \sigma_j)$ for each pentamer context, we also have an overall variance term $\sigma_0$ shared between the pentamers. Its full conditional is

$$\log p(\sigma_0|-) \propto \sum_{i=1}^{n}\left[\sigma_0\epsilon_{i0}x_i - \exp(\vec{\beta}\cdot\vec{c}_i + \mu_{j(i)} + \sigma_0\epsilon_{i0} + \sigma_{j(i)}\epsilon_{ij(i)})\right] + \log p(\sigma_0|\mathcal{H}).$$

We place an inverse gamma prior on $\sigma_0^2 \sim \mathcal{G}^{-1}(A, B)$, whose parameters have the same conjugate interpretation as the normal-inverse-gamma prior for $(\mu_j, \sigma_j)$, (E.1.6). As before, we perform change-of-variable $\tau_0 = 1/\sigma_0^2$ to actually implement this model structure.

Our gradient and Hessian are simply the first and second derivatives of $p(\sigma_0|-)$:

$$\begin{aligned} \frac{\partial}{\partial \tau_0}\log p(\tau_0|-) = &-\frac{1}{2\tau_0^{3/2}}\sum_{i=1}^{n}\epsilon_{i0}x_i \\ &+ \sum_{i=1}^{n}\exp\left(\vec{\beta}\cdot\vec{c}_i + \mu_{j(i)} + \frac{\epsilon_{i0}}{\sqrt{\tau_0}} + \frac{\epsilon_{ij(i)}}{\sqrt{\tau_{j(i)}}}\right)\frac{\epsilon_{i0}}{2\tau_0^{3/2}} + \underbrace{\frac{A-1}{\tau_0} - \frac{1}{B}}_{\frac{\partial}{\partial \tau_0}\log \mathcal{G}(A,B)} \end{aligned}$$

and

$$\frac{\partial^2}{\partial \tau_0^2} \log p(\tau_0|-) = \frac{3}{4\tau_0^{5/2}} \sum_{i=1}^{n} \epsilon_{i0} x_i$$
$$+ \sum_{i=1}^{n} \exp\left(\vec{\beta} \cdot \vec{c}_i + \mu_{j(i)} + \frac{\epsilon_{i0}}{\sqrt{\tau_0}} + \frac{\epsilon_{ij(i)}}{\sqrt{\tau_{j(i)}}}\right) \epsilon_{i0} \left[-\frac{3}{4\tau_0^{5/2}} - \frac{\epsilon_{i0}}{4\tau_0^3}\right]$$
$$\underbrace{-\frac{A-1}{\tau_0^2}}_{\frac{\partial^2}{\partial \tau_0^2} \log \mathcal{G}(A,B)}.$$

As before, we actually perform the Newton-Raphson iterations on $\log p(\exp(\tau_0)|-)$, transforming the PDF and first/second derivatives with the appropriate Jacobian and chain rule factors, respectively.

**E.1.c.iii** $\vec{\beta}$ *block.* Our full conditional is

$$\log p(\vec{\beta}|-) \propto \sum_{i=1}^{n} \left[\vec{\beta} \cdot \vec{c}_i x_i + \exp(\vec{\beta} \cdot \vec{c}_i + \mu_{j(i)} + \sigma_0 \epsilon_{i0} + \sigma_{j(i)} \epsilon_{ij(i)})\right] + \log p(\vec{\beta}|\mathcal{H}),$$

with $p(\vec{\beta}|\mathcal{H})$ equal to multivariate normal prior $\vec{\beta} \sim \mathcal{N}_m(\vec{\mu}_\beta, \mathbf{\Sigma}_\beta)$.

For both notational convenience and computational efficiency, we express the sums of dot products (i.e., $\sum_i \vec{\beta} \cdot \vec{c}_i$) as matrix multiplications over covariate matrix $\mathbf{C} = \begin{bmatrix}\vec{c}_1 & \cdots & \vec{c}_n\end{bmatrix}^\intercal$ and data vector $\vec{x} = \langle x_1, \ldots x_n \rangle^\intercal$. We similarly vectorize our sets of parameters and latent random variables, with parameter vectors $\vec{\sigma}_\mathbf{J} = \langle \sigma_{j(1)}, \sigma_{j(2)}, \ldots, \sigma_{j(n)} \rangle^\intercal$ and $\vec{\mu}_\mathbf{J} = \langle \mu_{j(1)}, \ldots, \mu_{j(n)} \rangle^\intercal$, and latent random variable vectors $\vec{\epsilon}_0 = \langle \epsilon_{10}, \epsilon_{20}, \ldots, \epsilon_{n0} \rangle^\intercal$ and $\vec{\epsilon}_\mathbf{J} = \langle \epsilon_{1j(1)}, \ldots, \epsilon_{nj(n)} \rangle^\intercal$.

Our gradient is

$$\vec{\nabla} p(\vec{\beta}|-) = \vec{x}^\intercal \mathbf{C} - \underbrace{\exp\odot(\mathbf{C}\vec{\beta} + \vec{\mu}_\mathbf{J} + \sigma_0 \vec{\epsilon}_0 + \vec{\sigma}_\mathbf{J} \cdot \vec{\epsilon}_\mathbf{J})^\intercal}_{\vec{q}} \mathbf{C} \underbrace{- \mathbf{\Sigma}_\beta^{-1}(\vec{\beta} - \vec{\mu}_\beta)}_{\vec{\nabla} p(\vec{\beta}|\vec{\mu}_\beta, \mathbf{\Sigma}_\beta)},$$

where $\exp\odot$ is the elementwise exponential.

Our Hessian is simply

$$\mathbf{H} = -\mathbf{C}^\intercal \operatorname{diag}(\vec{q})\mathbf{C} - \mathbf{\Sigma}_\beta^{-1}.$$

Because $\mathbf{C}$ can have millions of rows, we speed up calculation of the Hessian by noting that any matrix multiplication of the form $\mathbf{C}^\intercal\mathbf{D}\mathbf{C}$ with diagonal $\mathbf{D}$ can be expressed as

$$\mathbf{C}^\intercal\mathbf{D}\mathbf{C} = \mathbf{C}^\intercal \left(\sum_{j=1}^{n} d_j \mathbf{E}_{jj}\right) \mathbf{C} = \sum_{j=1}^{n} d_j \mathbf{C}^\intercal \mathbf{E}_{jj} \mathbf{C} = \sum_{j=1}^{n} d_j \vec{c}_j \otimes \vec{c}_j$$

where $d_j$ is the $j$th diagonal element of $\mathbf{D}$, $\mathbf{E}_{jj}$ the corresponding standard basis matrix, and $\vec{c}_j$ the $j$th row of $\mathbf{C}$. Because the covariate matrix $\mathbf{C}$ is constant, we only need to calculate each outer product $\vec{c}_j \otimes \vec{c}_j$ once ahead of time, greatly saving time when computing the Hessian, which only requires computing a linear combination of the outer product matrices.

**E.1.c.iv** $\boldsymbol{\epsilon}$ *blocks.* For each shared latent random variable $\epsilon_{i0} \in \boldsymbol{\epsilon}_0$, our full conditional is

$$\log p(\epsilon_{i0}|-) \propto \sigma_0 \epsilon_{i0} x_i - \exp(\vec{\beta} \cdot \vec{c}_i + \mu_{j(i)} + \sigma_0 \epsilon_{i0} + \sigma_{j(i)} \epsilon_{ij(i)}) - \frac{\epsilon_{i0}^2}{2}.$$

Likewise, each nested latent random variable's full conditional is

$$\log p(\epsilon_{ij(i)}|-) \propto \sigma_j \epsilon_{ij(i)} x_i - \exp(\vec{\beta} \cdot \vec{c}_i + \mu_{j(i)} + \sigma_0 \epsilon_{i0} + \sigma_{j(i)} \epsilon_{ij(i)}) - \frac{\epsilon_{ij(j)}^2}{2}.$$

Each of these full conditionals is univariate, so their gradients/Hessians will simply be the first/second derivatives, which are of the form

$$\frac{\partial}{\partial \epsilon_{i0}} \log p(\epsilon_{i0}|-) = \sigma_0 x_i - \sigma_0 \exp(\vec{\beta} \cdot \vec{c}_i + \mu_{j(i)} + \sigma_0 \epsilon_{i0} + \sigma_{j(i)} \epsilon_{ij(i)}) - \epsilon_{i0}$$
$$\frac{\partial^2}{\partial \epsilon_{i0}^2} \log p(\epsilon_{i0}|-) = -\sigma_0^2 \exp(\vec{\beta} \cdot \vec{c}_i + \mu_{j(i)} + \sigma_0 \epsilon_{i0} + \sigma_{j(i)} \epsilon_{ij(i)}) - 1,$$

substituting the appropriate $\sigma/\epsilon$ as necessary.

. . . . . . . . . . . . . . . . . .

#### E.2 Posterior predictive calculation

To actually infer whether an observed level of recurrent mutation significantly exceeds the expected background, we compute a posterior predictive $p$-value as stated in Equation (D.4.3). This entails integrating the parameters of the log-normal-Poisson distribution over the posterior probabilities of the model parameters, whose domain we denote $\boldsymbol{\Theta}$. Given some observed number of mutations $\tilde{x}$ with covariates $\tilde{\vec{c}}$ and category (i.e., pentamer context) $\tilde{\jmath}$, the posterior predictive is

$$p(\tilde{x}|\{x\}, \{\vec{c}\}, \tilde{\vec{c}}) = \int_{\boldsymbol{\Theta}} \overbrace{\left[\int_{-\infty}^{\infty} d\epsilon_0 \, d\epsilon_{\tilde{\jmath}} \; \text{pois}(\tilde{x}| \exp(\vec{\beta} \cdot \tilde{\vec{c}} + \mu_{\tilde{\jmath}} + \sigma_0 \epsilon_0 + \sigma_{\tilde{\jmath}} \epsilon_{\tilde{\jmath}})) \mathcal{N}(\epsilon_0|0,1) \mathcal{N}(\epsilon_{\tilde{\jmath}}|0,1)\right]}^{I(\tilde{x}, \tilde{c}, \mu_{\tilde{\jmath}}, \sigma_0, \sigma_{\tilde{\jmath}}, \vec{\beta})}$$
$$\times \, p(\mu_{\tilde{\jmath}}, \sigma_0, \sigma_{\tilde{\jmath}}, \vec{\beta}|\{x\}, \{\vec{c}\}, \mathcal{H}) \, d\mu_{\tilde{\jmath}} \, d\sigma_{\tilde{\jmath}} \, d\sigma_0 \, d\vec{\beta}.$$

To compute the outer integral over model parameters, we simply average draws from the MCMC:

$$\int_{\boldsymbol{\Theta}} d\mu_{\tilde{\jmath}} \, d\sigma_{\tilde{\jmath}} \, d\sigma_0 \, d\vec{\beta} \, I(\tilde{x}, \tilde{c}, \mu_{\tilde{\jmath}}, \sigma_0, \sigma_{\tilde{\jmath}}, \vec{\beta}) \approx N^{-1} \sum_{i=1}^{N} I(\tilde{x}, \tilde{c}, \mu_{\tilde{\jmath}}^{(i)}, \sigma_0^{(i)}, \sigma_{\tilde{\jmath}}^{(i)}, \vec{\beta}^{(i)}),$$

where $(i)$ is the $i$th MCMC draw.

To compute the inner integrals over latent parameters, we use Hermite quadrature. Hermite polynomials provide a basis for approximating integrals of the form $\int_{-\infty}^{\infty} dx \, f(x) \exp(-x^2)$. In our case, the inner integrand is

$$g(\epsilon_0, \epsilon_{\tilde{\jmath}}) = \text{pois}(\tilde{x}| \exp(\vec{\beta} \cdot \tilde{\vec{c}} + \mu_{\tilde{\jmath}} + \sigma_0 \epsilon_0 + \sigma_{\tilde{\jmath}} \epsilon_{\tilde{\jmath}})) \mathcal{N}(\epsilon_0|0,1) \mathcal{N}(\epsilon_{\tilde{\jmath}}|0,1)$$
$$\propto \exp\left[(\vec{\beta} \cdot \tilde{\vec{c}} + \mu_{\tilde{\jmath}} + \sigma_0 \epsilon_0 + \sigma_{\tilde{\jmath}} \epsilon_{\tilde{\jmath}})\tilde{x} - \exp(\vec{\beta} \cdot \tilde{\vec{c}} + \mu_{\tilde{\jmath}} + \sigma_0 \epsilon_0 + \sigma_{\tilde{\jmath}} \epsilon_{\tilde{\jmath}})\right]$$
$$\times \exp(-\epsilon_0^2/2) \exp(-\epsilon_{\tilde{\jmath}}^2/2),$$

omitting normalizing constants for brevity. We can transform this into a function of a single variable by noting that the sum of normal random variables $\epsilon_0$ and $\epsilon_{\tilde{\jmath}}$ is equivalent to the single normal random variable $\mu_{\tilde{\jmath}} + \sqrt{\sigma_0^2 + \sigma_{\tilde{\jmath}}^2}\mathcal{N}(0,1)$. Our integrand becomes

$$\begin{aligned} g(\epsilon) &= \text{pois}\left(\tilde{x}\middle| \exp\left[\vec{\beta}\cdot\tilde{\vec{c}} + \mu_{\tilde{\jmath}} + \epsilon\sqrt{\sigma_0^2 + \sigma_{\tilde{\jmath}}^2}\right]\right)\mathcal{N}(\epsilon|0,1) \\ &\propto \underbrace{\exp\left[\left(\vec{\beta}\cdot\tilde{\vec{c}} + \mu_{\tilde{\jmath}} + \epsilon\sqrt{\sigma_0^2 + \sigma_{\tilde{\jmath}}^2}\right)\tilde{x} - \exp\left(\vec{\beta}\cdot\tilde{\vec{c}} + \mu_{\tilde{\jmath}} + \epsilon\sqrt{\sigma_0^2 + \sigma_{\tilde{\jmath}}^2}\right)\right]}_{f(\epsilon)}\exp(-\epsilon^2/2), \end{aligned}$$

which is of the form $f(x)\exp(-x^2)$ after change-of-variable $\xi = \epsilon/\sqrt{2}$, and therefore amenable to approximation via Hermite quadrature,

$$\int_{-\infty}^{\infty} d\epsilon\, f(\epsilon)\exp(-\epsilon^2/2) = \sqrt{2}\int_{-\infty}^{\infty} d\xi\, f(\xi)\exp(-\xi^2) \approx \sqrt{2}\sum_{i=1}^{n} w_i f(x_i),$$

where $x_i$ is the $i$th root of Hermite polynomial $H_n$, and $w_i$ is the $i$th Hermite quadrature weight. Note the additional factor of $\sqrt{2}$ from transforming the volume element $d\epsilon$.

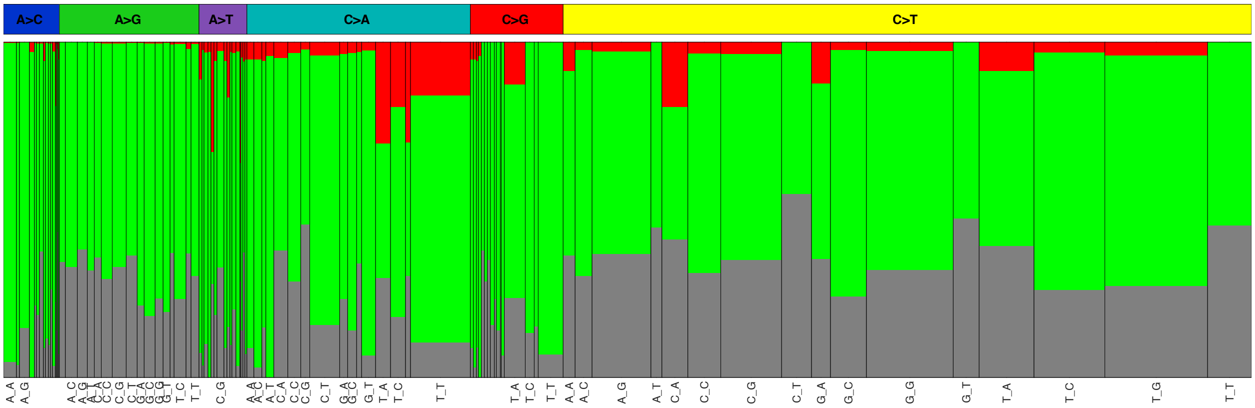


**Figure S1:** Expected protein-coding effects for each of the 96 trinucleotide substitutions. Width of each bar is proportional to observed mutation burden in our cohort for that substitution. Note that A(A→[CT])T and A(C→[AG])T substitutions can never generate synonymous (gray) mutations, and [CG](C→G)[ACG]/T(C→G)[CG]/[ACGT](C→T)T substitutions can never generate nonsense (red) mutations.

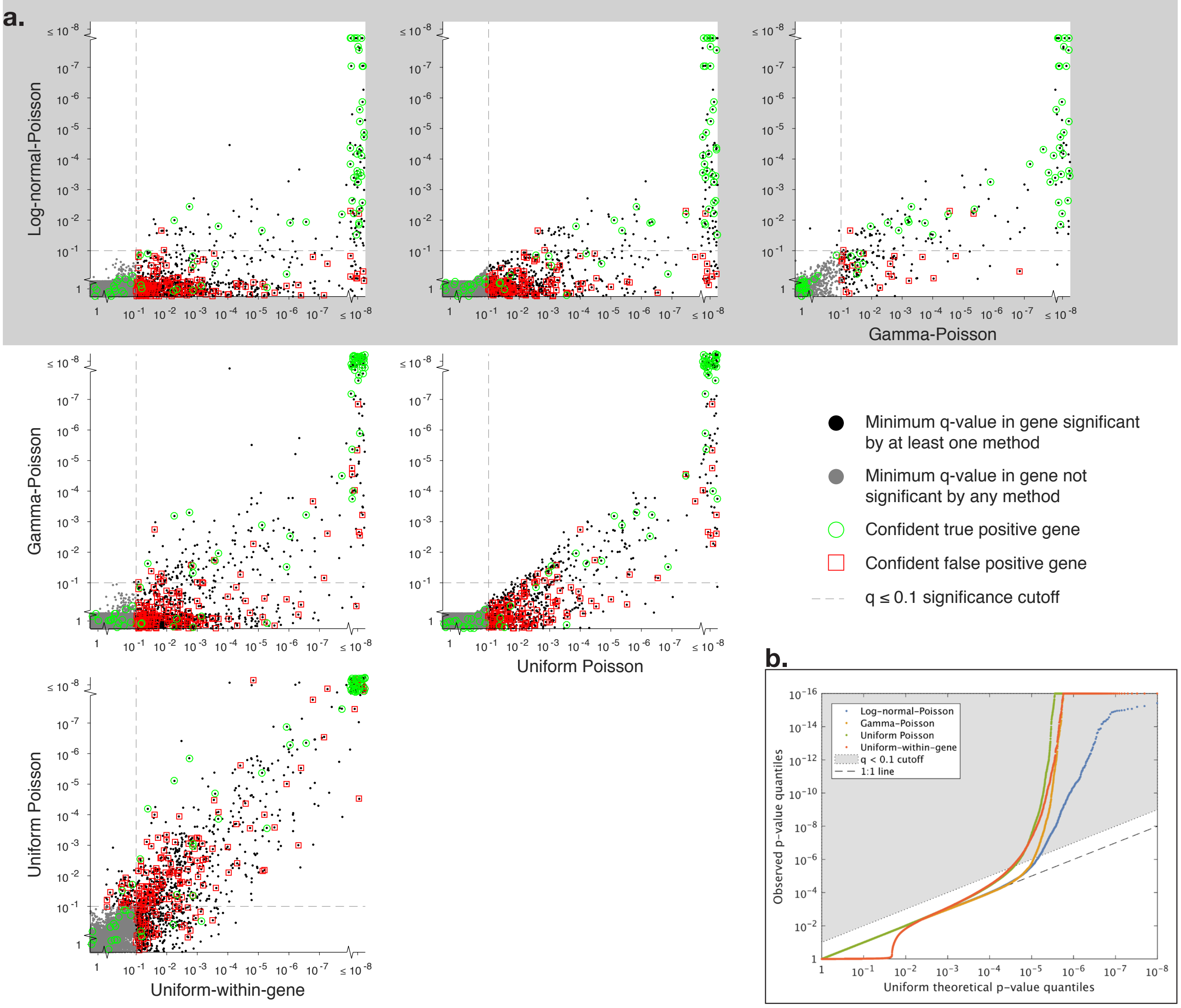


**Figure S2:** Comparison of methods' statistical calibration. **a.** Cross-method comparison of minimum $q$-values across all mutations in each gene. Genes in false-positive/true-positive truth sets are circled in red and green, respectively. $q$-value cutoff of 0.1 shown as dashed lines. For clarity, points at $q = 1$ or $q < 10^{-8}$ are randomly jittered. Comparison of Log-normal-Poisson method against other methods highlighted in gray. **b.** Quantile-Quantile (QQ) plots of each method's $p$-values. Dashed 1:1 line corresponds to uniformly distributed p-values; gray area represents region of Benjamini-Hochberg $q$-value $< 0.1$. For legibility purposes, observed $p$-values are capped at a minimum value of $10^{?16}$.

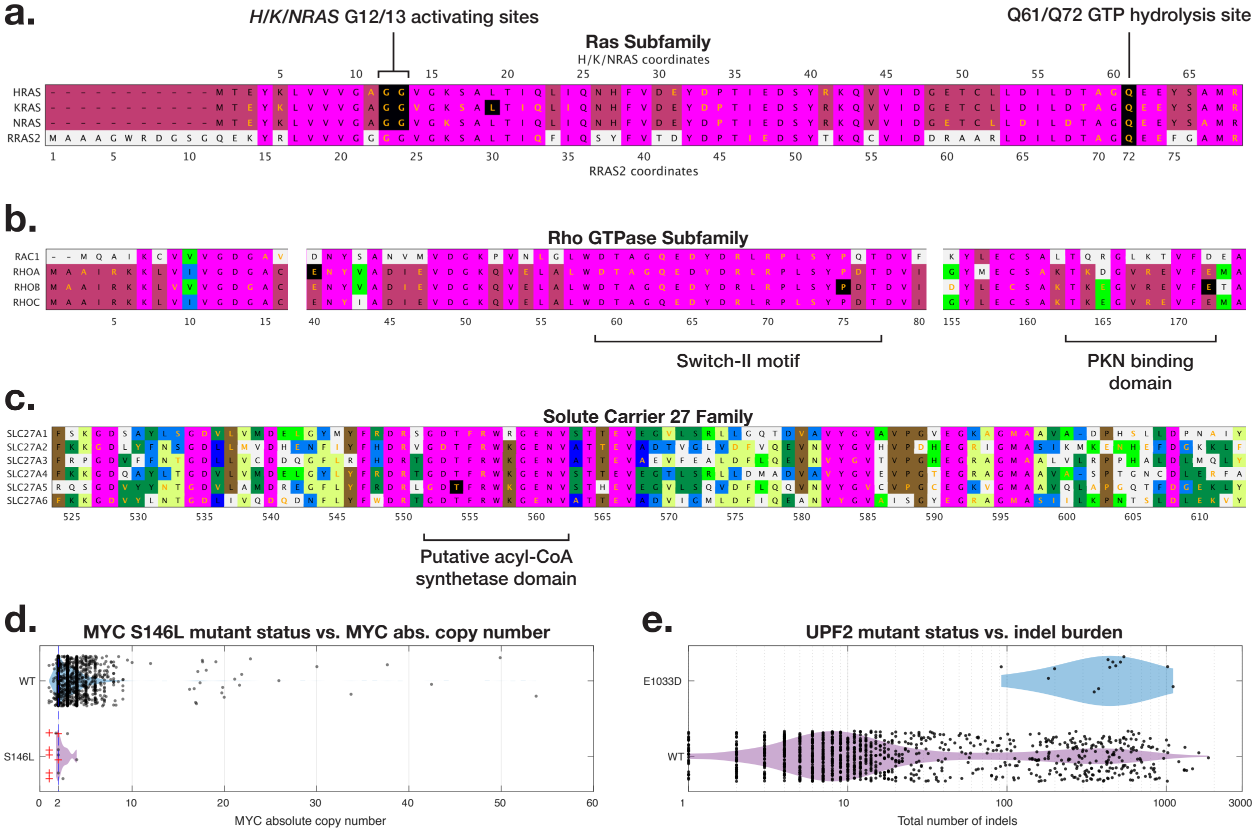


**Figure S3:** Functional exposition of novel putative driver hotspots. In all sequence alignment plots, amino acids with orange letters are mutated; amino acids with black backgrounds are significant hotspots (by the Log-normal-Poisson model). Other colors represent the number of matching amino acids across the alignment. **a.** Sequence alignment of first 68 amino acids of canonical Ras subfamily members *H/K/NRAS* and first 79 amino acids of *RRAS2*, with both *H/K/NRAS* and *RRAS2* coordinates shown to emphasize homology between hydrolysis site Q61 (and well-known driver locus) of canonical subfamily members with Q72 in *RRAS2*. **d.** Sequence alignments of three subsections of Rho GTPase subfamily members *RAC1* and *RHOA/B/C*. The first subsection comprises amino acids 1-16, in order to showcase similarity of N termini. The next subsection contains the highly conserved Switch-II motif, which contains a significant hotspot in *RHOB* and has high mutation burden across the four Rho GTPase subfamily members. The third subsection contains the PKN-binding domain, which also contains a significant hotspot in *RHOB*. **c.** Sequence alignment of solute carrier 27 family members, illustrating significant hotspot in *SLC27A5* occurring in highly conserved motif (putative acyl-CoA synthetase domain) within a larger nonconserved context. **d.** Absolute copy number at *MYC* of all wildtype patients versus S146L mutants. Red crosses show allelic copy number of somatic mutations. Ranksum of mutant vs. wildtype absolute copy number $p$-value $= 0.02$. **e.** Total somatic indel burden of *UPF2* E1033D mutants versus wildtype patients in the same cohort as the mutants. Ranksum of indel burden $p$-value $= 2.3 \times 10^{-7}$

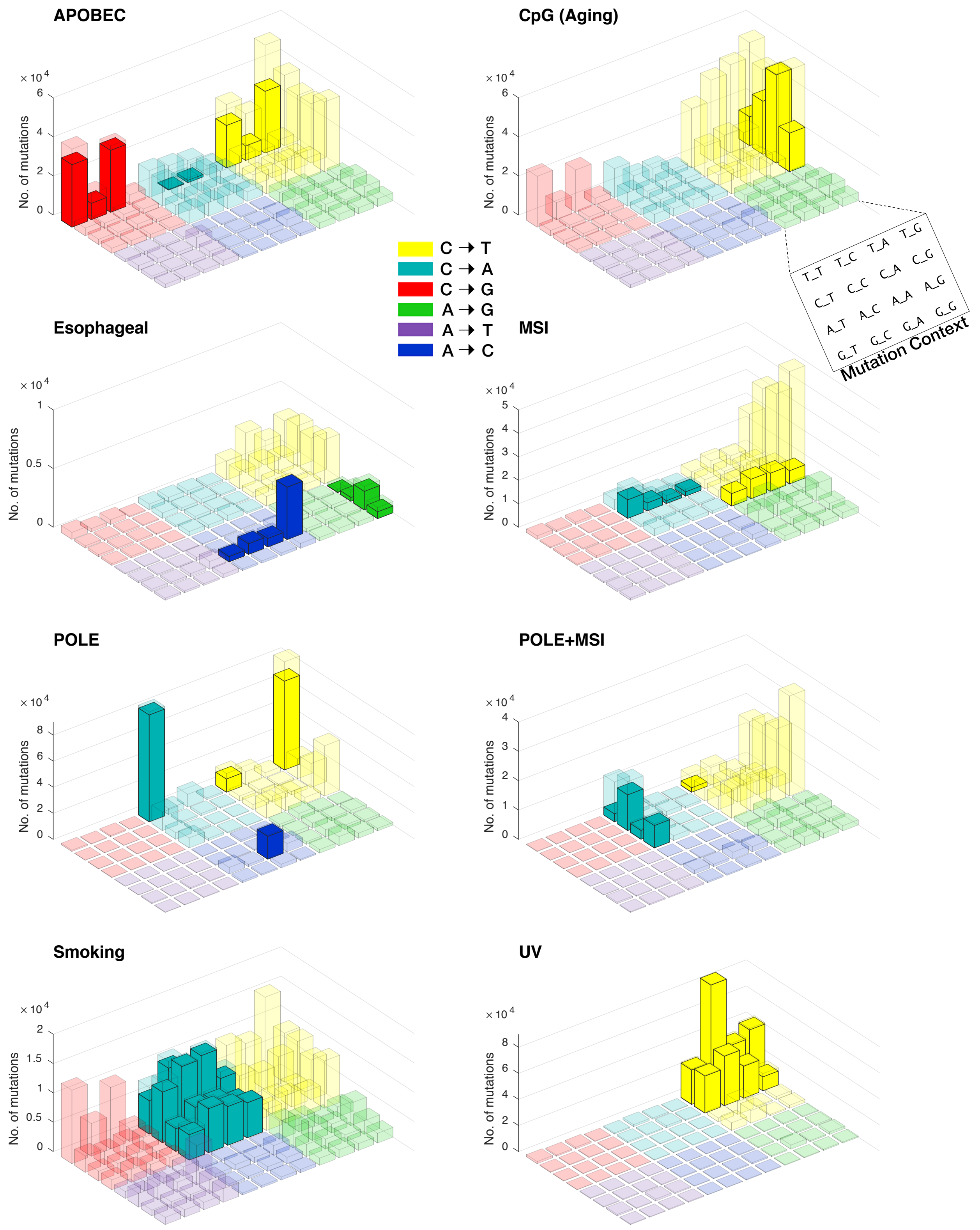


**Figure S4:** Total mutation counts of the 96 trinucleotide substitutions in each mutational process-centric subcohort. Opaque bars show counts of mutations definitively attributed to the relevant mutational process (Bayesian NMF factor assignment probability ≥0.75); transparent bars show counts of all other mutations in the process-centric subcohort not attributed to the relevant mutational process.

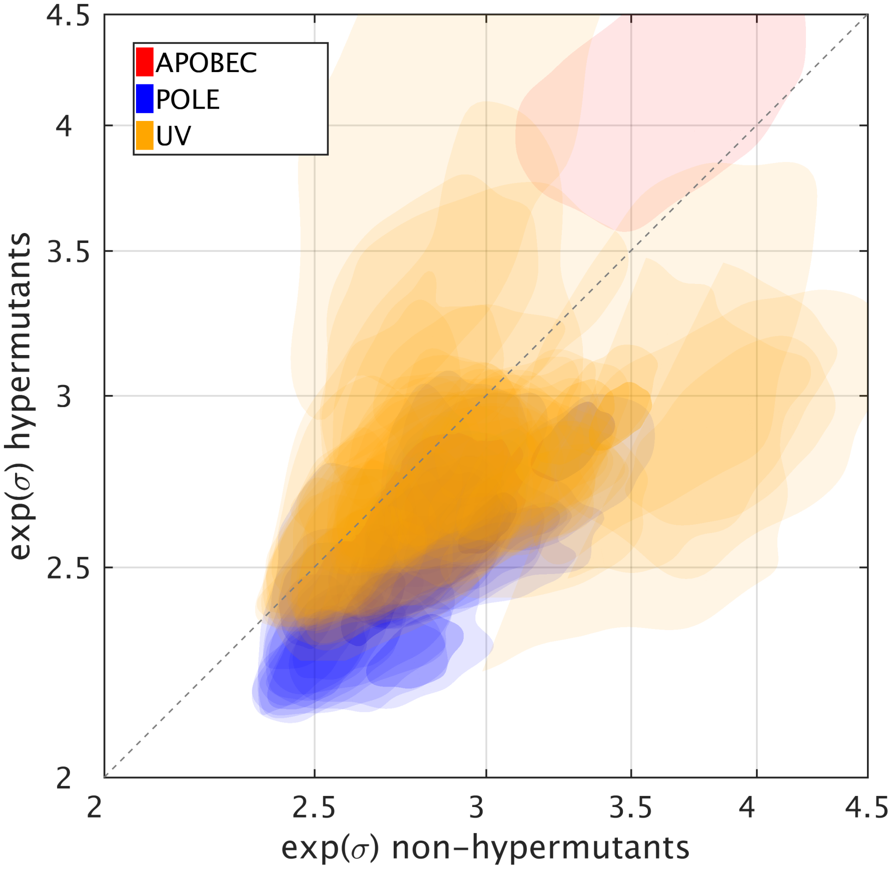


**Figure S5:** Joint posterior distributions of geometric standard deviation $e^{\sigma}$ for non-hypermutants vs. hypermutants. Each colored area is the 95% confidence region for a pentamer context associated with a given mutational process. Only mutational processes with sufficiently tight posterior densities in both the hypermutant and non-hypermutant partitions are shown.

**Description of table fields**

| | |
|---|---|
| name | Gene name (GENCODE v19) |
| neu_prob | Probability $0.8 \leq dN/dS \leq 1.1$ (false positives only) |
| dNdS_mean | Mean $dN/dS$ |
| dNdS_95CI | 95% confidence interval of $dN/dS$ distribution |
| dNdS_change | Percent change in mean $dN/dS$ when removing recurrent sites (true positives only) |
| sig_UWG<br>sig_UP<br>sig_NB<br>sig_LNP | Gene contains $\geq$1 significant hotspot ($q \leq 0.1$) according to given method |

(To view table, please see Excel workbook Supp_Table_1.xlsx)

**Table S1:** List of genes in true positive/false positive truth sets. False-positive set comprises 404 genes confidently under neutral selection (Prob[$0.8 \leq dN/dS \leq 1.1$] $\geq$ 80%); true-positive set comprises 44 known cancer genes whose $dN/dS$ drops by more than 5% after removing all sites near significance ($q \leq 0.25$) by the least conservative Uniform-within-gene model.

**Description of table fields**

| | |
|---|---|
| chr | Chromosome (23 represents X chromosome) |
| pos | Position in hg19 coordinates |
| base_change | Strand-collapsed base substitution (and flanking pentamer sequence) |
| gene | Gene name (GENCODE v19) |
| count | Total number of mutated patients |
| ttypes | Count of mutated patients' tumor types (TCGA nomenclature) |
| sig_assignments | Number of mutated patients assigned to each signature subcohort with ≥75% probability. Patients with <75% assignment probability are not counted here. |
| protein_change | Amino acid substitution |
| coding_effect | Protein coding effect: missense, nonsense, synonymous, or splice (if mutation is within 5 bp of a splice site; "splice" supersedes all other effects) |
| is_KCG | Known cancer gene status |
| p_UWG<br>q_UWG<br>p_UP<br>q_UP<br>p_NB<br>q_NB<br>p_LNP<br>q_LNP | $p/q$-values according to given method |
| APOBEC_hairpin | Mutation occurs at a palindromic sequence predicted to be targeted by APOBEC deaminase enzymes; probable false positive. |

**Table S2:** List of hotspot mutations significant ($q \leq 0.1$) by each model.

(To view table, please see Excel workbook Supp_Table_2.xlsx)

**Table S3:** Annotations for every non-known cancer gene significant by LNP model.

| Gene | Prot. Ch. | N. Mut. | Tumor types | Signatures | LNP q-value |
|---|---|---|---|---|---|
| *RXRA* | S427F | 14 | BLCA:12, LIHC:1, PAAD:1 | APOBEC:9, Misc:1 | $5.40 \times 10^{-7}$ |
| S427F mutant experimentally characterized (PMID 29143738) to activate peroxisome proliferator-activated receptors (PPAR) signalling, which stimulates downstream growth pathways. WT PPARG activation requires both heterodimerization with RXRA and a bound co-activator; S427F mutant was demonstrated to be able to activate PPARG even without this co-activator. | | | | | |
| *MB21D2* | Q311E | 19 | BLCA:3, BRCA:1, ESCA:1, HNSC:5, LUAD:3, LUSC:6 | APOBEC:19 | $3.45 \times 10^{-6}$ |
| Located in palindromic sequence context amenable to APOBEC mutagenesis. Conserved in vertebrates. No PubMed papers dealing specifically with this gene, but physical interactions with several ciliary or centrosome proteins (AHI1, CENPJ, CEP104, CEP170, CEP89, CNTROB, MKS1, RAB3IP/RABIN8, STIL and TMEM216), or with 6 TRAPP complex subunits detected in high-throughput studies suggest potential functions in ciliary functions and/or protein trafficking. | | | | | |
| *ERCC2* | N238S | 9 | BLCA:9 | Misc:9 | $1.93 \times 10^{-5}$ |
| ERCC2 encodes a DNA helicase essential for nucleotide excision repair. N238 falls in its helicase ATP-binding domain, whose mutation is associated with a distinct mutational signature (PMID 27111033), suggesting that a damage checkpoint is being evaded, which could speed cell proliferation or evade damage-induced apoptotic responses. | | | | | |
| *SLC27A5* | T554I | 13 | SKCM:13 | UV:13 | $2.99 \times 10^{-5}$ |
| "Activation" of fatty acids by thioesterification to acyl-coenzyme A is required for their participation in both anabolic and catabolic pathways. There are 26 acyl-coA synthetase genes in the human genome, among which the six members of the SLC27A gene family (SLC27A1-6) specifically activate long chain fatty acids. SLC27A5, also known as FATP5, is expressed in liver and plays a role in the uptake of long-chain fatty acids from the blood stream (PMID 16618416), as well as in bile acid metabolism by serving as a bile acid coA ligase (PMID 16618417). Amino acid T554 maps to a protein segment (GDTFRWKGENV) that is highly conserved not only between SLC27A5 orthologs, but also between paralogs in several species. | | | | | |
| *OXA1L* | L57F | 9 | HNSC:1, SKCM:8 | UV:9 | $3.50 \times 10^{-5}$ |
| Budding yeast Oxa1p is a protein involved in the insertion of nuclear and mitochondrial encoded proteins into the mitochondrial inner membrane. Association of the mitochondrial ribosome with Oxa1 requires its C-terminal region. Some human OXA1L isoforms have a 60 AA N-terminal extension not present in any chimp, mouse, dog, or zebrafish orthologs. However, orthologs from other species do include regions corresponding to this N-terminal segment. | | | | | |
| *BCL2L12* | F17F | 13 | SARC:1, SKCM:12 | UV:13 | $3.63 \times 10^{-5}$ |
| F17F previously identified as a synonymous mutation in melanoma that increased BCL2L12 mRNA and protein because of differential targeting of WT and mutant BCL2L12 mRNA by hsa-miR-671-5p (Seed sequence GGAAGCC, PMID: 23901115, see also 28026089). Mutation looks like rs267605591 synonymous C/T SNP. BCL2L12 interacts with CASP7 and controls its maturation (PMID: 17210792, 18669646). Knockdown leads to cisplatin resistance in breast cancer cell line (PMID 18930135). Endogenous BCL2L12 interacts with TP53 and inhibits its function (PMID: 20837658). Gene overlaps with that coding for IRF3. | | | | | |
| *SOX17* | S403I | 7 | UCEC:7 | Misc:1 | $4.79 \times 10^{-5}$ |

**Table S3:** Annotations for every non-known cancer gene significant by LNP model.

<table>
<tr><th>Gene</th><th>Prot. Ch.</th><th>N. Mut.</th><th>Tumor types</th><th>Signatures</th><th>LNP q-value</th></tr>
<tr><td colspan="6">Member of the SOX family of DNA binding proteins involved in transcriptional regulation. Closest relatives are SOX7 and SOX18. S403I hotspot maps near the C-terminus of the 414 residue SOX17 protein, in a segment that is well conserved between SOX17 orthologs. Sox17 orthologs in mice (PMID 11973269), Zebrafish (PMID 10531029), Xenopus (PMID 9363948) and man (PMID 18682240) function in endoderm formation. Human Sox17 also has a role in the maintenance of fetal and neonatal hematopoietic stem cells (PMIDs 17655922, 21828271) and in the specification of primordial germ cell fate (PMID 25543152). A C-terminal transactivation domain of Xenopus Sox17 (roughly corresponding to amino acids 350-370 of human SOX17) physically interacts with beta-catenin (PMID 15163629). Human SOX17 also interacts with beta-catenin (PMID 17875931). SOX17 is epigenetically inactivated in several cancers, including in colorectal (PMID 18413743) and breast cancer (PMID 19301122), and has been implicated as a tumor suppressor in endometrial cancer (PMID 27738313). However, frequent SOX17 frameshift mutations in endometrial cancer are functionally distinct from recurrent A96G or S403I missense mutations in that the latter do not appear to affect transcription activity (PMID 28978154).</td></tr>
<tr><td>PDE3A</td><td>L275L</td><td>12</td><td>BLCA:5, CESC:1, ESCA:1, HNSC:2, LUAD:2, LUSC:1</td><td>APOBEC:11</td><td>$6.02 \times 10^{-5}$</td></tr>
<tr><td colspan="6">Located in palindromic sequence context amenable to APOBEC mutagenesis. cGMP-inhibited cyclic nucleotide phosphodiesterase best known for its roles in cardiac function, smooth muscle contraction, and oocyte maturation. Silent hotspot overlaps partial antisense lncRNA RP11-284H19.1, which comprises two exons. First exon of lncRNA overlaps portion of first exon of PDE3A; 3' end of lncRNA is 8.5kb upstream of PDE3A TSS. Nothing in literature is known about RP11-284H19.1. Nucleic acid itself is nonconserved across vertebrates (as one would expect, since leucine has fourfold degeneracy at third codon position). Amino acid is mostly conserved across mammals. Serine/arginine substitutions suggest nonessentiality of amino acid, since L/S/R are similar codon sequence but different in amino acid properties.</td></tr>
<tr><td>SPTLC3</td><td>R97K</td><td>13</td><td>SKCM:12, UCEC:1</td><td>UV:12</td><td>$7.08 \times 10^{-5}$</td></tr>
<tr><td colspan="6">Widely conserved subunit of serine palmitoyltransferase (together with SPTLC1, SPTLC2, SPTSSA and SPTSSB). Implicated in the generation of C-16 backbone containing sphingolipids.</td></tr>
<tr><td>PCBP1</td><td>L100Q</td><td>11</td><td>COAD:6, LIHC:1, READ:2, STAD:1, THCA:1</td><td>Misc:10</td><td>$9.16 \times 10^{-5}$</td></tr>
<tr><td colspan="6">Poly(RC) Binding Protein 1 binds to poly-C regions of mRNAs, thus blocking their translation. L100 lies in the protein's KH domain, responsible for binding RNA. Although an exhaustive list of PCBP1 targets is unknown, it has been experimentally validated to suppress translation of metastasis-associated PRL-3 phosphatase (PMID 20609352), with knockdown "caus[ing] upregulation of PRL-3 protein levels, activation of AKT, and promotion of tumorigenesis."</td></tr>
<tr><td>STK19</td><td>D89N</td><td>8</td><td>SKCM:8</td><td>UV:8</td><td>$1.46 \times 10^{-4}$</td></tr>
<tr><td colspan="6">Nuclear protein serine/threonine kinase. Gene maps to the human major histocompatibility complex. STK19 physically interacts with CSB/ERCC6 and cells depleted for STK19 were deficient in the recovery of transcription after DNA damage (PMID 27184836).</td></tr>
<tr><td>C3orf70</td><td>S6L</td><td>16</td><td>BLCA:8, CESC:3, ESCA:2, LUSC:3</td><td></td><td>$1.93 \times 10^{-4}$</td></tr>
<tr><td colspan="6">Located in palindromic sequence context amenable to APOBEC mutagenesis. Conserved in vertebrates, only physical interactor identified in high throughput affinity capture/MS screen is RBM3.</td></tr>
<tr><td>EEF1A1</td><td>T432I; T432S</td><td>6; 5</td><td>GBM:1, HNSC:1, LIHC:4; HNSC:1, LIHC:4</td><td>Misc:3; Misc:2</td><td>$2.20 \times 10^{-4}$; $1.24 \times 10^{-2}$</td></tr>
</table>

**Table S3:** Annotations for every non-known cancer gene significant by LNP model.

<table>
<tr><th>Gene</th><th>Prot. Ch.</th><th>N. Mut.</th><th>Tumor types</th><th>Signatures</th><th>LNP q-value</th></tr>
<tr><td colspan="6">Eukaryotic translation factor 1 alpha 1, 93% identical to EEF1A2. T432 is conserved in EEF1A2 and in 38-40% identical HBS1 and ERF3A. Around 40 human EEF1A1 pseudogenes are known to exist. Reads supporting hotspot mutation were confirmed to map uniquely to EEF1A1. Eukaryotic elongation factor-1 consists of the GTP-binding EEF1A1/2 subunit together with EEF1B2, EEF1G and EEF1D. The complex transfers aminoacyl-tRNA to the ribosome. Aminoacyl-tRNA is released upon GTP hydrolysis and EF1-alpha-GDP is recycled to EF1-alpha-GTP by the beta, gamma and delta subunits. Beyond translation initiation, EEF1A1 has been implicated in the activation of heat shock factor 1 (PMID 16554823), and as a subunit of the BAT complex (with PCBP1) in the TGF-beta regulated translation of EMT transcripts DAB2 and ILEI (PMID 21329880). EEF1A1 physically interacts with $\sim$420 other proteins and testing whether the T432I mutation specifically affects the interaction with some of these might provide clues as to whether and how this mutation contributes to cancer. In addition, EEF1A1 is a pan-essential gene according to the Achilles genome-wide CRISPR knockout screen.</td></tr>
<tr><td>RQCD1</td><td>P131L</td><td>12</td><td>SKCM:12</td><td>UV:12</td><td>$2.79 \times 10^{-4}$</td></tr>
<tr><td colspan="6">Also known as CNOT9. Evolutionary conserved subunit of the CCR4-NOT complex that functions as a key regulator of eukaryotic gene expression (reviewed in PMID 26821858). CCR4-NOT complex regulates gene expression at multiple levels, including mRNA deadenylation and degradation, and miRNA targeting and gene silencing (PMIDs 24768538 and 24768540). Two tryptophan binding pockets of CNOT9 implicated in binding miRNA processing protein GW182/TNRC6, are also essential for binding tristetraprolin/ZFP36, a protein involved in mRNA deadenylation. However, a BTG2-Caf1-Ccr4-CNOT9-CNOT1 pentameric complex containing the melanoma-associated CNOT9 P131L variant supports mRNA deadenylation similar to a complex containing the wild type CNOT9 protein (PMID 30309886).</td></tr>
<tr><td>SUCO</td><td>M1I</td><td>9</td><td>LUSC:1, SKCM:8</td><td>UV:9</td><td>$3.49 \times 10^{-4}$</td></tr>
<tr><td colspan="6">Interaction between budding yeast SUCO ortholog SLP1 and EMP65 (orthologous to human TAPT1) implicated in ER membrane protein folding. SUCO/TAPT1 interaction is conserved in man. Mouse Suco also known as Osteopotentia (ost) regulates osteoblast maturation, bone formation and skeletal integrity (PMID 20440000).</td></tr>
<tr><td>C10orf76</td><td>Q267K</td><td>5</td><td>BRCA:1, ESCA:3, SARC:1</td><td>MSI:1, Misc:3</td><td>$5.37 \times 10^{-4}$</td></tr>
<tr><td colspan="6">Conserved in vertebrates, Drosophila and S. pombe. siRNA-mediated suppression alters the expression of a limited set of genes, including NSBP1 and NFYB (PMID 17959645). C10orf76 and PI4KB (critical for the maintenance of the Golgi and trans Golgi network) physically interact and were both synthetic lethal with PTAR1.</td></tr>
<tr><td>KLF5</td><td>E419Q; E419K</td><td>7; 7</td><td>LUAD:1, LUSC:6; BLCA:1, CESC:5, LUAD:1</td><td>APOBEC:7; APOBEC:7</td><td>$5.65 \times 10^{-4}$; $6.14 \times 10^{-3}$</td></tr>
<tr><td colspan="6">Krüppel-like factor 5 (KLF5, a.k.a. BTEB2) is a C2H2 zinc finger putative transcription factor expressed in the intestinal tract among other places. Missense mutations affecting E419 in the second of three C2H2 zinc finger motifs affect the DNA binding specificity of KLF5 (PMID 28963353). The authors of this study noted that the E419K mutation occurs predominantly in cervical squamous cell carcinomas whereas the E419Q mutation is found only in lung cancers.</td></tr>
<tr><td>AHR</td><td>Q383H</td><td>9</td><td>BLCA:7, KIRP:1, LUAD:1</td><td>APOBEC:8</td><td>$6.82 \times 10^{-4}$</td></tr>
<tr><td colspan="6">Aryl hydrocarbon (dioxin) receptor implicated as a ligand activated transcription factor AND as a ligand-dependent E3 ubiquitin ligase that targets sex hormone receptors (AR and ESR1) for degradation (PMID 17392787). Ahr(-/-) mice lack a teratogenic response to 2,3,7,8-tetrachlorodibenzo-p-dioxin, and also lack Benzo[α]pyrene carcinogenicity (PMIDs 9427285, 10639156). Among other findings, murine AHR suppresses intestinal carcinogenesis in Apc-min/+ mice (PMID 19651607) and regulates both T(reg) and T(H)17 cell differentiation in a ligand-specific fashion (PMIDs 18362915, 18362914). Q383H mutation maps at C-terminal edge of PAS domain.</td></tr>
<tr><td>TPTE2</td><td>E51G; S40R</td><td>9; 8</td><td>COAD:7, STAD:2; COAD:6, STAD:2</td><td>Misc:6; Misc:7</td><td>$6.89 \times 10^{-4}$; $8.45 \times 10^{-2}$</td></tr>
</table>

**Table S3:** Annotations for every non-known cancer gene significant by LNP model.

| Gene | Prot. Ch. | N. Mut. | Tumor types | Signatures | LNP q-value |
|---|---|---|---|---|---|
| Encodes transmembrane phosphoinositide 3-phosphatase and tensin homolog, also known as TPIP. Human genome includes one closely related paralog (TPTE, 88% identical) and seven peseudogenes; we verified that reads supporting this hotspot mutation mapped uniquely. TPIP has PtdIns(3,4,5) P (3) 3-phosphatase activity and is localized to the endoplasmic reticulum (PMID 15046604). A SNP highly significantly associated with HCC ($p = 1.74 \times 10^{-12}$) lies in TPTE2 (PMID 21105107). Over-expression of TPIP-C2 isoform in HeLa cells strongly (up to 85%) inhibited cell growth/proliferation and caused apoptosis in a caspase 3-dependent manner (PMID 22164291). A Ciona intestinalis ortholog known as Ci-VSP includes four N-terminal TM domains that serve as a voltage sensor that directly translates changes in membrane potential into the turnover of phosphoinositides (PMID 15902207). The corresponding region of human TPTE2 encompasses AA 53-203 of isoform NP_954863.2 (PMID 22396523). Perhaps the E51G mutation affects this function and alters the catalytic activity of TPTE2. | | | | | |
| *NUP93* | Q15* | 6 | BLCA:3, THCA:3 | Misc:4 | $7.04 \times 10^{-4}$ |
| Nuclear pore protein 93. PMID 24572986 is a review of NUP93's role in pore assembly and nucleocytoplasmic transport. Mutations in steroid resistant nephrotic syndrome (26878725). Evolutionary conserved role in the nuclear trafficking of SMAD proteins (26878725). Loss of function mutation likely to impair nucleocytoplasmic transport of SMAD family members, and likely various other proteins. | | | | | |
| *LPAR6* | F316F | 8 | BLCA:6, BRCA:1, LUAD:1 | APOBEC:8 | $8.47 \times 10^{-4}$ |
| Located in palindromic sequence context amenable to APOBEC mutagenesis. Receptor for lysophosphatidic acid (LPA). Recessive truncating and missense mutations cause woolly hair/hypotrichosis. LPA signaling through LPAR4 and LPAR6 negatively regulates motility of colon cancer cells (PMID: 27993681). LPAR6 knockdown reduced HCC cell and tumor growth (PMID 25589345) and abolished metastasis of androgen-independent PCa cells (PMID 25285406). | | | | | |
| *AGAP5* | H109R | 9 | STAD:8, UCEC:1 | MSI:2 | $1.03 \times 10^{-3}$ |
| Member of a family of nine closely related ArfGAPs. Family also includes seven pseudogenes. No publications dealing specifically with AGAP5. AGAP1, AGAP2 and AGAP3 include N-terminal GTPase domain followed by PH, ARFGAP and ANK motifs. AGAP5 and the remaining AGAPs lack the N-terminal GTPase-like sequence. The AGAP5 H109R mutation does not affect any known motif and given high level of sequence identity between AGAP4, AGAP5, AGAP6, AGAP9 and LOC107984026 (all localized on chromosome 10 and not conserved beyond primates), it is not clear what this AGAP5 missense mutation would do from sequence analysis alone. | | | | | |
| *ZBTB7B* | K388N | 4 | BLCA:2, KIRC:1, UCEC:1 | Misc:3 | $1.94 \times 10^{-3}$ |
| Zbtb7b is a transcription factor involved in CD4+ T lymphocyte development (PMIDs 15729333, 15750595, 16448546) and that has additional roles in the activation of the thermogenic gene program in adipocytes (PMID 28784777) and in mammary gland lactation (PMID 29420538). Transgenic expression leads to high incidence of T-cell lymphomas in mice (PMID 26056302). Mutated residue K388 is a putative DNA binding residue in the 2nd Zn-finger of Zbtb7b. | | | | | |
| *ZNF623* | T500K | 6 | HNSC:5, LUSC:1 | smoking:2 | $2.02 \times 10^{-3}$ |
| Protein that includes 14 H2C2 zinc fingers. Conserved in mammals but not beyond. Putative transcription factor. | | | | | |
| *ARL16* | G6R | 5 | SKCM:5 | UV:5 | $2.09 \times 10^{-3}$ |
| Binds to a C-terminal domain and functions as an inhibitor of RIG-I (a.k.a DDX18), which recognizes virus produced RNAs and stimulates interferon production. | | | | | |
| *ACTB* | G158R | 7 | BLCA:5, CESC:1, UCS:1 | CpG:2 | $3.58 \times 10^{-3}$ |
| Gene encodes beta-actin. Missense mutation causes Baraitser-Winter syndrome (PMID 22366783). G158R mutation in cancer may reflects beta-actin's role as a subunit of SWI/SNF and other chromatin remodeling complexes (PMID 12045110,12805231,18442483), but this is pure speculation at this point. | | | | | |
| *DCAF13* | T118T | 7 | BLCA:1, BRCA:1, COAD:1, ESCA:1, STAD:3 | Misc:2 | $3.97 \times 10^{-3}$ |

**Table S3:** Annotations for every non-known cancer gene significant by LNP model.

| Gene | Prot. Ch. | N. Mut. | Tumor types | Signatures | LNP q-value |
|---|---|---|---|---|---|
| Localized at recurrently focally amplified locus in several cancers. Budding yeast ortho-log SOF1 is subunit of SSU processosome, involved in ribosome biogenesis (PMID 12068309). Human DCAF13 interacts with several SSU processosome subunits. | | | | | |
| *LTN1* | S19F | 6 | LUSC:1, SKCM:5 | UV:6 | $4.29 \times 10^{-3}$ |
| Listerin E3 ubiquitin ligase 1. Budding yeast ortholog is a subunit of the RQC complex involved in protein quality control (PMID 20835226). Interaction between budding yeast RQC complex subunits RKR1 (LTN1 ortholog) and RQC2 (NEMF ortholog) is conserved in man. The LTN1 S19F mutation may allow persistence of proteins that would otherwise be degraded by RQC complex mediated quality control. The N-terminal region of LTN1 where this mutation occurs is conserved in primates/monkeys but not in other mammals. | | | | | |
| *MYO10* | G1223R | 7 | COAD:1, STAD:2, UCEC:4 | MSI:2 | $4.96 \times 10^{-3}$ |
| Mutation alters conserved residue in the loop between the beta-1 and beta-2 sheets of the 2nd of three MYO10 PH domains (see alignment and discussion in PMID 9736615). A MYO10 R1231C mutation affecting a conserved residue in the beta-2 sheet of the same PH domain (a residue required for high-affinity PIP3 binding) abolishes MYO10's ability to inhibit phagocytosis (PMID 12055636). Much evidence also implicates MYO10 in filopodia formation and behavior (e.g. PMIDs 16894163, 22124140, 28289096, 29057977). Thus, mutant protein may have altered subcellular localization, altered interaction with phospholipids or other proteins, and impaired function. | | | | | |
| *ITGB1* | L378I | 5 | STAD:5 | MSI:4 | $4.99 \times 10^{-3}$ |
| Integrin subunit beta-1. Integrins are heterodimers of alpha and beta subunits that function as adhesion (cell-cell or cell-matrix) receptors. L378I mutation in putative vWFA (also known as an "I"-like) domain (PMIDs 9009218, 10779511) may affect dimerization and/or interaction with adhesion substrates. | | | | | |
| *ANKRD40* | D99E | 6 | COAD:1, STAD:4, UCEC:1 | Misc:5 | $5.48 \times 10^{-3}$ |
| Orthologs exist in vertebrates. D99E mutation involves 2nd D residue in the following sequence: EEEDDDDDDDD, which is not well conserved. No publications. Only physical interactor detected by more than one lab is adenosylhomocysteinase (AHCY). | | | | | |
| *MYCN* | P44L | 7 | SKCM:1, UCEC:6 | UV:1, Misc:3 | $6.45 \times 10^{-3}$ |
| Member of the MYC family of transcription factors. NMYC gene is amplified and over-expressed in neuroblastoma, retinoblastoma, and small cell lung cancer. Proline-44 maps closely upstream of the conserved Myc-box-1 (K51-R65), a segment containing two phosphorylation (Thr58 and Ser 62) sites whose modification controls NMYC protein stability (PMID 14660435). | | | | | |
| *KRT15* | V205I | 15 | COAD:2, GBM:3, HNSC:1, KIRC:1, LGG:2, LIHC:1, PAAD:1, PRAD:1, SARC:1, SKCM:1, UCEC:1 | CpG:12, Misc:1 | $7.26 \times 10^{-3}$ |
| Cytokeratin 15 expressed in various epithelial tumors and marks hair-follicle epithelial stem cells (PMIDs 9763512, 14708593). Valine-205 is highly conserved in other cytokeratins. Mutation may affect cytoskeletal integrity and perhaps the behavior of K15 expressing stem and/or progenitor cells. | | | | | |
| *RGP1* | R17K | 5 | SARC:1, SKCM:4 | UV:5 | $7.71 \times 10^{-3}$ |
| Budding yeast RGP1 is a subunit of a dimeric Golgi membrane exchange factor (Ric1p-Rgp1p) that catalyzes nucleotide exchange on Ypt6p. Human RIC1-RGP1 complex is a guanine nucleotide exchange factor for the late Golgi Rab6A GTPase and an effector of the medial Golgi Rab33B GTPase (PMID 23091056). | | | | | |
| *FAM221B* | M1I | 7 | LUSC:1, SKCM:6 | UV:7 | $9.14 \times 10^{-3}$ |
| Conserved in mammals but not beyond. No publications on FAM221A or FAM221B. | | | | | |
| *SMTNL2* | | 13 | BRCA:6, UCEC:7 | MSI:1, Misc:8 | $1.07 \times 10^{-2}$ |

**Table S3:** Annotations for every non-known cancer gene significant by LNP model.

<table>
<tr><th>Gene</th><th>Prot. Ch.</th><th>N. Mut.</th><th>Tumor types</th><th>Signatures</th><th>LNP q-value</th></tr>
<tr><td colspan="6">Smoothelins (SMTN, SMTNL1 and SMTNL2) are markers of smooth muscle cells (SMCs). Smoothelin itself comes in two forms, Smoothelin-A (59 kDa), found in visceral SMCs, and -B (110 kDA) found in vascular SMCs. SMTN deficient mice show decreased SMC contractility (PMID 16285958). SMTNL1 has been implicated in cGMP/cAMP-mediated adaptations to exercise through mechanisms involving direct modulation of contractile activity (PMID 18310078), and as a bifunctional regulator of the progesterone receptor during pregnancy. The latter function involves a direct physical interaction between the PGR and SMTNL1 in uterine SMCs that leads to higher levels of PGR expression (PMID 21771785). SMTNL2, expressed in many mammalian tissues, with a notably high expression in skeletal muscle. binds with high affinity to JNK1-3 and ERK2 (PMID 23981301). Another clue to its function is that SMTNL2 was identified in a siRNA screen as one of a small number of regulators of epithelial morphogenesis (PMID 22820376).</td></tr>
<tr><td>GPR125</td><td>G42C</td><td>5</td><td>BRCA:1, CESC:1, SARC:3</td><td></td><td>$1.09 \times 10^{-2}$</td></tr>
<tr><td colspan="6">Surface marker of human spermatogonial stem cells (PMID 17882221). Interacts with DLG1 at cell membranes (PMID 15021905). Related GPR124/ADGRA2 controls angiogenesis in normal and tumor tissues (PMIDs 21071672,21282641, 21421844), plays a role in Wnt7 signaling (PMID 25558062), and is essential for the integrity of the blood/brain barrier (28288111).</td></tr>
<tr><td>FGFR1</td><td>N546K</td><td>5</td><td>LGG:1, PCPG:2, SARC:1, UCEC:1</td><td>Misc:5</td><td>$1.22 \times 10^{-2}$</td></tr>
<tr><td colspan="6">Fibroblast growth factor receptor 1 (FGFR1), a transmembrane tyrosine kinase that is frequently amplified in lung and breast cancer, and that is also activated by several cancer-associated chromosomal translocations. Residue N546 lies adjacent to residue I545 important for ATP binding and corresponds exactly to N550 of FGFR2, which is frequently mutated to either N550K or N550H in endometrial tumors (PMID 22383975). The FGFR1 N546K mutation is also frequently found in cancer and is known to activate FGFR1 signal transduction (e.g. PMID 19224897).</td></tr>
<tr><td>GNA13</td><td>R200G</td><td>4</td><td>BLCA:4</td><td>Misc:4</td><td>$1.57 \times 10^{-2}$</td></tr>
<tr><td colspan="6">Heterotrimeric G-protein alpha subunit that promotes Rho GTPase signaling by binding to and stimulating the activity of the related Rho exchange factors ARHGEF1/p115 RhoGEF (PMID 9641916), ARHGEF11/PDZ-RhoGEF (PMID 10026210) and ARHGEF12/LARG (PMID 18084302). Among its roles, GNA13 is required for growth factor stimulated cell migration (PMID 16740474). Arginine residue 200 is conserved among heterotrimeric G-protein alpha subunits, but maps away from Thr274 and Asn278 in the α 3 helix of Gα13 whose mutation impairs the ability of Gα13 to stimulate Rho activation in cells (PMID 21507947).</td></tr>
<tr><td>PRKCI</td><td>Y265*</td><td>15</td><td>COAD:15</td><td>Misc:12</td><td>$1.64 \times 10^{-2}$</td></tr>
<tr><td colspan="6">Possible recurrent sequencing artifact: reads supporting this variant map perfectly (with microsatellite deletion) to a paralogous region on chrX. In addition, many reads supporting variant have discordant mates also mapping the same region on chrX. The human genome includes two genes for atypical protein kinase C paralogs, PKCiota/lambda (PRKCI) and PKC-zeta (PRKCZ). Subunit of the conserved PAR-6/aPKC/PAR-3 complex that regulates epithelial cell polarity. Suggesting a positive role in carcinogenesis, gene is amplified in multiple cancers (PMID 17570678) and expression of kinase-dead PKC-iota in Ras-transformed rat intestinal epithelial cells blocked Ras-mediated Rac1 activation, cellular invasion, and anchorage-independent growth (PMID 15024028). Also implicated as an oncogene in human NSCLC (PMID 16204062) and ovarian cancer (PMID 16116079). Unclear what a truncating mutation would do to promote carcinogenesis, but perhaps this is a late step leading to loss of epithelial integrity.</td></tr>
<tr><td>SOS1</td><td>N233Y</td><td>9</td><td>LUAD:3, LUSC:1, UCEC:5</td><td>smoking:3, Misc:5</td><td>$1.65 \times 10^{-2}$</td></tr>
<tr><td colspan="6">Son-of-sevenless 1 (SOS1) is a dual guanine nucleotide exchange factor for Ras and Rac and has been implicated in integrating signaling by these GTPases PMID 9438849). The N223Y mutation maps to the RhoGEF domain. Missense mutations affecting three residues (Thr264, Met267 and Thr376) in the RhoGEF domain of SOS2 that activate the protein by interfering with its auto-inhibition have been found in Noonan syndrome (PMID 26173643). SOS1 is also mutated in Noonan syndrome, and multiple cancer-associated SOS1 missense mutations mapping throughout the entire protein are listed in COSMIC, but N233Y does not appear to be among the previously identified changes.</td></tr>
<tr><td>OLFML3</td><td>S4S</td><td>4</td><td>LUSC:3, UCEC:1</td><td></td><td>$1.70 \times 10^{-2}$</td></tr>
<tr><td colspan="6">Olfactomedin-like-3, a secreted glycoprotein (PMID 15280020). Blockade of Olfml3 by anti-Olfml3 antibodies is highly effective in reducing tumor vascularization, pericyte coverage, and tumor growth (PMID 23002094).</td></tr>
</table>

**Table S3:** Annotations for every non-known cancer gene significant by LNP model.

<table>
<tr><th>Gene</th><th>Prot. Ch.</th><th>N. Mut.</th><th>Tumor types</th><th>Signatures</th><th>LNP q-value</th></tr>
<tr><td>DDX17</td><td>H504R</td><td>4</td><td>SKCM:4</td><td>Misc:4</td><td>$1.73 \times 10^{-2}$</td></tr>
<tr><td colspan="6">U1snRNP-associated RNA helicase also known as p72 (PMID 12193588) that forms a heterodimer with paralog DDX5 (p68; 90% identical in central region, PMID 12595555) and that has been implicated in various processes, including RNA unwinding/annealing (PMID 11353078), alternative splicing (PMIDs 12138182,23022728,24275493), and miRNA maturation (PMIDs 25126784,26947125). Both DDX5 and DDX17 also interact with HDAC1 and repress transcription in a promoter-specific manner (PMID 15298701). Other proteins that interact with both DDX5 and DDX17 (among many others) are p53 (PMID 15660129), and ESR1 (PMIDs 19718048,19995069). Both proteins also interact with UPF3B and function in nonsense-mediated mRNA decay (PMID 23788676). Potential roles in cancer are reviewed in PMIDs 21345143, 23523990.</td></tr>
<tr><td>WDR89</td><td>L142W</td><td>7</td><td>MESO:7</td><td>Misc:2</td><td>$1.82 \times 10^{-2}$</td></tr>
<tr><td colspan="6">Mutation occurs exclusively in mesotheliomas, which is evidence of positive selection. However, closer inspection of the variant revealed that supporting reads map with lower edit distance (3 vs. 0) elsewhere in the genome. Unclear why a mapping issue would only occur in a single tumor type processed with identical downstream pipelines. WDR89 (WD repeat containing protein 89) is conserved in vertebrates, Drosophila, budding and fission yeast (among many other species). Residue 142 in 2nd WD repeat is either an L (mammals) or an F (Xenopus, Zebrafish, Drosophila). Wdr89-/- mice have neuroanatomical defects (PMID 29078390).</td></tr>
<tr><td>UPF2</td><td>E1033D</td><td>8</td><td>COAD:2, STAD:3, UCEC:3</td><td></td><td>$1.83 \times 10^{-2}$</td></tr>
<tr><td colspan="6">UPF2, regulator of nonsense transcripts 2. A mRNA surveillance complex consisting of UPF1, UPF3A and UPF3B targets an mRNA for nonsense-mediated decay (NMD) when bound downstream of a termination codon (PMID 11163187). See PMIDs 28536849 and 28866327 for recent discussions of more general mRNA quality control mediated by this complex.</td></tr>
<tr><td>NPNT</td><td></td><td>3</td><td>COAD:2, UCEC:1</td><td></td><td>$1.87 \times 10^{-2}$</td></tr>
<tr><td colspan="6">Nephronectin is an ECM ligand for integrin alpha-8/beta-1. Mice lacking nephronectin display renal agenesis (17537792). Expression frequently lost in melanoma and re-expression increases cell adhesion and reduces migration (18271919). On the other hand, expression was found to be high in metastatic breast tumors and decreased expression inhibited metastasis (15671244; but also see 28842827 for more recent analysis). Implicated in osteoblast differentiation (see 28513838 for review).</td></tr>
<tr><td>CDK4</td><td>K22M</td><td>4</td><td>SKCM:4</td><td></td><td>$2.15 \times 10^{-2}$</td></tr>
<tr><td colspan="6">CDK4 and its paralog CDK6 associate with D cyclins to phosphorylate the retinoblastoma (Rb) protein and promote cell cycle progression. The CDK4 gene is frequently amplified in cancer, and its CDKN2A inhibitor is often mutated or deleted, arguing that dysregulated CDK4 activity promotes tumorigenesis. A previous study employing extensive site-directed mutagenesis implicated CDK4 K22 residue as critically important for CCND1 binding (PMID 9228064), reporting “the most dramatic effect on cyclin D1 binding was observed for CDK4-MUT3 (residues 22-25), which decreased cyclin D1 binding to 10% relative to wild type. [Furthermore], CDK4-MUT3A (K22A) bound cyclin D1 as poorly as the triple mutant (15%). Thus, Lys-22 appears to play a greater role in cyclin D1 binding.”</td></tr>
<tr><td>SYT5</td><td>A309T</td><td>4</td><td>GBM:1, KIRP:1, UCEC:2</td><td>POLE:1, Misc:1</td><td>$2.21 \times 10^{-2}$</td></tr>
<tr><td colspan="6">Synaptotagmin 5, member of a mammalian gene family that includes 17 synaptotagmin genes (SYT1-17), and five synaptotagmin-like genes (SYTL1-5). Synaptotagmins contain a transmembrane segment and two C2 domains, the first of which binds phospholipids in a Ca++-dependent manner, and the 2nd of which has been implicated in Ca++-independent binding to the AP2 adaptor. The A309T SYT5 mutation maps to the 2nd C2 domain. Among synaptotagmins, the most extensively studied SYT1 protein is an abundant synaptic vesicle protein that functions as a calcium sensor controlling the exocytosis of neurotransmitters. Initial report indicated that SYT1, SYT2, SYT3 and SYT5 are almost exclusively expressed in neural tissues (PMID 7791877). However, SYT5 mRNA is also expressed elsewhere and has been implicated in Ca++-dependent insulin exocytosis (11309201, 15190121), as well as in macrophage phagocytosis (18832684).</td></tr>
<tr><td>PRSS48</td><td>L62W</td><td>3</td><td>CESC:3</td><td>Misc:1</td><td>$2.22 \times 10^{-2}$</td></tr>
<tr><td colspan="6">Serine protease 48, no publications. Three very low frequency alleles ($\sim 10^{-4}$) listed in ExAC. Orangutan and several other primates have a W at position 62. Based on this, the L62W mutation looks fairly benign.</td></tr>
</table>

**Table S3:** Annotations for every non-known cancer gene significant by LNP model.

<table>
<tr><th>Gene</th><th>Prot. Ch.</th><th>N. Mut.</th><th>Tumor types</th><th>Signatures</th><th>LNP q-value</th></tr>
<tr><td>BCLAF1</td><td>V347L</td><td>4</td><td>GBM:1, STAD:1, UCEC:2</td><td>POLE:3, POLE_MSI:1</td><td>$2.22 \times 10^{-2}$</td></tr>
<tr><td colspan="6">The majority of this gene is not unique on the 75mer level; this event occurs within one of the few uniquely mappable regions. As such, this event is likely a false positive.</td></tr>
<tr><td>PCDHGA3</td><td>L714H</td><td>3</td><td>LUAD:1, STAD:1, UCEC:1</td><td>smoking:1</td><td>$2.24 \times 10^{-2}$</td></tr>
<tr><td colspan="6">The entire second half of this gene (AA 500 onwards) has minimal uniqueness on the 75mer level; furthermore, the C-terminus is completely devoid of high-quality coverage. As this event occurs within this non-unique region, it is likely a false positive.</td></tr>
<tr><td>ZNF708</td><td>K353E</td><td>6</td><td>CESC:2, LUAD:3, SKCM:1</td><td>smoking:2, Misc:2</td><td>$2.46 \times 10^{-2}$</td></tr>
<tr><td colspan="6">Encoded protein contains an N-terminal KRAB box and 14 C2H2 zinc fingers (one of at least 358 KRAB/C2H2 genes in the human genome). Putative transcriptional repressor suggested by presence of KRAB box (see for example PMID 8065901). K417E mutation maps to short interval between Zn fingers 9 and 10.</td></tr>
<tr><td>RRAS2</td><td>Q72L</td><td>11</td><td>HNSC:1, LUAD:2, LUSC:1, PRAD:1, TGCT:1, UCEC:4, UCS:1</td><td>smoking:3, Misc:7</td><td>$2.58 \times 10^{-2}$</td></tr>
<tr><td colspan="6">RRAS2 shares the majority of its amino acid sequence with known Ras subfamily oncogenes HRAS, KRAS, and NRAS. Q72L mutation is paralogous to the well-known Q61 hotspot in H/K/NRAS, which abrogates GTP hydrolysis, leaving the protein constitutively active. The same constitutive activation in RRAS2 has been experimentally confirmed both in mechanism and oncogenic potency (PMID 8196649), suggesting that RRAS2 oncogenic activity is analogous to that of the more well-known Ras subfamily members.</td></tr>
<tr><td>MUC20</td><td>A8T</td><td>4</td><td>BRCA:2, CESC:1, KIRP:1</td><td>Misc:3</td><td>$2.67 \times 10^{-2}$</td></tr>
<tr><td colspan="6">The majority of this gene is not unique on the 75mer level or is insufficiently covered with high-quality reads to make mutation calls. As such, this event is likely a false positive.</td></tr>
<tr><td>CGB7</td><td></td><td>4</td><td>CESC:4</td><td></td><td>$2.79 \times 10^{-2}$</td></tr>
<tr><td colspan="6">The glycoprotein hormone family includes the pituitary hormones luteinizing hormone (LH), follicle-stimulating hormone (FSH), thyroid-stimulating hormone (TSH) and the placental hormone chorionic gonadotropin (CG). Each consist of a noncovalent dimer of a common alpha subunit and a hormone-specific beta subunit. There are 6 genes encoding CG beta subunits (CGB1-3, 5, 7 and 8) that encode very similar proteins.</td></tr>
<tr><td>RUSC1</td><td>C102S</td><td>3</td><td>COAD:1, STAD:2</td><td>Misc:3</td><td>$2.80 \times 10^{-2}$</td></tr>
<tr><td colspan="6">RUN and SH3 domain-containing protein 1, also known as NESCA. Translocates to nuclear envelope and has a role in neurotrophin-induced neurite outgrowth (15024033), perhaps by binding NEMO and TRAF6 and controlling NFkappaB signaling (19365808). Among the most believable evidence, a recent paper implicated human RUSC1 and its paralog RUSC2 as inhibitors of hedgehog signaling, based on their ability to bind and regulate SUFU/GLI complexes (most analysis done with RUSC2, but analysis in Xenopus also implicated xRUSC1 in hedgehog signaling (27633991). Mutated residue C102 is conserved in mammals but doesn't map to any obvious structural domain.</td></tr>
<tr><td>SETD6</td><td>C29F</td><td>3</td><td>KIRP:1, LIHC:1, UCEC:1</td><td>POLE:1, Misc:1</td><td>$2.80 \times 10^{-2}$</td></tr>
</table>

**Table S3:** Annotations for every non-known cancer gene significant by LNP model.

| Gene | Prot. Ch. | N. Mut. | Tumor types | Signatures | LNP q-value |
|---|---|---|---|---|---|
| Lysine methyltransferase SETD6. High throughput proteomic screens identified 190 potential substrates (22024134). SETD6 has been implicated as a regulator of NFkappaB signaling (by lysine methylation of RelA, 21131967), as a regulator of the expression of estrogen responsive genes by associating with the estrogen receptor alpha, HDAC1, metastasis protein MTA2, and transcriptional co-activator TRRAP (24751716). SETD6 also monomethylates histone variant H2AZ on lysines K4 and K7; H2AZ is upregulated in mESC and depletion of Setd6 in mESC caused differentiation, impaired self-renewal, and reduced clonogenicity (23324626). Among other potential functions, SETD6 may negative regulate the oxidative stress response (26780326), or activate Wnt/beta-catenin signaling by monomethylating PAK4 (26841865). A SETD6 dominant negative truncating mutation has been reported in familial colorectal cancer type X (28973356). Budding and fission yeast Setd6 orthologs monomethylate ribosomal protein Rpl42 and regulate ribosomal function (20444689). Mammalian Rpl42 ortholog RPL36A is among the ∼190 potential SETD6 targets. | | | | | |
| *INTS4* | S460A | 7 | COAD:2, KIRP:1, LIHC:1, LUSC:1, TGCT:2 | Misc:1 | $2.83 \times 10^{-2}$ |
| Integrator complex subunit 4, essential gene (core essentialome) widely conserved in metazoans. The integrator complex consists of 14 subunits and was initially implicated in the 3' processing of small nuclear RNA (snRNA) precursors. More recently this complex has also been shown to function in RNAPII pause-release and elongation and in eRNA (enhancer RNA) transcription. A separate SOSS complex that includes INTS3 and INTS6 (but no other INT subunits) plays a role in the DNA damage response. For a recent review see PMID 27427483. Mutation maps to penultimate HEAT repeat in an array of ∼10 HEAT repeats, which may be an interface for protein interactions. | | | | | |
| *GIGYF2* | E1116V | 3 | PRAD:1, STAD:1, UCEC:1 | Misc:1 | $2.84 \times 10^{-2}$ |
| GIGYF2, or GRB10-interacting GYF protein 2, was identified as a two-hybrid interactor with Grb10 and implicated in IGF-1 signaling (12771153, 19744960). GIGYF2 maps at PARK11 locus but missense mutations in familial Parkinson disease (18358451) are non-pathogenic polymorphisms (18923002, 19279319). Subunit of a mRNA CAP-binding EIF4E2-GIGYF2 translational repressor complex that is essential for mammalian development (22751931, 28698298). | | | | | |
| *PHF2* | T493A | 5 | COAD:1, ESCA:1, STAD:2, UCEC:1 | Misc:2 | $2.89 \times 10^{-2}$ |
| PHD finger protein 2, also known as lysine-specific demethylase PHF2, centromere protein 35, or jumonji C domain-containing histone demethylase 1E. Member of the KDM7 family (with PHF8 and KDM7A). C-terminus (AA 820-1097) of PHF2 associates with p53. PHF2 has an essential role in the p53 signaling pathway and may function as a tumor suppressor (25043306). PHF2 localizes to nucleolus and suppresses rRNA transcription by inhibiting the binding of the PHF8 transcriptional activator to rRNA promoters and by recruiting SUV39H1 (25204660). PHF2 also promotes bone formation by demethylating Runx2 (25257467) and is involved in mesenchymal-to-epithelial transition and loss of tumor-initiating activity (26941323). Frameshift mutations in MSI-H gastric and colorectal tumors (27744626). | | | | | |
| *PTP4A3* | A111D | 4 | BRCA:3, UCEC:1 | | $2.98 \times 10^{-2}$ |
| Protein tyrosine phosphatase type IVA. Small gene family with 3 members (PTP4A1-3) encode C-terminally prenylated proteins that undergo prenylation-dependent association with plasma membrane (10747914). Residue A111 is highly conserved in phosphatases and A111D mutant may lack activity, which would go against the firmly entrenched idea that PTP4A3 has positive roles in tumorigenesis and a therapeutic target (see 18224294,21053359, 26921331 for reviews). As argued in some of the 256 current papers dealing with PTP4A3/PRL-3, PTP4A3 is on small amplicon in metastasized colorectal cancers (11598267) and ectopic expression promotes migration and metastasis (12782572,15467431), potentially by regulating Rho GTPase signaling (16540666). Also implicated in promoting tumor angiogenesis (17018620). Murine PTP4A3 is a p53 target gene and its ability to induce cell cycle arrest depends on cell context and associated with inhibition of the PI3K-Akt pathway (18471976). Knockout reduces tumorigenesis (23555575 24950307). | | | | | |
| *MYC* | S146L | 8 | CESC:1, COAD:2, HNSC:3, LUAD:1, SKCM:1 | UV:1, APOBEC:4, Misc:1 | $3.00 \times 10^{-2}$ |

**Table S3:** Annotations for every non-known cancer gene significant by LNP model.

| **Gene** | **Prot. Ch.** | **N. Mut.** | **Tumor types** | **Signatures** | **LNP q-value** |
|---|---|---|---|---|---|
| *CRIP3* | K136M | 3 | LIHC:3 | Misc:1 | $3.03 \times 10^{-2}$ |
| Cysteine-rich protein 3, also known as thymus LIM protein or TLP, 204 amino acid protein harboring two LIM domains. K136M position maps to 2nd LIM domain and may affect protein interaction(s). Knockout results in reduced cellularity of murine thymus (11713292). | | | | | |
| *HIST1H4E* | R93T | 4 | CESC:1, ESCA:1, HNSC:1, LUSC:1 | APOBEC:4 | $3.11 \times 10^{-2}$ |
| The 14 members of the human HIST1H4 family all encode the same 103 residue core histone H4 protein. The COSMIC database lists two cancer samples harboring the HIST1H4E R93T mutation. A germline mutation affecting histone H4K91 was found in three individuals with a syndrome of growth delay, microcephaly and intellectual disability. H4K91 was previously implicated as important for DNA repair and genome stability and the authors suggested that ubiquitination of H4K91 affects genome stability during embryonic development (PMID 28920961). | | | | | |
| *TATDN1* | G171G | 4 | COAD:1, LIHC:2, TGCT:1 | Misc:2 | $3.22 \times 10^{-2}$ |
| Putative deoxyribonuclease TATDN1, member of the metallo-dependent hydrolase superfamily, (not very closely) related to TATDN2 and TATDN3. Yeast ortholog Tad-D/YBL055C and C. elegans ortholog crn-2 have roles in apoptosis (15657035). Residue 171 maps near catalytically important His-174. Knockdown of zebrafish TATDN1 ortholog, which is predominantly expressed in eye cells during embryonic development and which can de-concatenate DNA, results in abnormal cell cycle progression, polyploidy and aberrant chromatin structures (23187801). | | | | | |
| *KNSTRN* | R32R; V12G | 6; 3 | HNSC:1, SKCM:5; LUAD:1, STAD:2 | UV:6; Eso:2 | $3.51 \times 10^{-2}$; $9.26 \times 10^{-2}$ |
| Kinetochore localized astrin/SPAG5 binding protein. Missense mutations in 19% of cutaneous squamous cell carcinomas. Cancer associated mutations, most notably p.S24F, disrupt chromatid cohesion (PMID 25194279). Several other missense mutations documented in this study, but no mention of V12G mutation. | | | | | |
| *ZC3H4* | E779Q | 4 | COAD:1, STAD:3 | Misc:4 | $3.65 \times 10^{-2}$ |
| *CRNKL1* | S128F | 7 | SKCM:7 | UV:7 | $3.70 \times 10^{-2}$ |
| Crooked neck pre-mRNA splicing factor-like 1, widely conserved from yeasts to man, although many mammalian and other orthologs appear to start near position 160 of the 848 amino acid human protein (i.e. lack the N-terminal segment that includes the S128F mutation). Only some primate species include the extended N-terminal sequence of the human protein, which is an ortholog of Drosophila crooked neck; Budding yeast ortholog Clf1p is a member of the NineTeen Complex (NTC), which contains Prp19p and stabilizes U6 snRNA in catalytic forms of the spliceosome containing U2, U5, and U6 snRNAs. PMID 23774526 reported three genes, STAT5B, CRNKL1, and NEBL, with mutational hot spots at a single base in 3 of 12 basal cell carcinomas sequenced. The CRNKL1 mutation is the same S128F mutant. | | | | | |
| *HMGN5* | A60A | 6 | BRCA:3, ESCA:2, UCEC:1 | Misc:2 | $4.04 \times 10^{-2}$ |
| High mobility group nucleosome binding domain 5. Members of the HMGN family bind to nucleosomes and affect chromatin structure/function, including transcription and DNA repair. HMGN5 is enriched in euchromatin and decompacts chromatin by interacting with and counteracting compaction mediated by linker histone H1 (21518955). Also affects nuclear sturdiness (25609380). HMGN5 overexpression altered the expression of >2000 mouse genes, whereas siRNA silencing affected 308 genes (19748358, see also 23620591). Highly expressed and has positive role in several cancer types (21695596, 21373965,22504871, 25315189). | | | | | |
| *PDS5B* | R394* | 6 | STAD:2, UCEC:4 | MSI:1, Misc:1 | $4.09 \times 10^{-2}$ |

Table S3: Annotations for every non-known cancer gene significant by LNP model.

| Gene | Prot. Ch. | N. Mut. | Tumor types | Signatures | LNP q-value |
|---|---|---|---|---|---|
| PDS5B (also known as APRIN) is a sister chromatid cohesion protein crucial for the faithful segregation of duplicated chromosomes in lower organisms. Mammals have two paralogs, PDS5A and PDS5B with apparently partially redundant functions (PMIDs 15855230, 19412548, 24141881). A potential role as a tumor suppressor has been suggested: Loss of a cohesin-linked suppressor APRIN disrupts stem cell programs in embryonal carcinoma: an emerging cohesin role in tumor suppression (PMID:20383194). PDS5B physically interacts with BRCA2 and is required for maintaining genome integrity (PMID 22293751). | | | | | |
| *ONECUT2* | H184Q | 5 | COAD:4, STAD:1 | MSI:2 | $4.17 \times 10^{-2}$ |
| Functions of ONECUT family members include partially redundant roles in tissue differentiation (17400205), hepatoblast migration (17936262), and retinal development (25228773). | | | | | |
| *DTD2* | M1V | 5 | ACC:1, BRCA:1, GBM:1, KIRP:1, LUSC:1 | Misc:1 | $4.50 \times 10^{-2}$ |
| Several l-aminoacyl-tRNA synthetases can transfer a d-amino acid onto their cognate tRNA(s). This harmful reaction is counteracted by the enzyme d-aminoacyl-tRNA deacylase (see PMID 10918062). DTD2 (conserved in vertebrates) is distantly related to the more widely distributed DTD1 and has been implicated as a putative Dtyr-deacetylase by virtue of its sequence similarity. | | | | | |
| *SLC29A4* | E305G | 4 | BLCA:1, GBM:1, HNSC:1, PAAD:1 | Misc:1 | $4.88 \times 10^{-2}$ |
| Equilibrative nucleoside transporter 4, involved in transport of monoamines, such as serotonin (12838422, 15448143, 17046718). Also transports metformin (17600084); KO mice have impaired monoamine uptake (23255610). | | | | | |
| *TUBGCP2* | A615S | 5 | BRCA:4, MESO:1 | Misc:3 | $4.91 \times 10^{-2}$ |
| TUBGCP2, or gamma-tubulin complex component 2 (GCP2), is a subunit of multi-subunit $\gamma$-tubulin ring complexes ($\gamma$TURCs) that nucleate tubulin polymerization and are key components of microtubule organizing centers (see 20861304 for overview). TUBGCP2 is upregulated by Taxol. Residue 615 of isoform 2 is an alanine. The same is not true for the longer isoform 1. Unclear how A615S mutation might affect TUBGCP2 function, but MTOC problems may affect cell division or other cell functions. | | | | | |
| *ING1* | R339* | 12 | CESC:1, COAD:1, ESCA:1, LGG:1, READ:1, STAD:2, UCEC:5 | CpG:5 | $5.25 \times 10^{-2}$ |
| ING1 dimerizes with WT p53 (determined via coimmunoprecipitation, PMID 9440695). Overexpression in lung cancer cell line results in apoptosis (PMID 21286670), with apoptosis rate significantly higher in A549 cell line (with WT p53) than in SK-MES-1 (with mutant p53). R339 of 422 amino acid ING1 isoform 4 corresponds to R196 of the canonical 279 residue ING1 isoform 1. COSMIC uses isoform 1 coordinates and lists 16 samples with a R196* nonsense mutation, that truncate the protein before its C-terminal H3K4Me3-binding PHD domain (see PMID 23412501 for a review). In addition, P33ING1 has been implicated as a component of chromatin remodeling complexes that regulate histone acetylation (PMID 12015309). | | | | | |
| *HSPA1L* | K389N | 3 | CESC:3 | | $5.72 \times 10^{-2}$ |
| *ZNF658* | I255T | 5 | COAD:2, READ:1, UCEC:2 | Misc:1 | $6.75 \times 10^{-2}$ |
| The majority of this gene is not unique on the 75mer level; this event occurs within the only ∼100 bp window that is uniquely mappable. As such, this event is likely a false positive. | | | | | |
| *CDC25A* | R446* | 5 | COAD:1, STAD:1, UCEC:3 | | $6.81 \times 10^{-2}$ |

**Table S3:** Annotations for every non-known cancer gene significant by LNP model.

| Gene | Prot. Ch. | N. Mut. | Tumor types | Signatures | LNP q-value |
|---|---|---|---|---|---|
| Cell division cycle 25A encodes a protein dual-specificity phosphatase that positively regulates the G1/S and G2/M cell cycle transitions by removing inhibitory phosphates from the ATP binding site of cyclin-dependent kinases (1828290,1836978,18073536). Implicated as potential oncogene (7667636). Inactivation of CDC25A by ubiquitin-mediated degradation critical for DNA damage-induced checkpoint control. Cdc25A null mice die at embryonic day 5-7. Cdc25A(+/-) MEFs are resistant to transformation (17638870). Thus, heterozygous loss of CDC52A function resulting from the R446* mutation, which truncates CDC25A's C-terminal rhodanese-like phosphatase catalytic domain just downstream of the highly conserved CX5R motif (9604936), would be predicted to be deleterious for (cancer) cell viability. However, perhaps truncating CDC25A at R446 does not result in a complete loss-of-function phenotype. Among functions that involve the extreme C-terminus of CDC25A are cyclin-B and 14-3-3 binding and PLK3-mediated phospho-regulation (15640846). | | | | | |
| *GNAZ* | L131I | 4 | BRCA:1, COAD:1, SARC:1, UCEC:1 | POLE_MSI:1, Misc:1 | $6.83 \times 10^{-2}$ |
| Heterotrimeric G protein subunit Gz-alpha, member of the Gi subfamily of G-alpha proteins, is predominantly expressed in neuronal cells and platelets. PKC-mediated phosphorylation blocks interaction with $\beta\gamma$subunits (7559455). G(alpha)z couples neurotransmitter receptors to N-type Ca2+ channels when transiently overexpressed in rat sympathetic neurons (9856474). Murine knockout results in abnormal platelet activation (but see 11307826) and altered responses to psychoactive drugs (10954748). Gz-alpha also expressed and functionally important in pancreatic beta cells (16157560). Four GNAZ somatic missense mutations identified in 80 melanoma samples (N43S, S80L, P98S, G451A; PMID 20424519). | | | | | |
| *PPP2R2B* | V395G | 4 | BLCA:1, COAD:1, STAD:1, UCEC:1 | Misc:1 | $6.90 \times 10^{-2}$ |
| Protein phosphatase 2 55 kDa regulatory subunit Bbeta largely consists of WD40 repeats and is a member of a family of 4 highly related genes (PPP2R2A-D). Valine-395 of 432 amino acid isoform D corresponds to Valine-512 of longest (549 residue) isoform e and is conserved in all four family members. Regulatory subunit of protein phosphatase 2A that interacts with three subunits of the Ska complex (19387489), a key component of the kinetochore-microtubule interface. | | | | | |
| *MORC2* | Q99L | 4 | SKCM:4 | Misc:1 | $8.71 \times 10^{-2}$ |
| Microrchidia family CW-type zinc-finger 2, mutations in MORC2 cause axonal Charcot-Marie-Tooth disease (26497905, 26659848). | | | | | |
| *OR5D16* | R6G | 4 | STAD:4 | Eso:3 | $9.61 \times 10^{-2}$ |
| Olfactory receptor, a likely false positive. | | | | | |
| *ZNF276* | G163S | 4 | BLCA:2, HNSC:1, SKCM:1 | UV:1, Misc:1 | $9.84 \times 10^{-2}$ |
| Gene encoding a protein with N-terminal zf-AD (putative dimerization) domain and C-terminal C2H2 zinc fingers. Gene overlaps in tail-tail manner with final four exons of FANCA (PMID 10936049). Mutation analysis in breast cancer detected two missense mutations, E530D and R200W, the latter a common polymorphism. Nothing else is known. G163 maps just C-terminal to the zf-AD putative dimerization domain and is conserved in many species. | | | | | |
| *FMN2* | D162Y | 4 | LUAD:3, LUSC:1 | smoking:3 | $9.84 \times 10^{-2}$ |
| Formin 2/FMN2 is an actin filament nucleator that functions together with SPIRE1/2 (17923532,19605360,21705804,21730168,24586110). From PMID 27839864: "We found that FMN2 is upregulated in human melanomas and showed that disruption of FMN2 in mouse melanoma cells inhibits their extravasation and metastasis to the lung. Our results indicate a critical role for FMN2 in generating a perinuclear actin/FA system that protects the nucleus and DNA from damage to promote cell survival during confined migration and thus promote cancer metastasis." | | | | | |
